# *In silico* analyses of neuropeptide-like protein (NLP) profiles in parasitic nematodes

**DOI:** 10.1101/2021.03.03.433794

**Authors:** Fiona M. McKay, Ciaran J. McCoy, Nikki J. Marks, Aaron G. Maule, Louise E. Atkinson, Angela Mousley

**Author notes:** Corresponding author: Angela Mousley. School of Biological Sciences, Queen’s University Belfast, 19 Chlorine Gardens, Belfast, BT9 5DL, UK. Note: Supplementary data associated with this article.

## Abstract

Nematode parasite infections cause disease in humans and animals and threaten global food security by reducing productivity in livestock and crop farming. The escalation of anthelmintic resistance in economically important nematode parasites underscores the need for the identification of novel drug targets in these worms. Nematode neuropeptide signalling is an attractive system for potential chemotherapeutic exploitation, with neuropeptide G-protein coupled receptors (NP-GPCRs) representing the leading target candidates therein. In order to successfully validate NP-GPCRs as targets for parasite control it is necessary to characterise their function and importance to nematode biology. This can be aided through identifying receptor activating ligand(s) in a process known as receptor deorphanisation. Such efforts first require the identification of all neuropeptide ligands within parasites. Here we comb the genomes of nine therapeutically relevant pathogenic nematode species to comprehensively characterise the nematode parasite neuropeptide-like protein (NLP) complements, and details the discovery of several previously unreported, yet conserved, neuropeptides and neuropeptide-encoding genes. We identify the neuropeptides that are most highly conserved in all parasites examined, and characterise their physiological activity on the reproductive musculature of the parasite, *Ascaris suum*. These data suggest conserved neuropeptide functions in both free living and parasitic nematodes, and support the potential for exploitation of the neuropeptide signalling system as an anthelmintic target.

## 1. Introduction

Nematode parasites continue to threaten human health and global food security (Charlier et al., 2014; Lopes-Caitar et al., 2019; Hotez and Lo, 2020). The impact of nematode parasites is exacerbated by the limited number of available anthelmintic drugs and the escalation of anthelmintic resistance in key nematode pathogens (Kaplan and Vidyashankar, 2012; Sangster et al., 2018). The disruption of nematode neuromuscular signalling is a well-established route to parasite control (McVeigh et al., 2006a; Li and Kim, 2008; McVeigh et al., 2012). Given that neuropeptide signalling modulates nematode neuromuscular function, it represents an appealing unexploited source of novel anthelmintic targets.

Nematode neuropeptide ligands encompass the FMRF-amide like peptides (FLPs), Neuropeptide-like proteins (NLPs) and the Insulin-like peptides (INSs). Whilst INS peptides appear to signal through receptor tyrosine kinases, the majority of FLPs and NLPs are believed to act as ligands for G-protein coupled receptors (GPCRs) (Li and Kim, 2008). Within the neuropeptide system, neuropeptide GPCRs (NP-GPCRs) emerge as the most promising target candidate, with a proven history of ‘druggability’ (for review see (McVeigh et al., 2012). Ultimately, the validation of NP-GPCRs as therapeutic targets requires the interrogation of their individual functions to enable the selection of the most appropriate target candidates.

NP-GPCR deorphanisation, i.e. the identification of the activating endogenous ligand(s) of the receptor, is an important step in NP-GPCR functional characterisation. While heterologous approaches to NP-GPCR deorphanization in *Caenorhabditis elegans* have been successful, only a limited of *Caenorhabditis elegans* NP-GPCRs have been functionally deorphanized to date, and only two reports of successful NP-GPCR deorphanisation through heterologous expression and knockdown (RNA interference) phenocopy have been described in parasites (Atkinson et al., 2013; Anderson et al., 2014). To aid deorphanisation efforts in therapeutically/economically relevant nematode parasites, we first require a complete profile of all potential interacting neuropeptide ligands and receptors. Recent studies have highlighted comprehensive analyses of FLP and NP-GPCR profiles in parasitic nematodes (McCoy et al., 2014), however our current understanding of parasitic nematode NLP complements is comparatively outdated (McVeigh et al., 2008).

Here, we utilise an established neuropeptide BLAST (Basic Local Alignment Search Tool) approach to provide a comprehensive analyses of NLP-gene profiles in nine key therapeutically/economically important nematode parasite species, and examine the muscle-based responses of those NLPs emerging as the most highly conserved. These data advance understanding of nematode neuropeptide signalling and are critical to future deorphanisation efforts in parasitic nematodes that drive novel parasite control strategies.

## 2. Material and Methods

### Species selection

*Caenorhabditis elegans nlp*-gene sequelogues were identified in the genomic datasets of 9 parasitic nematode species (Table S1). Species were selected based on the availability of complimentary datasets detailing the neuropeptide GPCR and *flp*-gene complements of these nematodes (McCoy et al., 2014), and their importance as either animal (including human) or plant parasites. These species also encompass a range of nematode clades and parasite lifestyles.

### BLAST searches

A reciprocal BLAST method was used to identify *nlp*-gene sequelogues using the ‘prepropeptide search string’ approach (McCoy et al., 2014). Protein sequences for NLPs identified previously in *C. elegans* (Nathoo et al., 2001; Husson et al., 2005; Husson et al., 2007; McVeigh et al., 2008; Yamada et al., 2010; Husson et al., 2014; Howe et al., 2017; Van Bael et al., 2018b); Table S2) were obtained from the WormBase database (https://www.wormbase.org; <u>WS276</u>) and used as queries in tBLASTn and BLASTp searches of the genomic datasets. Translated nucleotide (tBLASTn) searches were carried out on the WormBase ParaSite server (https://parasite.wormbase.org/index.html; Howe et al., 2017), protein (BLASTp) searches were carried out on the WormBase ParaSite Server or via a locally ran Windows command line based NCBI-BLAST. All BLAST searches were carried out between April 2017 and August 2018. Where there were multiple genomes available for a species, both were mined (Table S1). BLAST searches were also carried out in four additional *Caenorhabditis* species [*C. brenneri, C. briggsae, C. remanei* and *C. japonica*; as in (McCoy et al., 2014)]. The so-called ‘antimicrobial’ *nlp*-genes (*nlp*-24-34), distinguished by their glycine-rich sequences were not included in this study due to their unsuitability for the BLAST search approach and their putative antimicrobial roles (McVeigh et al., 2008; Pujol et al., 2008; Dierking et al., 2016). Where *nlp*-genes encoded multiple isoforms, only the longest were used as query sequences unless isoforms were vastly different in sequence. Expect values were set to ≥1000 to avoid false negative hits. All returned sequences were examined by eye for known neuropeptide motifs flanked by putative mono/dibasic cleavage sites, signal peptide cleavage sites or C-terminal sequence ends to eliminate false positive hits (McVeigh et al., 2008; McCoy et al., 2014).

### Post-BLAST analysis

Neuropeptide-like protein encoding gene sequelogues were assigned primarily based on sequence similarity in the predicted peptide region in *C. elegans* (Li and Kim, 2008; Van Bael et al., 2018b). Sequences were aligned using Vector NTI Advance 11.5 AlignX® multiple sequence alignment tool (Lu and Moriyama, 2004). Signal peptide predictions were made using SignalP4.1 (Nielsen, 2017), however lack of a predicted signal peptide did not necessarily exclude sequences from designation as an NLP.

### CLANS analysis

The Clustered Analysis of Sequences (CLANS) algorithm was used to perform all-against-all BLASTp comparisons between the identified nematode RPamide and Allatostatin-C like prepropeptide sequences, and generate a 3D similarity matrix derived from these individual searches. Prepropeptide sequences were uploaded to the CLANS website (https://toolkit.tuebingen.mpg.de/#/tools/clans) where the most appropriate E-value limit was determined via trial and error for each specific input to facilitate sufficient cluster separation. All other parameters were set as default. The CLANS file output was examined and coloured after 10,000 clustering rounds using the Java-based desktop software.

### Ascaris suum ovijector physiology assay

Selected NLP peptides predicted in *A. suum* were tested for activity on the ovijector (reproductive muscle) of *A. suum* using an established physiology assay (Fellowes et al., 1998; Fellowes et al., 2000; Moffett et al., 2003). Adult female *A. suum* (>20cm) were collected from the intestines of freshly slaughtered pigs at a local abattoir (Cookstown, Northern Ireland) and transported back to the laboratory in mammalian saline (0.9% NaCl, 37°C). Worms were maintained in *Ascaris* Ringers Solution [ARS; 13.14mM NaCl, 9.47mM CaCl_2_, 7.83mM MgCl_2_, 12.09mM C_4_H_11_NO_3_(Tris), 99.96Mm NaC_2_H_3_O_2_, 19.64mM KCl, pH 7.8] at 37 °C and 5 % CO_2_ for up to four days with media changes twice daily. The ovijector was dissected from healthy, turgid worms and transferred to Hank’s balanced salt solution (HBSS; Life technologies) at 37°C in a 4ml recording chamber, where the tissue was attached between a flexible and inflexible pipette. Tissue activity was amplified and recorded via a photo-optic transducer system (Fetterer et al., 1977; Marks et al., 1996). Ovijectors which displayed regular, spontaneous contractility were equilibrated for at least five minutes before peptide addition. Inactive or erratically active ovijectors were discarded. Peptides were synthesised (Genosphere Biotechnologies Inc.) and 10mM stock solutions prepared in double distilled (dd)H_2_O, Dimethyl sulfoxide (DMSO), or Dimethyl formaldehyde (DMF) and stored in aliquots at −20°C until use. Addition of 4μl ddH_2_O, DMSO or DMF had no effect on ovijector activity. Peptide was added to the 4ml water bath such that addition of 4μl gave a final peptide concentration of 10μM. Peptide effects were recorded for 10 minutes before media was replaced with fresh HBSS. The ovijector was subsequently recorded for a further 10 minutes to observe recovery. Contraction amplitude and frequency were measured 2 minutes prior to time 0 (peptide addition), as well as 2, 5, 10 and 20 minutes post-addition. In some cases, it was necessary to analyse additional time points (e.g. 50 seconds post addition) to capture the transient effects. Change in tension was also measured at time 0, 2, 5, 10 and 20 minutes post-addition, where time 0=0mg/mm. Note that muscle relaxation caused an increase in circular muscle tension (+mg/mm; shortening of the tissue) whereas circular muscle contraction caused a decrease in tension (-mg/mm; lengthening of tissue). Data were statistically analysed (Graphpad Prism 8) with repeated measures analysis of variance (ANOVA) followed by Dunnetts post-test to compare each time point to the time 0.

## 3. Results and discussion

### 3.1 Parasitic nematodes possess a reduced complement of Caenorhabditis elegans NLP-encoding genes

326 *nlp*-gene sequelogues were identified in nine nematode parasite species of key importance (Table 1; Fig. S1; Table S3), selected based on the quality and availability of genome data and their importance to human, animal and plant health (Table S1). These included 73 previously reported NLP-encoding genes identified through *in silico* studies (Nathoo et al., 2001; McVeigh et al., 2008; Jarecki et al., 2011; Koziol et al., 2016; Warnock et al., 2017; Van Bael et al., 2018b), and 253 novel sequences identified here. These data provide a comprehensive, phylum-spanning insight into *nlp* conservation in nematode parasites, enabling comparison between distinct clades and lifestyles. As noted with parasite FLP and NP-GPCR datasets (McCoy et al., 2014; Unpublised observations) the parasitic nematode species investigated here displayed reduced complements of the *C. elegans* NLP (*Ce*-NLP) profile (Table 1). The human hookworm *Necator americanus* exhibited the largest share (67%) of *C. elegans nlps* (*Ce-nlps*), this is unsurprising considering they are both members clade 9 (Holterman et al., 2006). In contrast, the clade 2 species *Trichinella spiralis* and *Trichuris muris* displayed a significantly reduced *nlp* complement of 15% and 14% of the *C. elegans nlp* profile respectively. This is likely a result of both specific gene loss in the clade 2 species, as well as *nlp* gene duplication events within the lineages that led to the extant nematodes that now comprise the ‘crown’ clades (Holterman et al., 2006) that exhibit broader *nlp* complements.

**Table 1.**
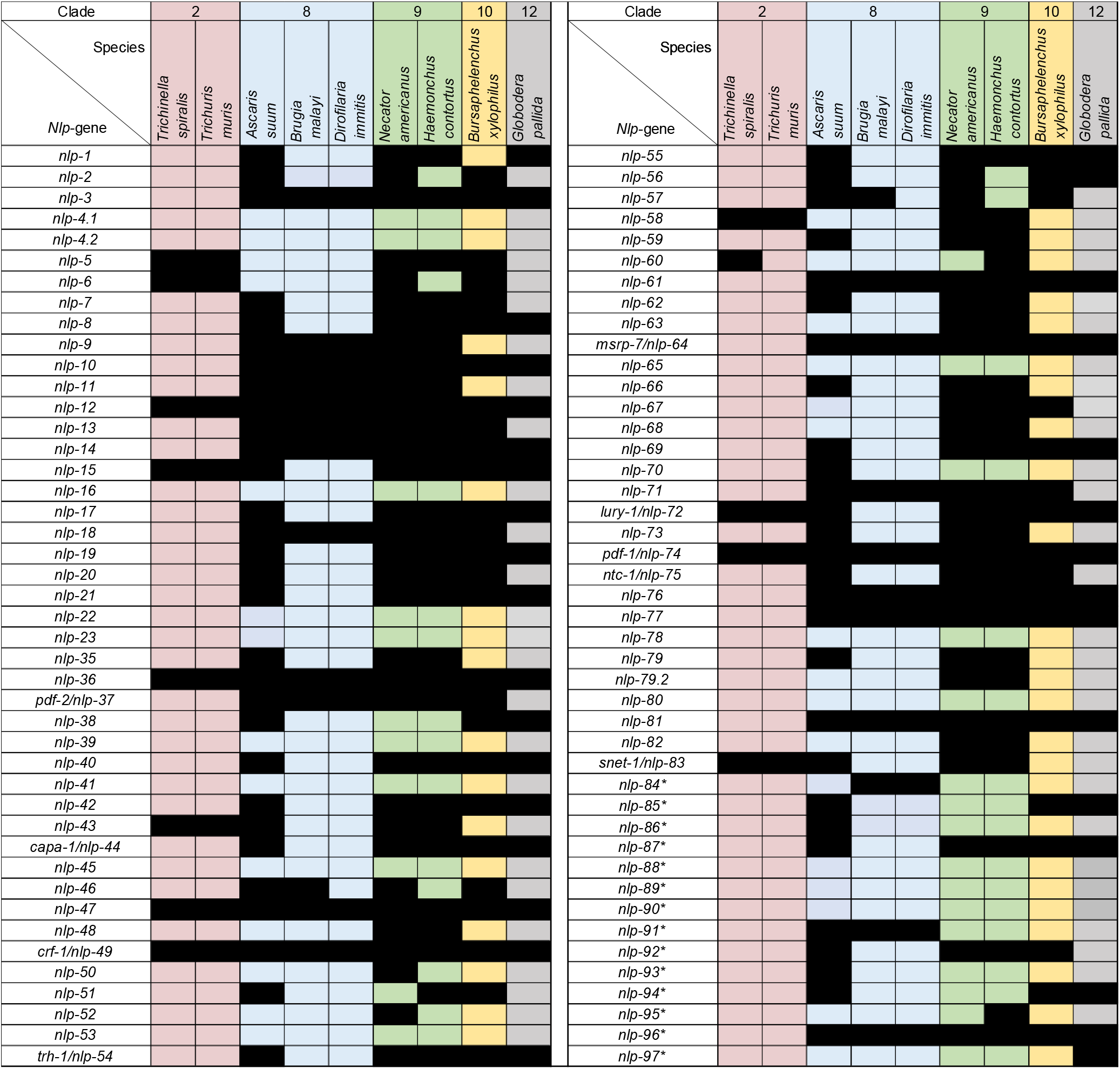
*nlp*-gene sequelogues in nine species of nematode parasites. A black box indicates the presence of a sequelogue identified via BLAST search. An Asterix indicates a novel gene identified here.

### 3.2 *Parasite* nlp *profiles are broadly conserved within nematode clades*

Within the majority of nematode clades (Holterman et al., 2006) *nlp* profiles were relatively similar, however more variation in *nlp* profile patterns was noted between clades, mirroring trends observed for *flps* and NP-GPCRs (McCoy et al., 2014).

The clade 2 nematodes (*T. spiralis* and *T. muris*) possess a near-identical *nlp* profile, however notable differences in *nlp* profiles were observed within clade 8 species. Indeed, although the clade 8 filarids displayed near-identical *nlp* profiles, they were different from that observed for *A. suum*, also within clade 8. This is not surprising given that the filarid species are more closely related to each other than to *A. suum*, and likely reflects key differences in the lifestyles of the filarial species (insect vectored and mammalian host circulatory system dwelling parasites) relative to *A. suum* (gastrointestinal nematode). Although the plant parasitic nematode species in this study (*Bursaphelenchus xylophilus* and *Globodera pallida*) are each the sole representatives of their respective clades (10 and 12), they possess *nlp*-profiles which overlap. This pattern of *nlp*-profile conservation could be linked to their shared lifestyle or may reflect the profile displayed by their last common ancestor.

### 3.3 *The degree of* nlp *conservation is gene dependent*

Five *Ce-nlps* were completely conserved across all nine species interrogated in this study: *nlp*-12, -36, -47, -49 (*crf*-1) and *pdf*-1(*nlp*-74) (Table 1; Fig. S1), highlighting their likely functional importance to nematode biology/behaviour, and supporting their presence in the last common ancestor of all nematodes. By contrast, ten already annotated *Ce-nlps* were absent from all parasites examined: *nlp*-4.1 -4.2 -16, -39, -41, -45, -53, -65, -78, and *nlp*-80, and so have either been lost independently in multiple parasitic nematode lineages, or have evolved relatively recently in the specific lineage that led to the model worm. Indeed, although all other *Caenorhabditis* species examined here possess a similar NLP profile to that of *C. elegans* (Table S4), the other *Caenorhabditids* examined lacked *nlp*-39, suggesting that this gene is either highly divergent or arose following the split of *C. elegans* lineage from *C. briggsae*, *C. brenneri* and *C. remanei* (~5-30 MYA; Frezal & Felix (2015)). Similarly, the absence of *nlp*-4.1, -4.2, and -65 in *C. japonica* suggest that these genes likely arose following the separation of the *Japonica* and *Elegans* groups (~125-190 MYA; Lemos (2007), Frezal & Felix (2015)). Please note that we designate the *C. elegans* genes F59C6.6 and F59C6.18 as *nlp*-4.1 and *nlp*-4.2 respectively, as both genes appear to have been independently annotated as ‘*nlp*-4’ previously (see both WormBase (WS276) and Nathoo et al. (2001)).

### 3.4 The pattern of NLP amino acid residue conservation across key nematodes varies between nlps

The extent of the amino acid residue conservation within the NLPs encoded on each gene varied (Fig. S1). For example, in some instances primary sequence conservation was observed across the entire length of the encoded peptide (e.g. NLP-3, -43), however in other cases conservation was biased toward either the C-(e.g. NLP-17) or N-termini (e.g. NLP-9, -21) of the encoded peptides. Some NLPs (e.g. NLP-6) displayed relatively low levels of sequence conservation across the entire length of the peptide, however this did not preclude their identification using the motif-based BLAST approach employed here. These observations suggest different levels of functional constraint between peptides, and are likely indicative of the motif that is important for receptor recognition or binding.

### 3.5 Some Ce-NLPs identified by mass spectrometry are not conserved in nematode parasites challenging their functional relevance

For the most part, the conserved motifs identified through sequence alignments of putative prepropeptide nematode parasite NLP sequelogues map to the previously predicted peptide regions identified via *C. elegans in silico* and mass spectrometric analyses (Husson et al., 2005; Van Bael et al., 2018b).

However, for several NLPs (NLP-35, NLP-36, NLP-55, NLP-56, NLP-76 and NLP-79) some of the peptides identified by *C. elegans* mass spectrometry (Husson et al., 2005; Van Bael et al., 2018b) are not conserved in any of the parasite species examined in this study (Fig. S1; Table S5), and are therefore less likely to represent functionally important peptides but instead by-products of neuropeptide processing events. Moreover, sequence alignments of some parasite- and *C. elegans*-NLP prepropeptides (e.g. NLP-56, NLP-79; Fig. 1) reveals alternative highly conserved motifs, that were not previously considered as putative peptides by Van Bael et al. (20018b), but may represent functional NLPs. These data are significant in building peptides libraries for future deorphanisation efforts and underscore the value of pan-phylum *in silico* analyses for peptide prediction.

**Fig. 1.**
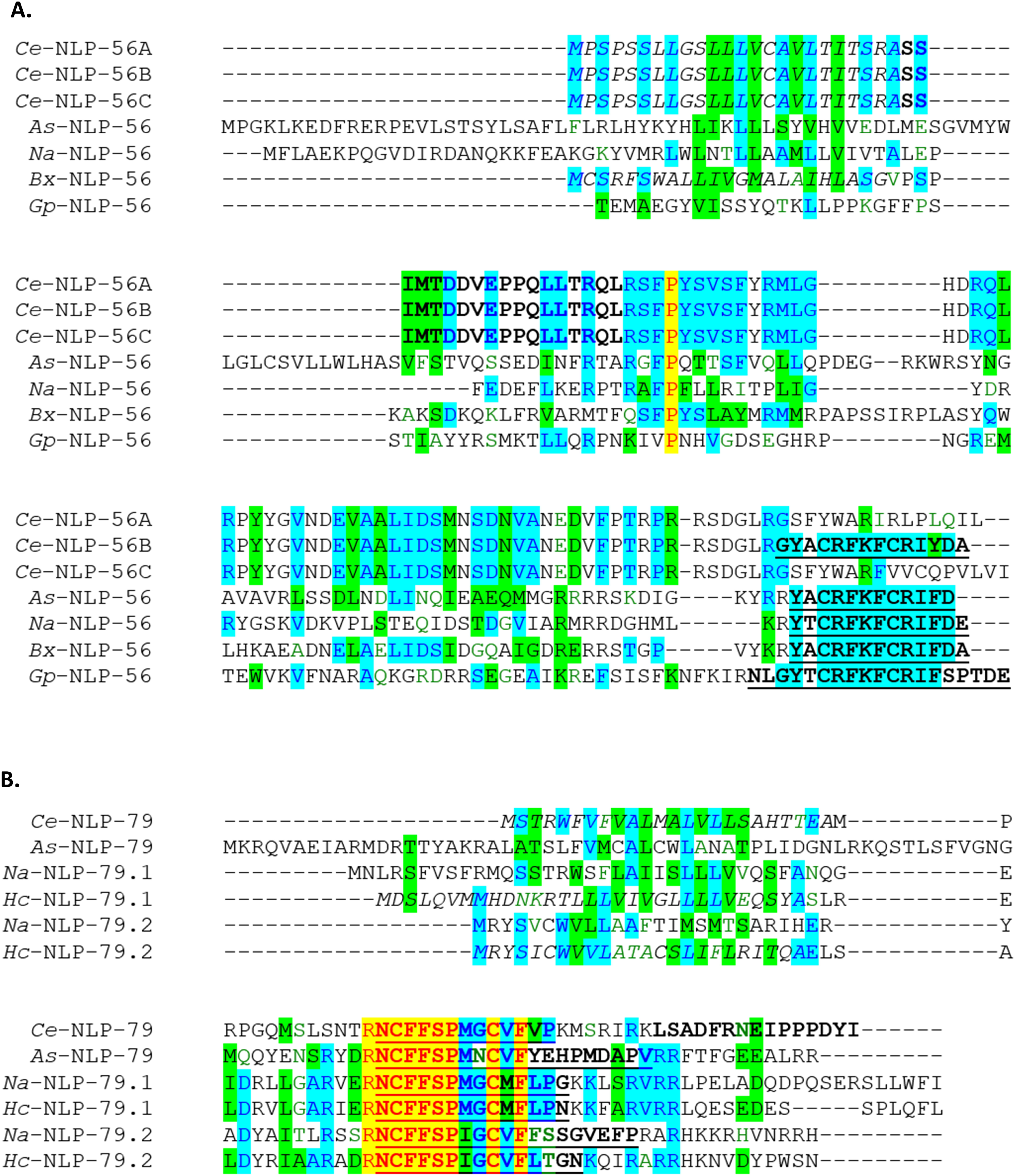
Novel predicted *Ce*-NLPs are conserved in parasitic nematodes. Multiple sequence alignments were performed using Vector NTI Advance 11.5 AlignX® (Lu and Moriyama, 2004). Italic text indicates the presence of a signal peptide as identified using Signal P 4.1 (Nielsen, 2017). Bold text indicates peptides predicted previously (see Van Bael et al., 2018b). Bold underlined text indicates novel peptides predicted here based on their conservation in other nematode species. Yellow and blue highlighted regions indicate completely and partially conserved amino acid residues respectively. Green indicates similar residues. *Ce* denotes *C. elegans*, *As* denotes *Ascaris suum*, *Na* denotes *Necator americanus*, *Hc* denotes *Haemonchus contortus*, *Bx* denotes *Bursaphelenchus xylophilius*, *Gp* denotes *Globodera pallida*

### 3.6 *Parasitic nematode* in silico *analyses reveals novel NLP-encoding genes and highlights the need for nomenclature revision*

Phylum-spanning *in silico* analyses of neuropeptide-encoding genes, using the reciprocal BLAST approach outlined here, not only drives novel peptide discovery in parasites but also provides an opportunity for identification of additional *nlps* in *C. elegans*.

This study discovered 14 novel *nlps* (Tables 1 and 2), which have been designated *nlp*-84 to *nlp*-97. For continuity with previously identified *Ce*-*nlps* (McVeigh et al., 2008; Nathoo et al., 2001; Van Bael et al., 2018b), and before naming any further novel *nlp*-genes here, we have reassigned *snet*-1 as *nlp*-83. Although not classed as an *nlp*-gene by the current version of WormBase (WS276), *snet*-1(*nlp*-83) was identified as a *C. elegans* neuropeptide gene in an olfactory plasticity study and previously considered as a member of the NLP family (Yamada et al., 2010; Hobert, 2013). The novel *nlps* discovered in this study include additional members of (i) RPamides, (ii) Allatostatins and a number of entirely novel *nlps* that had not previously been reported in *C. elegans*.

#### 3.6.1 Novel RPamide-encoding genes were identified in parasitic nematodes

The *C. elegans* RPamide family consists of the *nlp*-2, -22 and -23 gene cluster first described by Nathoo et al. (2001), plus *nlp*-46, identified by McVeigh et al. (2008). The NLP-2, -22 and 23 peptides are highly similar, making sequelogue designation difficult for BLAST hits using the *C.elegans* seeded BLAST approach. We therefore extended our BLAST searches using all of the RPamide encoding BLAST returns to query all available nematode genomes, and performed multiple sequence alignments to manually identify patterns of sequence conservation. Of the known RPamide encoding genes, *nlp*-22 and -23 sequelogues were only identified in *Caenorhabditis spp*, whereas *nlp*-2 and *nlp*-46 were conserved in a number of parasites (Table 1, Table S4).

Three novel RPamide-encoding genes were identified in parasitic nematodes: (i) A novel RPamide sequence [A(A/V/T)MISGRGFRPG], initially identified in *B. malayi* and *D. immitis,* and sharing a common motif with RPamides encoded in 14 other filarial species, was classified as a novel filarid-specific RPamide-encoding gene (*nlp*-84); (ii) A second novel RPamide peptide-encoding gene (designated here as *nlp*-85; GRW(G/Q)LRPG), identified initially in *A. suum*, *B. xylophilius* and *G. pallida* was also present in 11 additional nematode species representing both free-living and parasitic lifestyles; finally (iii) A third novel *nlp (nlp-86*; S(I/L)ALGR(F/L)(S/N)LRPG), identified initially in *A. suum*, and subsequently in seven additional nematode species appears distinct from peptides encoded on *nlp*-2 and *nlp*-85.

Both the presence of *nlp*-2, -85, and -86 in the *A. suum* genome, encoding distinct RPamide peptide motifs, and subsequent CLANS analysis provides evidence to support the novel *nlp* designations (Fig. 2 A). However, several previous studies have been unable to accurately delineate the RPamide-encoding *nlps* such that there are conflicting designations in the literature to what we describe here (McVeigh et al., 2008; Jarecki et al., 2011; Nelson et al., 2013 and Koziol et al., 2016). This underscores the value of phylum-spanning *in silico* analyses to unravel the complexity of the NLP family and associated nomenclature. In addition, our approach offers a route to interrogate neuropeptide evolution; indeed, both motif and CLANS analyses reveal that *nlp*-60 (a non-RPamide encoding gene) likely represents an RPamide family member that is closely related to *nlp*-86 but has lost all RPamide peptides (Fig. 2 A).

**Fig. 2.**
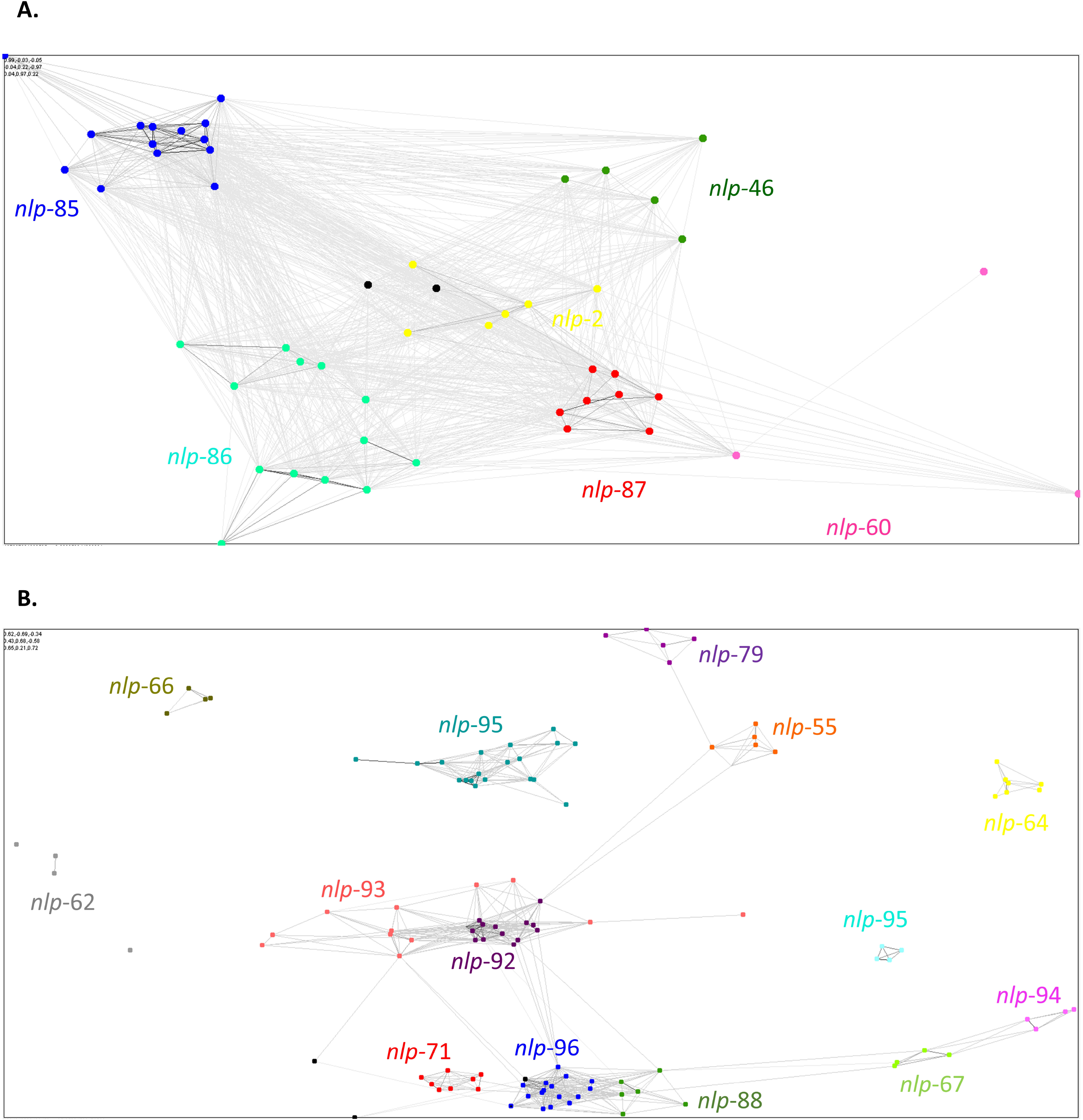
CLANS clustering supports RPamide and Allostatin-C like peptide encoding gene nomenclature. (A) Similarity matrix derived from all-against-all BLASTp comparisons between all identified nematode RPamide encoding prepropeptide sequences (E-value limit = 1). (B) Similarity matrix derived from all-against-all BLASTp comparisons between all identified nematode Allatostatin-C like prepropeptide sequences (E-value limit = 1E-5).

#### 3.6.2 Novel Allatostatin-C-like peptide encoding genes were identified in C. elegans and parasitic nematodes

Our pan-phylum motif-based and reciprocal BLAST approach enabled us to identify nine novel Allatostatin-C-like neuropeptide encoding genes (*nlp*-87, -88, -89 and -90, 91, -92, -93, -94 and -95), in addition to those previously reported [*nlp*-55, -62, -64, -66, -67, -71 and -79; (see Mirabeau and Joly, 2013; Van Bael et al., 2018).

Four of the novel Allatostatin-C-like neuropeptide encoding genes (*nlp*-87, -88, -89 and -90) were found in the *C. elegans* genome, shared a common [RNCFF(S/T)P(V/A)QC] motif, possess a signal peptide, and appear to be enriched in neurons (see WormBase (WS276): WBGene00010848 (M04B2.6); WBGene00016436 (C35B1.7); WBGene00019292 (K02A6.1); WBGene00019293 (K02A6.1)). *nlp*-88, -89 and -90 appear to be restricted to Caenorhabditids, whereas *nlp*-87 is conserved across parasitic nematodes (Table 1; Fig. S4). The additional five novel Allatostatin-C-like neuropeptide encoding genes identified here (*nlp*-91, -92, -93, -94 and -95) share a common [(K/R)NC(F/Y)F] motif, are conserved across key parasite species, but are absent from *C. elegans*. Again, CLANS analysis broadly supports the peptide motif based novel gene designations highlighted here (Fig. 2 B).

#### 3.6.3 Novel NLP families are absent from Caenorhabditis elegans

Two additional novel putative NLP-encoding genes, that display distinct peptide motifs (NPYSW), and (S(L/V)AP(T/S)TSAX_3-4_VS), were identified in this study and have been designated *nlp*-96 and -97 respectively; these novel *nlps* were not observed in *C. elegans*. *nlp*-96 sequelogues were identified in a broad range of parasites, spanning clades 8, 9, 10 and 12 (Table 1), whilst *nlp*-97 appears to be *Globodera spp.* specific (see Fig. S1). Note that *G. pallida* and *G. rostochiensis* each possess two highly similar *nlp*-97 sequelogues (*nlp*-97.1, -97.2); we also note recent *nlp* duplications in parasitic nematodes for <u>*nlp*</u>- X -X -X and have designated these paralogues as *nlp*-X.1, *nlp*-X.2 etc (see Figure S1).

### 3.7 Highly conserved NLPs are bioactive on the Ascaris suum ovijector

To determine the potential role of the most highly conserved NLPs in regulating muscle function, we examined the effects of: *As*-PDF-1 (A and B); As-PDF-2 (C) *As*-NLP-12 (A1, B, C, D), NLP-36 (A1, A2, B1, B2, C1, C2), *As*-NLP-47 (C) and *As*-NLP-49 (B) on the reproductive musculature (ovijector) of *A. suum*, using established methods (Fellowes et al., 1998; Fellowes et al., 2000; Moffett et al., 2003). Note that NLP-12 (A1, B, C and D), and NLP-36 (A1, A2, C1 and C2) peptides were inactive (see Table S6).

NLP-36B1, -36B2, -47, PDF-1A, -1B peptides exhibited distinct excitatory effects on ovijector muscle (Fig. 3, Table S6) as follows; (i) *As*-NLP-36B1 and *As*-NLP-36B2 induced an increase in contraction amplitude and frequency that aligned with the previously described ovijector response type (RT) 5 (Moffett et al., 2003); (ii) *As*-NLP-47 induced a transient lengthening of the ovijector with an increase in contraction frequency, similar to ‘RT2’ type responses, whilst (iii) *As*-PDF-1A and *As*-PDF-1B caused a decrease in contraction amplitude alongside an increase in contraction frequency that aligned to RT5. By contrast, *As*-NLP-49 was the only peptide tested shown to induce relaxation of the ovijector muscle (RT1; Fig. 3, Table S6).

**Fig. 3.**
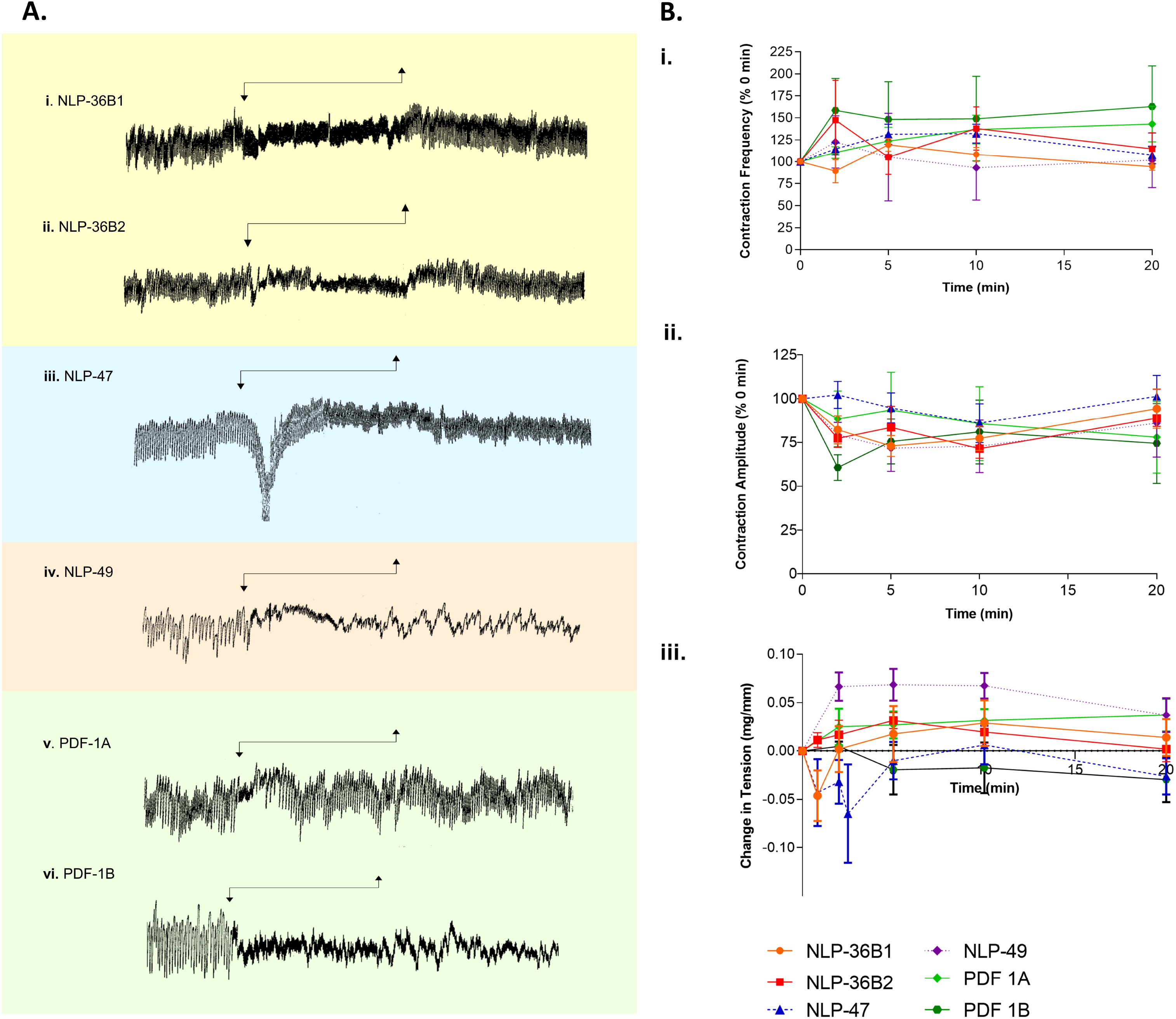
Selected NLP predicted peptides are bioactive on the *Ascaris suum* ovijector muscle. (A) Representative muscle tension recordings showing the effect of *As*-NLP-36, *As*-NLP-47, *As*-NLP-49, and *As*-PDF-1 peptides. Arrows indicate peptide exposure. (B) The effects of NLP peptides on *A. suum* ovijector contraction frequency (i), contraction amplitude (ii) and baseline tension (iii). Peptide present from 0-10 minutes.

The muscle physiology described here represents the first reports of NLP activity on the *A. suum* ovijector (Moffett et al., 2003; Mousley et al., 2004; Mousley et al., 2005; McVeigh et al., 2006b; McVeigh et al., 2008). Intriguingly, *pdf*-1, *nlp*-47 and *nlp*-49 are all linked to the control of egg-laying in *C. elegans* (Lindemans et al., 2009; Meelkop et al., 2012; Chew et al., 2018), a role which may be conserved in parasitic nematodes.

This study provides a comprehensive library of NLPs in nine key parasitic nematodes, and highlights that parasites possess a reduced and variable complement of the *C. elegans* NLP profile. We identify 14 novel *nlps*, 10 of which are not found in *C. elegans*, bringing the total number of *nlps* in nematodes to 99. Five *nlps* display complete conservation across the nine phylogenetically dispersed parasitic nematodes examined here, four of which encode peptides that modulate nematode muscle function. The data presented here advance our understanding of neuropeptide signalling in parasitic nematodes, support the possibility for conservation of neuropeptide function across multiple nematode species, and could direct neuropeptide-receptor deorphanisation studies in therapeutically relevant parasite species.

## Acknowledgements

The authors are grateful to Karro, Cookstown, NI for assistance in the collection of nematodes, and wish to thank Dr Isabel Beets and Professor Timothy G Geary for helpful comments on the data. The authors acknowledge support for this work from: Biotechnology and Biological Sciences Research Council grant (BB/H019472/1), Biotechnology and Biological Sciences Research Council/Boehringer Ingelheim grants (BB/MO10392/1, BB/T016396/1), and Department for the Economy Northern Ireland (DfE).

