## Supplementary Figure 1 for "*In silico* analyses of neuropeptide-like protein (NLP) profiles in parasitic nematodes"

**Figure S1**: **Nematode neuropeptide-like protein (NLP) sequelogue alignments from *Caenorhabditis elegans* and parasitic nematodes.** Multiple sequence alignments were performed using Vector NTI Advance 11.5 AlignX® (Lu and Moriyama, 2004). Italic text indicates the presence of a signal peptide as identified using Signal P 4.1 (Nielsen, 2017). Ce-NLP-56A, B and C represent isoforms of the *C. elegans nlp*-56 gene. Parasite sequelogues designated NLP-X.1, -X.2 etc. represent paralogues (likely recent gene duplications) of *C. elegans* NLPs in these species. Note: pan-phylum alignments are not an exhaustive list of all NLP-sequelogues across all nematodes. Ce denotes *C. elegans*, Ts denotes *Trichinella spiralis*, Tm denotes *Trichuris muris*, As denotes *Ascaris suum*, Bm denotes *Brugia malayi*, Di denotes *Dirofilaria immitis*, Na denotes *Necator americanus*, Hc denotes *Haemonchus contortus*, Bx denotes *Bursaphelenchus xylophilius*, Gp denotes *Globodera pallida*. Yellow and blue highlighted regions indicate completely and partially conserved amino acid residues respectively. Green indicates similar residues.

NLP-1

```
                 1                                                         60
 Ce-NLP-1    (1) MKATFVLACLLVIAAVSHADLLPKRMDANAFRMSFGKRSVSNPAEAKRMDPNAFRMSFGK
 As-NLP-1    (1) ---------------MEVKLKKENNRFFFNFIKRYLERVIFHRAFGKRMDASKFYMGFGK
 Na-NLP-1    (1) ------------------------------------------------------------
Hc-NLP-1.1   (1) -----------------------------------------------------------M
Hc-NLP-1.2   (1) -----------------------------------------------------------M
Gp-NLP-1.1   (1) -----------------------------------------------------------M
Gp-NLP-1.2   (1) ---------------------------------------------------------MAS
Gp-NLP-1.3   (1) ------------------------------------------------MSPNLFIVFSLL
                 61                                                       120
 Ce-NLP-1   (61) RSAEQNEQANKEDKATSDKLYDDTKFEEMKRMDANAFRMSFGKRSDAHQAADDQVEYVND
 As-NLP-1   (46) RDGYIRRLPEINRTIIDSEVHLQFSGKAAKEDGCNSEEFSLNAKATS--LAR-------H
 Na-NLP-1    (1) ----------MVIVAALSQDYDEFIGGKRAAMDPHAFRMSFG------------------
Hc-NLP-1.1   (2) NNFRLTVLSFMVVVAVFSQEFEVFTEGKQAEMYPRALPFRMSRG----------------
Hc-NLP-1.2   (2) DNLRITILSLMIVVAVFTQEYEVFTEGKRAVMFPRTFGALFG------------------
Gp-NLP-1.1   (2) SRKFELFLAITLLFMAQSNKRSFVALAELQNQDRMAQIRTLP------------------
Gp-NLP-1.2   (4) FQYFAVILFSLLILRLITLNEVRAAPTGGLQLDRSAFRMSFGKRSP--AAAAESDSSELN
Gp-NLP-1.3  (13) LLAVPLRLANAEPEFVMAKKETEVG-PTGFQMDRNAFRMSFGKRDEALVPFRMLR-----
                 121                                                      180
 Ce-NLP-1  (121) DFSLPEQKRMDANAFRMSFGKRVNL-------------------------DPNSFRMSFG
 As-NLP-1   (97) IYEPLMEKRMDASRFYLGFGKRADEQAPYG----------N-ILLNLLDSERLETEIAVD
 Na-NLP-1   (33) --KRFVGRNIDPNAFR--MSFGKRS-------------------------ASPQLEIVNP
Hc-NLP-1.1  (46) --KRSTGIRFDPNASRTRMSFGKR--------------------------NAYGSNEWNE
Hc-NLP-1.2  (44) --KRLTGNRIGLNTFR--MGSGIQ--------------------------KAYGINDWNE
Gp-NLP-1.1  (44) --SVG--LQMDPNVFQIGFGKREQQ-------------------------QNVQQPMEAF
Gp-NLP-1.2  (62) NANIAIPLMYDMSEAETTMKQQQQP---------------R-HKVFPSGGVGRAPWPSRW
Gp-NLP-1.3  (67) --SNPRQLAMDRNAFRMSFGKRSPAGSSSSNELINANIDANHQQKYRAFWSPPAANGWEA
                 181                 203
 Ce-NLP-1  (156) KRSTVGYNLDARNYFVGLGRR--
 As-NLP-1  (146) GVVKSPSELEPGVHTNHTLKKSA
 Na-NLP-1   (64) YDQELVKRMDMKHYYIGLGKR--
Hc-NLP-1.1  (78) YLKENTKRMDMRHDFVGLGNR--
Hc-NLP-1.2  (74) YIKDNDKRMDMKHYFVGLGRR--
Gp-NLP-1.1  (75) LLDRNVFRMSFGKRELVKN----
Gp-NLP-1.2 (106) KASLFSKRLDRNLYNIGFGR---
Gp-NLP-1.3 (125) AAALPPKRLDRNLFNVRFGRRRR
```

NLP-2

```
                 1                                               50
 Ce-NLP-2    (1) ---------------------------------------MRATLVLFALL
 As-NLP-2    (1) --------------------------------------------------
 Na-NLP-2    (1) MERKSCEVNNIAIFDYVDDDDVLAYKQVEVNSPFELGGYENEVNGLEPYI
 Bx-NLP-2    (1) ----------------------------------------MARTIFGLFF
                 51                                             100
 Ce-NLP-2   (12) CAVYSEAVPLQVYRPDESSAVDVVVLENSPELYDSEDEDEWKQEEEFTEG
 As-NLP-2    (1) -----------RYDLRFSNFVFKAISESE--FPAELQKRSMALGRLAFR-
 Na-NLP-2   (51) YIRYNCDKPSYRSIMFRMTFIASLMLRNKSWRADMLPLVGFLTISAIWVV
 Bx-NLP-2   (11) GPNKVAKSTVTACRCADRSVIRQTNIVQLRARAPANPFSKEMWPRIMMTF
                 101                                            150
 Ce-NLP-2   (62) AMGKRSIALGRSGFRPGKR---SMDNFHTVDVSDLIMKRSMAMGRLGLRP
 As-NLP-2   (37) -PGKRSLALGRVDFRPGKRNVDIADKDIEYLMEDSMGKRSAAFGRFHFRP
 Na-NLP-2  (101) AWAGLTSPLGASLGNESPREILSVLPLHVVTRRPMDEFSPRVRTRLGRYK
 Bx-NLP-2   (61) AFTSMVMLIEAAPHTAGRN---AEDGEVVLMAAENPNARYLYMQPQEQVE
                 151                                            200
 Ce-NLP-2  (109) GKRSMAYGRQ----GFRPGKRSMAYGRQGFRPGKRSMAYGRQG-------
 As-NLP-2   (86) GKRSLALGRS----GFRPGKRSLALGRVGFRPGKRASGEVLIDPCSIEID
 Na-NLP-2  (151) PEDDVSNETQRFFTEERMGKRSIALGRSGFRPGKRSDSLESHG-------
 Bx-NLP-2  (108) PDWAEVFSQP----DPRMEKRSVALSRLNFRPGKRSVPLATQADLVEQG-
                 201                       229
 Ce-NLP-2  (148) ---------FRPGKRSNDMKEVFPQHVPE
 As-NLP-2  (132) SLWGDMYDIAQVGMRSNFSREAFYYC---
 Na-NLP-2  (194) ---------LRPGRGSFDEM---------
 Bx-NLP-2  (153) -----IYRLNALQKRSVAISRKGFRPAKK
```

NLP-3

```
                 1                                     40
 Ce-NLP-3    (1) MSKIVACLVLLALSVMCVYSAPYEFRAKR-----AINPFL
 As-NLP-3    (1) ---MQPLSVTAQHTGLATHAPPTNAPFPL-----LLYFAL
 Bm-NLP-3    (1) MTTAICLLMV---MILLSYRTESQYVSRQGNYIDYNRGDL
 Di-NLP-3    (1) MTKATCLLMVIWSMILLAYRTESQYASRRGFYVDHNRGDL
 Na-NLP-3    (1) --MLKAVFLSLLVLVSIVFTAYADMRAKR-----AINPFL
 Hc-NLP-3    (1) --MLKGMVFSLLVFVTLLFSAYSDTRAKR-----AINPFL
 Bx-NLP-3    (1) MSPFRGVSLASVLFAVLFLATTLSAFLQDEVP---LITEI
 Gp-NLP-3    (1) MFSSSTLPVLLQLVAVIFISNPSEAYSAG----RLYLRDS
                 41                                    80
 Ce-NLP-3   (36) DSMGKRAVNPFLDSIGKR-----------SFRPDMITEEK
 As-NLP-3   (33) FARFRRAINPFTDSIGKRN----------TGIAAERDLPK
 Bm-NLP-3   (38) YMRNFGIINPFSDSDSRR-----------IRFDSSRELPK
 Di-NLP-3   (41) YMRNFGVINPFSDADSKR-----------IRFDNSRELPK
 Na-NLP-3   (34) DSMGKRALNPFLDSIGKRA----------SQYQPYYGYEK
 Hc-NLP-3   (34) DSMGKRAVNPFLDSFGKRS----------SRYQPYYHLDK
 Bx-NLP-3   (38) EGVAKRAINPFMDSIGKRSGNVYVPGSRFYRHSLLFQRPS
 Gp-NLP-3   (37) YPDSARFFIPNALLADQLLASGPVSSLRPALFDRSPIPAK
                 81                                   120
 Ce-NLP-3   (65) RYFDSLAGQSLGKRSNNRYEMLENYY--------------
 As-NLP-3   (63) RYFDALAGQSLGKRASPIYYLVDE----------------
 Bm-NLP-3   (67) RYFDALAGQSLGKRSTTIL---------------------
 Di-NLP-3   (70) RYFDALAGQSLGKRSATIL---------------------
 Na-NLP-3   (64) RYFDSLAGQSLGKRSIQTYDA-------------------
 Hc-NLP-3   (64) RYFDSLAGQALGKRSIQAYDA-------------------
 Bx-NLP-3   (78) RYFDSLAGQSLGRR--------------------------
 Gp-NLP-3   (77) RYFDSLAGQSLGKRTAPPATEEEMPRGDDARVNSQRAMRL
                 121
 Ce-NLP-3   (91) ----------
 As-NLP-3   (87) ----------
 Bm-NLP-3   (86) ----------
 Di-NLP-3   (89) ----------
 Na-NLP-3   (85) ----------
 Hc-NLP-3   (85) ----------
 Bx-NLP-3   (92) ----------
 Gp-NLP-3  (117) DALRPGWPMW
```

NLP-5

```
                 1                                               50
 Ce-NLP-5    (1) --------------------------------------------------
 Ts-NLP-5    (1) MVHHNWESHKWTTSFERCDQFLPDEPTVEPELAKIRHQVVSPLTDGIDLF
 Tm-NLP-5    (1) --------------------------------------------------
 Na-NLP-5    (1) --------------------------------------------------
 Hc-NLP-5    (1) --------------------------------------------------
 Bx-NLP-5    (1) --------------------------------------------------
                 51                                             100
 Ce-NLP-5    (1) ------------------------------------------------ML
 Ts-NLP-5   (51) TWQLSAFIAPLTSSFSSTYCRQYKRFPPNARLPLEATSLCTVVHFLQSSK
 Tm-NLP-5    (1) --------------------------------------------------
 Na-NLP-5    (1) -------------------------------------------MYVVVLL
 Hc-NLP-5    (1) ------------------------------------------MYIILFSC
 Bx-NLP-5    (1) --------------------------------------------------
                 101                                            150
 Ce-NLP-5    (3) MKIMVLMGIANIAASFVVSSVRFQSAPMRALIEINRELAKRSVSQLN---
 Ts-NLP-5  (101) HTMVLKATLNGLLLFWLMISTQLISTLTTARFIYNIPSDTKSLSDRTFEK
 Tm-NLP-5    (1) --MSALRGSPRALLLAIIMSTQLISNVCLGRYLRVYPEELRALRNFDGDS
 Na-NLP-5    (8) FVVAQTACSTENLDTSEHDKRALSSFDTLAGIGLGKRAQLSSIDSLAGLG
 Hc-NLP-5    (9) LVAAQTVLSETAGFDLTGDKRALSSFDTLGGIGLGKRTQLSSIDSLGGLG
 Bx-NLP-5    (1) ----MLGIYVNLLLLTVLASAEIIPSNVLHKRQAKMEIKKDNLGYSD---
                 151                                            200
 Ce-NLP-5   (50) ----QYAGFDTLGGMGLGKRSEPDQAGEKRALSTFDSLGGMGLGKRSSSS
 Ts-NLP-5  (151) R-LPTQLYFDPLGGMALGKRAG----PSAISGQYFDPLAGMAFGKRAG-L
 Tm-NLP-5   (49) HRLVIRQYFDPLGGMALGKRAS-DLDRIGSRLQFFDPLGGMALGKRSP--
 Na-NLP-5   (58) LGKRALSSFDTLGGIGLGKRSF---VVVKKAVSSFDTLGGIGLGKRSE--
 Hc-NLP-5   (59) LGKRALSSFDTLGGIGLGKRSE---DTAKKALSSFDTLGGIGLGKRDD--
 Bx-NLP-5   (44) ----EMFNIDTLAGIGLGKRSS--PTPTRPRYALIVNLEDLVEGLDRS--
                 201                                            250
 Ce-NLP-5   (96) SRVFVYDKRALQHFSSLDTLGGMGFGRK----------------------
 Ts-NLP-5  (195) IRPTWNAGFPEPALNSSHYLDMNVSSVDLSTLNNPL--------------
 Tm-NLP-5   (96) -----------PAKFHPDALDGALWNSPFAR-------------------
 Na-NLP-5  (103) --LFRGDK---KSLSSFDTLAGIGLGKRSRPLTQQYYIPFRGIMVSANRK
 Hc-NLP-5  (104) --MLAGEK---KSVSSFDTLAGIGLGKRSRLFSTYYYLPYRDSLEDMDQN
 Bx-NLP-5   (86) ------AQ---KVRARFNRFHEI---------------------------
                 251
 Ce-NLP-5  (124) ---
 Ts-NLP-5  (231) ---
 Tm-NLP-5  (116) ---
 Na-NLP-5  (148) EQD
 Hc-NLP-5  (149) VQE
 Bx-NLP-5  (100) ---
```

NLP-6

```
               1                                                    55
 Ce-NLP-6    (1) -------------------------------------------------------
 Ts-NLP-6    (1) -------------------------------------------------------
 Tm-NLP-6    (1) -------------------------------------------------------
 Na-NLP-6    (1) MAVSKFFSSFGVQRFQDILLREFVDIEIFTDLAECCTNASNPYDGCIHSCKHRRR
 Bx-NLP-6    (1) -------------------------------------------------------
                 56                                                  110
 Ce-NLP-6    (1) -------------------------------MLSFSRLAFVLLVSACVMAMAAPK
 Ts-NLP-6    (1) -----------------------------------MNIFITLFAFIVVMQQFSVD
 Tm-NLP-6    (1) -----------------------------MAATVLPKVYSALLCLFVIVHSFASF
 Na-NLP-6   (56) QNPEKFSLQVFPVRFHSLRKSHGCALTLSVKMARITRFLVCLLLAVVGLLNASPK
 Bx-NLP-6    (1) -----------------------------MHSLVITTLLLAAISNSVIGLGYIYE
                 111                                                 165
 Ce-NLP-6   (25) QMVFGFGKRS------MADISDMPEDVPMKRYKPRSFAMGFGKRAAMRSFNMGFG
 Ts-NLP-6   (21) SNELSNNGRR--------IHFQNYFQRMQKRLDPQYFQLGFGKRFIVNEENYPTN
 Tm-NLP-6   (27) EPMLIMAGAQ----P---LWKAPRSSSAKRPFDPYHFQMGFGKRVASYAAQDQPF
 Na-NLP-6  (111) QHVFGFGKRG----GEALGDPEVDWNAMSKRLSNGKFMMGFGKRGLNKQFTMGFG
 Bx-NLP-6   (27) DAYYPIPRGSQIVV--EAPSIPDEKVIPQRRFDPAWRHIGLGKRTAEERRTVSGE
                 166                                                 220
 Ce-NLP-6   (74) KRSAEPQIDYEDMSADLEEIPVFIQK----------------RRLIMGLGRR---
 Ts-NLP-6   (68) MQCCAQLQLLAAMEDGF--------------------------------------
 Tm-NLP-6   (75) SVVKLCADVDNGRTLMKNCRHYVIAE-----------------------------
 Na-NLP-6  (162) KRVPADEELLSEDYIDQSGRTSLLTKGLGKRLYGLNDLPGLRTIQYMGLGKRSVL
 Bx-NLP-6   (80) RSWAMIGLGR---------------------------------------------
                 221            238
 Ce-NLP-6  (110) ------------------
 Ts-NLP-6   (85) ------------------
 Tm-NLP-6  (101) ------------------
 Na-NLP-6  (217) SREVAANLRRFNLGLGRR
 Bx-NLP-6   (90) ------------------
```

NLP-7

```
                 1                                               50
 Ce-NLP-7    (1) ---------MYIKAALLIVVLFGVASQITS-------------------A
 As-NLP-7    (1) MSYFSPLSAVAIILYASSVMSSSMFRSDDS------------------HA
 Na-NLP-7    (1) ---------MIFLYLLLGVVSASVLQYDPR---------------YLMSH
 Hc-NLP-7    (1) ---------MIFVYVCLGVVSASVLHYEPRTVLHPLPSKRLVIPSYLSSH
 Bx-NLP-7    (1) ------MMRSLILFLFGLLMPVLLAKSEPS--------------------
                 51                                             100
 Ce-NLP-7   (23) LYLKQADFDDPRMFTSSFGKRSAIESEPQAYPKSYRAIRIQRRS------
 As-NLP-7   (33) YHNSFLTSGMPVELIAAENEGNPARFNHIQGAASTKNKRFADEW------
 Na-NLP-7   (27) YDKRTLDFDDPRLFSSAFGKRSGFPARSLS-FQIVKRSPYFGS-------
 Hc-NLP-7   (42) YDKRTLDFDDPRLFSTAFGKRNGLVTTTLNRPRFIKRSPFLGTNVGLAYI
 Bx-NLP-7   (25) -AG-GVYEFNPRDWKADMSPHNFYRFRRYN--------------------
                 101                                            150
 Ce-NLP-7   (67) MDDLDDPRLMTMSFGKRMILPSLADLHRYTMYDKRG---SDIDDPRYFLF
 As-NLP-7   (77) --D--DPRFLSSAFGKRSGMMEFDDPRFLSSAFGKRNHLIGFDDPRLFSS
 Na-NLP-7   (69) -DD--S-----EDTYEKRDGMELMDPRFFTNSFGKR-SDFLFDDPRFSSV
 Hc-NLP-7   (92) KRADMDPRFISNSFGKRSTLYDFDDPRFASLSFGKR-SGFDFDDPRFSSM
 Bx-NLP-7   (53) -----------VFSNDKRSEMEFDDPRYFSTAFGKRASSFL---------
                 151                                            200
 Ce-NLP-7  (114) SNRLTCRC------------------------------------------
 As-NLP-7  (123) SYGKR----------ALFELQKKYDAAFDDPRFLSLAFGKK---------
 Na-NLP-7  (110) SFGKRR--------------------------------------------
 Hc-NLP-7  (141) SFGKRSGFDLEDPRFASMSFGKRSGFNFEDPRFASLSFGKRSGFDLDDPR
 Bx-NLP-7   (83) --------------------------------------------------
                 201                      228
 Ce-NLP-7  (122) ----------------------------
 As-NLP-7  (154) ----------------------------
 Na-NLP-7  (116) ----------------------------
 Hc-NLP-7  (191) FASMSFGKRSGSDLEDPRYWSMSFGKRR
 Bx-NLP-7   (83) ----------------------------
```

NLP-8

```
                 1                                                    55
 Ce-NLP-8    (1) --------------------------------MSQKLLPISPLQLLFLQCLLIGF
 As-NLP-8    (1) MKTVSLPNRLLKACTGGRILRSKRAFDRLDESALGLFRKRSFDRIDGSAFGPHRH
 Na-NLP-8    (1) ----------------------------------MAGHTSTFMRCLLVKFAVLAV
 Hc-NLP-8    (1) -----------------------------------MTQYSSLLRLHLVQMVFLAL
 Bx-NLP-8    (1) -----------------------------------MTDWAGGERVPKTGGALGEH
Gp-NLP-8.1   (1) -----------------------------------MTSKGALLSFVLTLLTVDAY
Gp-NLP-8.2   (1) --------------------------------MAFRYSEAIFITIQLFNFFLLTN
                 56                                                  110
 Ce-NLP-8   (24) TAAYPYLIFPASPSSGDSRR--------LVKRAFDRFDNSGVFSFGAKRFDR---
 As-NLP-8   (56) KKAFDRIDGSDFGLVKRAFDRIEGAGFGLSKRAFDRIEGS-GFGLDKRAFDRLEG
 Na-NLP-8   (22) CSAYPYLLDPDYELSDVIVS----------KRAFDRLEGN-EFGLLKRALN----
 Hc-NLP-8   (21) CSAYPYVLNSDYELTDVAERPFG----GLTKRAFDRIESN-DFGLFKRSAA----
 Bx-NLP-8   (21) KILGKVMRTIFTVLTVCFLQVLCLS---QGQNTEDRSREQ-------R-------
Gp-NLP-8.1  (21) PAFYMVRNLPQLQHPQNALQ----------SSELSRLRPS---------------
Gp-NLP-8.2  (24) AMNMHDFLWASAVRPAQSSQ---------NSALTDTLSYETNKSPAKLLMTVP--
                 111                                                 165
 Ce-NLP-8   (68) ---------YDDETAYGYGFD-NHIFKRSADP-YRFMSVPTKKAFDRMDN-----
 As-NLP-8  (110) SDFGLVKRAFDRLEGSDFGLD-KRAFDRVGESNFGFDKRSFDRVADSTAFRLSRG
 Na-NLP-8   (62) ------KRAFDRLDMADFGLRRKRAFDRIARAEFGLTGLRRKRSSDLSTSNEE--
 Hc-NLP-8   (67) ------KRAFDRIEMADFGFRRKRAFDRVGRTEFGFEGVLRKRAADRLAD-----
 Bx-NLP-8   (59) --YVLHKRAFDRVDLSPFDFG---AYSKRYNNDLRSLYRKKKSFDRLDQGPFG--
Gp-NLP-8.1  (51) ------KRAFDRLDASPFDFD--SISKRFSDDELAAMPLNDLYLSSPYAFG----
Gp-NLP-8.2  (68) -----FKRAFDRLDVSNFDFN-----KRFMEVLDVAPPPRSKLLFNIHRMPSELP
                 166                                                 220
 Ce-NLP-8  (107) --------------------SDFFGAKRKRSFDRMGGTEFGLMKRSAPE-----S
 As-NLP-8  (164) AFSRIDDNDFGLFKLNAVNPGRRIHLLQKRPFDRIERSEFGLSKRFDSSALPAVR
 Na-NLP-8  (109) -----------------------FGLRRRRALDRIGGSEFGLIKRSVDPHL---S
 Hc-NLP-8  (111) -----------------------IGFRNKRGLDQLDGTDLGLMFDPVRP-----A
 Bx-NLP-8  (107) ------------------------LVKRKRVFDRVDAGFGFGKRSSEMAVPGRIS
Gp-NLP-8.1  (94) ------------------------PF-RKRSFDRLEESAFFGQKRKRSFGP----
Gp-NLP-8.2 (113) --------------------------NKRKLAEGLKRRRWAMQAKQQLQAAG---
                 221                                    262
 Ce-NLP-8  (137) REQLINNLAESIITLRRAREAESSPESQRTIITYDD------
 As-NLP-8  (219) YVEINPELVDLLPLDDVVYSFVKQDVVPFNSFTIADEIPRKR
 Na-NLP-8  (138) REELINDLADSIVSLHEDSALHGPKLVVFPVPAEERESNR--
 Hc-NLP-8  (138) REELIDRLAYTIAAMG-----HATPVAISAVPLVDDQK----
 Bx-NLP-8  (138) LSDLVSQAQSIEPID---------------------------
Gp-NLP-8.1 (120) -ERILRDN----------------------------------
Gp-NLP-8.2 (139) RGPTVSRVLGFGLSFGGMRRLRRTMLK---------------
```

NLP-9

```
                 1                                               50
 Ce-NLP-9    (1) ------MDRFATRFIALLLVLLQIGSIFATPIAEAQGAPEDVDDRRELEK
 As-NLP-9    (1) ---------------------------------------IGFIWRRLIQK
 Bm-NLP-9    (1) MLTMKLFLHCAIFYLLTYAHDINDDDKERTPILNNPEAEDLERQLFWTRK
 Di-NLP-9    (1) ----------------------------------------------MMKK
 Na-NLP-9    (1) ----------------MKNSCESVSGSLQNVFTDSGNSVRDARSWTIVAK
 Hc-NLP-9    (1) -----------------MPSSSTYAALLTIVLLVCLSAAQNSENYPSLAK
                 51                                             100
 Ce-NLP-9   (45) RGGARAFYGFYN-----AGN-SKRDQAAALPYYLYEKRGGGR-----AFN
 As-NLP-9   (12) RAGARSFGPRIYDVESYLYP-TNKRFGSFSPYYFYQQPK-----------
 Bm-NLP-9   (51) RAGARAFGSMRGNNFENLFFPSNKRHQTFSPYYYYQQLK-----------
 Di-NLP-9    (5) RGGARAFGPMRSNNYEHLFVPNIKRHQTLSPYYYYYQQPK----------
 Na-NLP-9   (35) RGGARAFHSYFN-----VPN--AKRLSVAAPYYLYEKRGGGRAFVGGWQP
 Hc-NLP-9   (34) RGGARAFHGYFN-----MPS--SKRLSGEYPYYLYEKRGGGRAFFGGWQP
                 101                                            150
 Ce-NLP-9   (84) HNANLFRFDKRGGGRAFAGSWS-------PYLERFYDYKRSSYPVYFSDN
 As-NLP-9   (50) ----------RGGGRTFASYWIPPSGNGRSLLFGNIFFI-----------
 Bm-NLP-9   (90) ----------RGGGHPFYTYWSPSNAGNVKFVEGSPFYKKRSVLRDSDDY
 Di-NLP-9   (45) ----------RGGGHPFYTYLS-ENGGDGKFIEHLPFYKKRSLMHDSDDY
 Na-NLP-9   (78) FDGISGRMERRGGGRAFSGWWG-PNEYFSRFYENPYYKRSSNLWEFLEDR
 Hc-NLP-9   (77) YESLGARMDRRGGGRAFSGWLG-PNEYY-RFYENPYYKRSSSLWEFLEDR
                 151
 Ce-NLP-9  (127) SYY
 As-NLP-9   (79) ---
 Bm-NLP-9  (130) IY-
 Di-NLP-9   (84) AY-
 Na-NLP-9  (127) NGF
 Hc-NLP-9  (125) NAL
```

NLP-10

```
                  1                                          45
 Ce-NLP-10    (1) ---------------------------------------MWYIAL
 As-NLP-10    (1) ---------------------------------------------
 Bm-NLP-10    (1) ---------------------------------------------
 Di-NLP-10    (1) MINSHLLPLIIHARAGSNRRPSACKADVITTTPLDRCSVTDHFFN
 Na-NLP-10    (1) ---------------------------------------------
Hc-NLP-10.1   (1) ---------------------------------------------
Hc-NLP-10.2   (1) -----------------------------------MLQDVLVMLE
 Bx-NLP-10    (1) ---------------------------------------------
 Gp-NLP-10    (1) ---------------------------------------------
                  46                                         90
 Ce-NLP-10    (7) LLAVIATSVTAQKADDEPIVFLVRVPIDEMDDDSSLLESYYHP--
 As-NLP-10    (1) -----------------MKNILDVLPFSGGLYG------------
 Bm-NLP-10    (1) ---------------------------------------------
 Di-NLP-10   (46) VSKISSKRCPNFIGNINQNIDLFCIKREIRNYYVFCFISTKNSLR
 Na-NLP-10    (1) -------------MLVPSLLFLVAVAYANEEEESSSSSDVVLLDL
Hc-NLP-10.1   (1) -MDCFGKGNPLETMIVPSLLFLVAVAYANEDEGMYGKRSEMPDDM
Hc-NLP-10.2  (11) SIPQLTTGNPLETMIVPSLLFLVAVAYANEDEDLASSS-------
 Bx-NLP-10    (1) ----------MRTVILCLILSVGCIGMASAASIYIDSN-------
 Gp-NLP-10    (1) ----MSSATISPLLLLPVLLVLLSVFGCHATPPSQHSQTEEN---
                  91                                        135
 Ce-NLP-10   (50) -----------RDILSKRAIPFNGGMYGKR-STMPFSGGMYGKRS
 As-NLP-10   (17) ---------------KRSSIPFHGGIYGKRTAALPFSG---G---
 Bm-NLP-10    (1) ----------------MVAILSEFLIYIILIYKIISAKEVFDGEV
 Di-NLP-10   (91) LVIGNTFNAKFNLRTVMVAILSEFLIYAILIHKITSANEAFDKKI
 Na-NLP-10   (33) SDYAPGINRGLRFDIKRAAMPFSGGMYGKR-AVMPFSGGMYGKRD
Hc-NLP-10.1  (45) YIERPVLPLSAGWQEKRAVMPFSGGLYGKR-AVMPFSG-------
Hc-NLP-10.2  (49) -------------EAIMIVPDYAPEILGR-----AFNN-------
 Bx-NLP-10   (29) ---------------RPDYEENLQKVYDNRLQPYAIEY-------
 Gp-NLP-10   (39) -------------VLLEELVREVRKLSNRMDSALGRGGAAEEEEL
                  136                                       180
 Ce-NLP-10   (83) G------QIFAQRRAAIPFSGGMYGKRS---LVPQSYSNNENQIK
 As-NLP-10   (41) --------LYGKRAGWIPFSGGLYGKRA---LTPLQGGIYG---K
 Bm-NLP-10   (30) QNVG--GGEAQKNDYPLPLSSNLYGKRVNYVLMPLMGGMYG---K
 Di-NLP-10  (136) RNG---FGHVENDEHSPSLSNNLYGKRVNYALMPLMGGMYG---K
 Na-NLP-10   (77) GPALGFSEGWAEKRAVMPFSGGVYGKRAP--ALPFSGGVYG---K
Hc-NLP-10.1  (82) --------GWQEKRAVMPFSGGLYGKRA---VMPFSGGLYG---K
Hc-NLP-10.2  (69) ----------GAKRAVMPYSGGMYGKRA---VMPFSGGLYG---K
 Bx-NLP-10   (52) ----------LAKRAALPYSGGIYGKRA---ALPYSGGIYG---K
 Gp-NLP-10   (71) IVE---GKRSPQGMAPSVLWNGAGDKRA---QLPFSGGIYG---K
                  181                                       225
 Ce-NLP-10  (119) RGAMPFSGGMYGR--------------------------------
 As-NLP-10   (72) RAVIPFSGGLYGKR--------TSQDFTKQNRL----------SL
 Bm-NLP-10   (70) RAHSDIYDKSYQPLSLSAADEYNNKRLSLLSKIIHKENDMAQFNP
 Di-NLP-10  (175) RTFVPFHGGMYGKRRSLPFNRYYNKQLLSPEITLNED-----LIP
 Na-NLP-10  (117) RAAMPFSGGLYGKR--------SERYIRSP---------------
Hc-NLP-10.1 (113) RAAMPFSGGLYGKR--------ADRYIRSP---------------
Hc-NLP-10.2  (98) RSVMPFSGEAVHLI--------MVLNER-----------------
 Bx-NLP-10   (81) RASVPFSGGIYGKRS------VDEAPVDHVNGI----------SI
 Gp-NLP-10  (107) RSPLPFSGGIYGKRS------SSHQLYRSPLIFPTLD-----RQT
                  226                                       270
 Ce-NLP-10  (132) ---------------------------------------------
 As-NLP-10   (99) RSIPMSGGFFG----------------------------------
 Bm-NLP-10  (115) LSHYHINEKVGHSPYQTLRQHYQLEDDNNDYYNQINDNSHLLQLL
 Di-NLP-10  (215) SVYYDGNGKFGQFPYKLWRQYQLKEDNSDYYKDNSYPSQPFPELV
 Na-NLP-10  (139) --MPISGGFFG----------------------------------
Hc-NLP-10.1 (135) --MPISGGIFG----------------------------------
Hc-NLP-10.2 (118) ---------------------------------------------
 Bx-NLP-10  (110) RAMPINGGFYG----------------------------------
 Gp-NLP-10  (141) RAMPFNGGMYGR---------------------------------
                  271                                       315
 Ce-NLP-10  (132) ---------------------------------------------
 As-NLP-10  (110) ---------------------------------------------
 Bm-NLP-10  (160) LKLRSLLMNDEFNN-------------------------------
 Di-NLP-10  (260) KPSNQFDYSFFKHNMQLLILSMFLLLAKFLFAENTLCIPPSIPAK
 Na-NLP-10  (148) ---------------------------------------------
Hc-NLP-10.1 (144) ---------------------------------------------
Hc-NLP-10.2 (118) ---------------------------------------------
 Bx-NLP-10  (121) ---------------------------------------------
 Gp-NLP-10  (153) ---------------------------------------------
                  316                                       360
 Ce-NLP-10  (132) ---------------------------------------------
 As-NLP-10  (110) ---------------------------------------------
 Bm-NLP-10  (174) ---------------------------------------------
 Di-NLP-10  (305) DSQNVDCCKLPREIFPHKTIYEEFCSVLYFGVMADGPDGRFCRLS
 Na-NLP-10  (148) ---------------------------------------------
Hc-NLP-10.1 (144) ---------------------------------------------
Hc-NLP-10.2 (118) ---------------------------------------------
 Bx-NLP-10  (121) ---------------------------------------------
 Gp-NLP-10  (153) ---------------------------------------------
                  361                                       405
 Ce-NLP-10  (132) ---------------------------------------------
 As-NLP-10  (110) ---------------------------------------------
 Bm-NLP-10  (174) ---------------------------------------------
 Di-NLP-10  (350) SSQKRQSINCDPGWIAFYEGLNQFCYKAFTINSTDKIVDIMSKKD
 Na-NLP-10  (148) ---------------------------------------------
Hc-NLP-10.1 (144) ---------------------------------------------
Hc-NLP-10.2 (118) ---------------------------------------------
 Bx-NLP-10  (121) ---------------------------------------------
 Gp-NLP-10  (153) ---------------------------------------------
                  406                                       450
 Ce-NLP-10  (132) ---------------------------------------------
 As-NLP-10  (110) ---------------------------------------------
 Bm-NLP-10  (174) ---------------------------------------------
 Di-NLP-10  (395) GDPCRKKKETARLVRIPNQNINMFLYVRVLNGAQNLLCDSVDRNT
 Na-NLP-10  (148) ---------------------------------------------
Hc-NLP-10.1 (144) ---------------------------------------------
Hc-NLP-10.2 (118) ---------------------------------------------
 Bx-NLP-10  (121) ---------------------------------------------
 Gp-NLP-10  (153) ---------------------------------------------
                  451                                       495
 Ce-NLP-10  (132) ---------------------------------------------
 As-NLP-10  (110) ---------------------------------------------
 Bm-NLP-10  (174) ---------------------------------------------
 Di-NLP-10  (440) YYYYIGLYRNDFDWRWRSYDEKAYSHLEEQGQQLLDYLNQPRSNH
 Na-NLP-10  (148) ---------------------------------------------
Hc-NLP-10.1 (144) ---------------------------------------------
Hc-NLP-10.2 (118) ---------------------------------------------
 Bx-NLP-10  (121) ---------------------------------------------
 Gp-NLP-10  (153) ---------------------------------------------
                  496                      523
 Ce-NLP-10  (132) ----------------------------
 As-NLP-10  (110) ----------------------------
 Bm-NLP-10  (174) ----------------------------
 Di-NLP-10  (485) TWPDGTRIFADGWHDVDDITTNVVMSNC
 Na-NLP-10  (148) ----------------------------
Hc-NLP-10.1 (144) ----------------------------
Hc-NLP-10.2 (118) ----------------------------
 Bx-NLP-10  (121) ----------------------------
 Gp-NLP-10  (153) ----------------------------
```

NLP-11

```
                 1                                                    55
Ce-NLP-11    (1) -------MMSTLALVSLAIFGIAVVCAAPKPATVPVANEEDYLAALYGFEAPGSQ
As-NLP-11    (1) -----MSKSILTLFTLLFIFAVAVIATREKRQLSPAVQLENELSPLIRLAGLHDG
Bm-NLP-11    (1) MYLIDLATVTAALAICFGDFVCAYGNHRQKKQLSPALQLEDNINSLLRLNELGY-
Di-NLP-11    (1) MYLIDLSTIT-ALAICFGDLMCGYASR-QKKQLSPALQLQDDINSLLRLNELSP-
Na-NLP-11    (1) ------MMLAQIAVLTIAVCVAGANSVRVKRQLSPAVRLESELHPLMMGMYGFG-
Hc-NLP-11    (1) ---MQVMALTQIVVLAFVVCMVGANSIRAKRQLSPAVRLETELHPLVMGMYGFGP
                 56                                                  110
Ce-NLP-11   (49) FKGAPLQS----KRHISPSYDVEIDAGNMRNLLDIGKR-SAPMASDYGNQFQMYN
As-NLP-11   (51) ---GYYKRSFDYKRHLSPSLNVEDQLGRLIALQSVGKR--HSPAEQYGQQFELTL
Bm-NLP-11   (55) -KRS--------FGEPSHAMEFHDGLDDLFSLQNIGKRPALSPADQFARQMDAMQ
Di-NLP-11   (53) -KRS--------SGDQPYVLPVMN-LDNLLSLQNIGKRPALSPADQFIQQFDIMQ
Na-NLP-11   (49) -SEASKRSPLSLKRHISPSFDVEEDVGNLRTLMDIGKR-QVSLADDVGRQMQMYH
Hc-NLP-11   (53) ENNAYKRSALSLKRHISPSFDVEEDVGNMRTLMDIGKR-QLSVADDVGRQMQMYH
                 111                                                 165
Ce-NLP-11   (99) RLIDAGKKKRSPAISPAY-----QFENAFGLSEALERAGRR--------------
As-NLP-11  (101) SISRGRLPFKVTASSNQSEFNKHYLKRIDNIVASTSRSGQKTCHLTIGRFTQSTI
Bm-NLP-11  (101) HLAAAGKK---RSLSPAT-----DFQQQLQLPHMLMDAGR---------------
Di-NLP-11   (98) HLMAAGKK---RSVSPAT-----DLEEQIKIPQVLIDVGR---------------
Na-NLP-11  (102) RLFDAGKKR--AVLSPSA-----DLQNAMDLTSYLERAGRK--------------
Hc-NLP-11  (107) RLFEAGKKR--AALSPSQ-----DLQSAVELSNYLERAGRK--------------
                 166        179
Ce-NLP-11  (135) --------------
As-NLP-11  (156) MVRVSRSSRSIVAF
Bm-NLP-11  (133) --------------
Di-NLP-11  (130) --------------
Na-NLP-11  (136) --------------
Hc-NLP-11  (141) --------------
```

NLP-12

```
                 1                                          45
Ce-NLP-12    (1) ---------------------------------------------
Ts-NLP-12    (1) ---------------------------------------------
Tm-NLP-12    (1) ---------------------------------------------
As-NLP-12    (1) ---------------------------------------------
Bm-NLP-12    (1) ---------------------------------------------
Di-NLP-12    (1) ---------------------------------------------
Na-NLP-12    (1) ---------------------------------------------
Hc-NLP-12    (1) ---------------------------------------------
Bx-NLP-12    (1) ---------------------------------------------
Gp-NLP-12    (1) MLLGICGFLSLSLMVVGFIAYSRRFYTYAYGRANAQMRFRRAVYT
                 46                                         90
Ce-NLP-12    (1) ---------------------------------------------
Ts-NLP-12    (1) -----------------------------------------MVII
Tm-NLP-12    (1) -----------------------------------------MITS
As-NLP-12    (1) ---------------------------------------------
Bm-NLP-12    (1) -------------------------------------------ML
Di-NLP-12    (1) ---------------------------------------------
Na-NLP-12    (1) ---------------------------------------------
Hc-NLP-12    (1) ---------------------------------------------
Bx-NLP-12    (1) --------------------------------------------M
Gp-NLP-12   (46) VRVRKSPRAPMGFQFGDFGEDTVVFNTLSCSVRHISTFNADDAII
                 91                                        135
Ce-NLP-12    (1) MLRHHSCALLMLILVFVEVFATQSPT-------------------
Ts-NLP-12    (5) RVSLMVFFAWLASFFLGQAALSQSQQ----VAKYNRAGFSPYLKR
Tm-NLP-12    (5) RMMITAILVSFLMLLMGQTRSASQQQQQQPSFKLNRNGFVPFFKN
As-NLP-12    (1) -MFGWIYLLLLLCIASSIHCDLTPTR-------------------
Bm-NLP-12    (3) RSWITSILFILGFISIAVNCSLQSER-------------------
Di-NLP-12    (1) ---------------------------------------------
Na-NLP-12    (1) --MIHCLLLLALVCTVLATADAQSQT-------------------
Hc-NLP-12    (1) --MIHYLLLLALACSVLATADAQSPT-------------------
Bx-NLP-12    (2) KLYILLMLALVGWVCAAAASEEEADR-------------------
Gp-NLP-12   (91) IIVFQHHAVLPNCFTTCHHCSNIAERHKQQAMFFDAVDVSSVGTV
                 136                                       180
Ce-NLP-12   (27) -----FDRQDRDYRPLQFGKRDGYRPLQFGKRD--YRPLQFGKRS
Ts-NLP-12   (46) FPALPSQYLKRSDMNNMLGIGVKRDDDQFDREDRDYRPLMFGKRY
Tm-NLP-12   (50) LPPLPRQYLKREILLPYYNAMAKREDDQFDREDRDYRPLMFGKRF
As-NLP-12   (26) -----FDRQDRDYRPLQFGKRDGYRPLQFGKKS--YRPLQFGKRR
Bm-NLP-12   (29) -----FDRQDRDYRPLQFGKRDGHRPLQFGKKS--FRPLQFGKRG
Di-NLP-12    (1) ---------------LQFGKRDGHRPLQFGKKS--FRPLQFGKRQ
Na-NLP-12   (25) -----FDRQDRDYRPLQFGKRDGYRPLQFGKRDS-YRPLQFGKRS
Hc-NLP-12   (25) -----FEREDRDYRPLQFGKRDGYRPLQFGKRD--YRPLQFGKRS
Bx-NLP-12   (28) -----FSRTDRDYRPLQFGKRGDFRPLQFGKKDN-YRPLQFGKRA
Gp-NLP-12  (136) SGE-QFERADRDYRPLQFGKRESYRPLQFGKRIVASRGLLFAPAV
                 181                            214
Ce-NLP-12   (65) SGSSGPVVLEPIWEWQ------------------
Ts-NLP-12   (91) PDYQLPMFG-------------------------
Tm-NLP-12   (95) PDYQPPMFG-------------------------
As-NLP-12   (64) A-YDFSEYTPY-----------------------
Bm-NLP-12   (67) M-DEAPDYFYDI----------------------
Di-NLP-12   (29) S-ERFDRQDRDYRPLQVG----------------
Na-NLP-12   (64) P--LASAYLVPVL---------------------
Hc-NLP-12   (63) P--LASAFLVPAL---------------------
Bx-NLP-12   (67) AVVNHYLDLLPIDEIAN-----------------
Gp-NLP-12  (180) APPAFMHNLLHNSGLEEMDPTMDQYVLMVEERRK
```

NLP-13

```
                  1                                               50
 Ce-NLP-13    (1) --MQRSLQIFCIMSAIAMAYSQGSRDDNQSAKRNDFSRDIMSFGKRSGN-
 As-NLP-13    (1) --MKAIPIQFLLLSLFMFVNSQIFYDVG---TKREFDRDFMHFGKRS---
Bm-NLP-13.1   (1) -----MTILILISLAAAIVQSEYYAN-----YNDRLARDFMHFGKRSPNI
Bm-NLP-13.2   (1) --METMRILILISLAAAIVQSEYYAN-----YNDPLTRDSMNFEKRAVPS
Bm-NLP-13.3   (1) --METMRILILISLAAAIVQSEYYAN-----YNDPLTRDSMNFEKRAVPS
Di-NLP-13.1   (1) --MEIITVLIVVSAVVITVQSEYYVN-----YDGQLARDFMHFGKRS---
Di-NLP-13.    (1) --------------------------------------------------
 Na-NLP-13    (1) --MTPPWSLVCLAVTITAVSSQYILATGDNEKRNDFSRDIMHFGKRS---
 Hc-NLP-13    (1) -MMMPPWSLVCLAVTVTVASTQYLSDGGENEKRNDFSRDIMHFGKRA---
 Bx-NLP-13    (1) MSTQTHFLAVVAATLCCLILCDAEDAFYSIGDKRNFDREFMHFGKRATS-
                  51                                             100
 Ce-NLP-13   (48) TADLYDRRIMAFGKRQ--PSYDRDIMSFGKRSAPSDFSRDIMSFGK----
 As-NLP-13   (43) --NAFDRNFMNFGKRE--SQFSRDFLNFGKRESNVRSLKQYYDDLR----
Bm-NLP-13.1  (41) IIPQFDRNIASLKR----GEFSRDFMNFGKRSIG--DDWSDITLKVGDNP
Bm-NLP-13.2  (44) MRNMINLNKRSDSDN---GAFDREFLHFGKREVP--SKHNMINLNK--RS
Bm-NLP-13.3  (44) MRNMINLNKRSDSDN---GAFDREFLHFGKREVP--SKHNMINLNK--RS
Di-NLP-13.1  (41) -----PDMIQQFDGDN--GSPKRGE-------------------------
Di-NLP-13.2   (1) ----NFCLFISIFF----FKFSRDFLNFGKRSVD--DNWSDTMIRIDDTP
 Na-NLP-13   (46) ---VLGLSALGNSN----PDFDRDMMSFGKRTG---LEREFMSFGK----
 Hc-NLP-13   (47) ---YGNGRLVAYGG----PAFERDMMAFGKRSGG--FEREMMSFGK----
 Bx-NLP-13   (50) --EDFGRSFMPFGKRATNDDFGRAFMPFGKRAANEEFGRAFMPFGKR---
                  101                                            150
 Ce-NLP-13   (92) --RSSS--MYDRDIMSFGKRS---------PVDYDRPIMAFGKR----AE
 As-NLP-13   (85) ----QK-KDFDRDFMHFGKR----------SDAFSRNFMNFGKR----DD
Bm-NLP-13.1  (85) ELLHKKSSTFDRDFMNFGKR----------AVPFERNLINSNKKSGIDSG
Bm-NLP-13.2  (87) --DSDNG-AFDREFLHFGKR----------EVPSKHNMINLNKRSDSNSG
Bm-NLP-13.3  (87) --DSDKG-AFDREFLHFGKR----------EVPSKHNMINLNKRSDSDNG
Di-NLP-13.1  (59) --------------------------------------------------
Di-NLP-13.2  (41) ESVHKKS-NFDREFMNFGKR----------MIPYDRNLINFNKKNSVKAD
 Na-NLP-13   (82) --RS----PYEREFLPFGKRD---------LSDFNREMMSFGKR-----E
 Hc-NLP-13   (84) --RS----PFEREFLSFGKR----------GSEFDREMLSFGKR-----D
 Bx-NLP-13   (95) --ADAYDESFNRAFMPFGKRSPMTLDQLIAKKNMNRDFMHFGKR--VAPS
                  151                                            200
 Ce-NLP-13  (125) DYERQIMAFGRRK-------------------------------------
 As-NLP-13  (116) AFSRDFLSFGKRR-------------------------------------
Bm-NLP-13.1 (125) AFDREFLSFGKR--------------------------------------
Bm-NLP-13.2 (124) AFDREFLHFGKR--------------------------------------
Bm-NLP-13.3 (124) AFDREFLHFGKREVPSKHNMINLNKRSDSDKGAFDREFLHFGKREVPSKH
Di-NLP-13.1  (59) --------------------------------------------------
Di-NLP-13.2  (80) AFKREFLSFG----------------------------------------
 Na-NLP-13  (112) DFDRNIMPFGRRRR------------------------------------
 Hc-NLP-13  (113) EFERSMMAFGRRRK------------------------------------
 Bx-NLP-13  (141) SFDREFMHFGKRR-------------------------------------
                  201                    226
 Ce-NLP-13  (138) --------------------------
 As-NLP-13  (129) --------------------------
Bm-NLP-13.1 (137) --------------------------
Bm-NLP-13.2 (136) --------------------------
Bm-NLP-13.3 (174) NMINLNKRSDSNSGAFDREFLHFGKR
Di-NLP-13.1  (59) --------------------------
Di-NLP-13.2  (90) --------------------------
 Na-NLP-13  (126) --------------------------
 Hc-NLP-13  (127) --------------------------
 Bx-NLP-13  (154) --------------------------
```

NLP-14

```
                 1                                          45
Ce-NLP-14    (1) ---------------------------------------------
As-NLP-14    (1) ---------------------------------------------
Bm-NLP-14    (1) -----MLSLILLVLALGEFNDAKSKFDEIKKMRDTIKGSRFRAKQ
Di-NLP-14    (1) MTAYQMLSLVLLLLELDKINCAKSEFNGMAKTQDPVKDS--RSNE
Na-NLP-14    (1) ---------------------------------------------
Hc-NLP-14    (1) ---------------------------------------------
Bx-NLP-14    (1) ----------------------------------MTVHSRICLSA
Gp-NLP-14    (1) ---------------------------------------------
                 46                                         90
Ce-NLP-14    (1) -------------------------------MLHLIVLLVALSSA
As-NLP-14    (1) -----MRNTFISLETRHEKYILRFSDSRLPRFILQFISATSISRS
Bm-NLP-14   (41) SLNSIDGSEFDGLGGIRIRIEKRSLDALQGEGFGMKKRALDALEG
Di-NLP-14   (44) KMRQIEEFRR-G---------KRALDALDDDGLGMDLRNMKASDD
Na-NLP-14    (1) ------------------------MLFSLARCVLLVGCAQFICGL
Hc-NLP-14    (1) ------------------------MLFFLTRCVLVVGLSQLVCSE
Bx-NLP-14   (12) ALFVMFCSAFVSSAVVVRLPVAPTRSVFFTPALFTKRALDALEGS
Gp-NLP-14    (1) --------------MAEAKNICRTKTPFSDFKLMVLLGIVLCSLF
                 91                                        135
Ce-NLP-14   (15) VTAGRPRRALDGLDGSGFG-FDKRALNSLDGAGFGFEKRALNSLD
As-NLP-14   (41) KDAKK--PSLDILEGAGFSPLKKRALDTLEGSGFGFSKRALDDLE
Bm-NLP-14   (86) EGFGMKKRALDALEGEGFG-MKKRALDALEGEGFGMKKRALDALE
Di-NLP-14   (79) AESRREKRSVDSVDGDGYG-MRLRYLGALEDDGFGLKKRALDELE
Na-NLP-14   (22) PEATHSPPALDDLDGNGFGGLKRVLSRRK---------RALDSLE
Hc-NLP-14   (22) QDATHPPPALDDLEGNGFGGMKKFSSRRK---------RALDSLE
Bx-NLP-14   (57) DFGLKRKRALDSLEGSDFG-LRKRALDSLEGSDFGLRKRALDSLE
Gp-NLP-14   (32) TPSTSIKESSRSSSLGRTGAIKVLALRPR---------RALDILE
                 136                                       180
Ce-NLP-14   (59) GQ--GFGFEKRALDGLDGAGFG-FDKRALNSLDGA-GFGFEKRAL
As-NLP-14   (84) GVGFGGMLRKRALDSLEGTDFG-LKKRALDYLEGS-GFGLMKRAF
Bm-NLP-14  (130) GE--GFGMKKRALDALEGEGFG-MKKRALDALEGE-GFGMKKRAL
Di-NLP-14  (123) GE--GFGLKKRALDALEGEGFG-FNKRALDALEGE-GFGFNKRAL
Na-NLP-14   (58) GGGFGFDKKKRALDELDGSGFG-FDKRALNALEGT-GFGFDKRAL
Hc-NLP-14   (58) GDGFGGLFKR-SLDSLEGDGFG-FDKRALNALDGT-GFGFDKRAL
Bx-NLP-14  (101) GA--DFGLKKRALDSLEGADFG-LKKRALDSLEGA-DFGLRKRAL
Gp-NLP-14   (68) SDD-FGGFRKRALDVMDGDGFGSFEKRALDTLEGDDFMGLKRKRL
                 181                                       225
Ce-NLP-14  (100) DGLD----GSGFGFDKRALNSLDGAGFGFEKRALNSLDGAGFGFE
As-NLP-14  (127) DAIEDDLEGAGFGLKKRALDAMEGAGFGFDKRALDSMEGTGFGFH
Bm-NLP-14  (171) DALE----GEGFGMKKRALDALEGEGFGMKKRALDALEGEGFGMK
Di-NLP-14  (164) DALE----GAGFGFKKRALDELEGAGFGLKKRALDSLEGTGFGIE
Na-NLP-14  (101) NSLE----GEGFGFDKRALNSLDGSGFGFDKRALNSLEGEGFGFD
Hc-NLP-14  (100) NSLE----GTGFGFDKRSLNSIEGTGFGFDKRSLNSIEGTGFGFD
Bx-NLP-14  (142) DSLE----GADFGLRKRALDSLEGSDFGLKKRALDALEGTGFGFD
Gp-NLP-14  (112) NELEG---DGFMGLDKRALDILDGDDFTGFSKRSEVNGG--LRRE
                 226                                       270
Ce-NLP-14  (141) KRALDGLDGAGFGFDKR--ALNSLDGAGFGFEKR---ALDGLDGA
As-NLP-14  (172) KRALDGLEGDGFGFTKK--ALDSLEGTDFDFGKRSYGPLKHLVFD
Bm-NLP-14  (212) KRALDALEGEGFGMKKR--ALDALEGEGFGMKKR---ALDALEGE
Di-NLP-14  (205) KKALDALEGAGFGLRKR--ALDSLEGAGFGIEKK---ALDALEGA
Na-NLP-14  (142) KRALNSLEGEGFGFDKR--ALNSLEGEGFGFDK---RALNSIEGD
Hc-NLP-14  (141) KRSLNSIEGTGFGFDRRRRSLDSTEGTGFGYRR---GKRTLPQIT
Bx-NLP-14  (183) KRALDMLEGSDFGLKKR--ALDSLEGADFGLKKR---ALDSLEGS
Gp-NLP-14  (152) LALLGKVQRQRQMRARR--ALDALEGNSFGFRR------------
                 271                                       315
Ce-NLP-14  (181) GFGFDKRALNSLDGNGFGFDKRTFKHS-----SN--KLRSVFRNL
As-NLP-14  (215) RNRPSLRYSLAQSNKNRKRLRSQLVRL--LEHHNYWPVFVVLLRR
Bm-NLP-14  (252) GFGMKKRALDALEGEGFGIDKRVLDALEGAGLRMNRPLKSSDSRT
Di-NLP-14  (245) GFGLKKRVIDVLKNAGYRRIRPKMNKGRFDGMRN-YPFQSSGLRT
Na-NLP-14  (182) GFGFDKRYYSHIINRNTSTLQNLKKQG--FDGRTAETRLTL----
Hc-NLP-14  (183) GTHPYLRLYKGKQSRGYKGDRSNP------NGSNTYLHNN-----
Bx-NLP-14  (223) DFGLRKRALDSLEGSDFGLKKRSPIARFYATGDDIRKLNDLKTQL
Gp-NLP-14  (183) ---------------------------------------------
                 316                                  355
Ce-NLP-14  (219) KGFKQH----------------------------------
As-NLP-14  (258) SLIKPYAMKRDQAASDWLRSLPHA----------------
Bm-NLP-14  (297) KGFHMNRFSTNGRHSHPHLTKF------------------
Di-NLP-14  (289) NRFRINRYFSNGRHSRPSKMIISSGNFAIRFISLSSIFLS
Na-NLP-14  (221) ----------------------------------------
Hc-NLP-14  (217) ----------------------------------------
Bx-NLP-14  (268) EVELRRRLEKEAEA--------------------------
Gp-NLP-14  (183) ----------------------------------------
```

NLP-15

```
                  1                                          45
 Ce-NLP-15    (1) MPSSSSSSSFFAAVLLVIVMMSTVESAAVRLRPVGSLFFLNRPHE
 Ts-NLP-15    (1) -----------------------MYRALGGKKNFNQFDVGLSGFS
 Tm-NLP-15    (1) ------MLTVHQIKAQSDWRFDNVVGSEQRPLTAMNNLYKALAQK
 As-NLP-15    (1) ---MNRLFSFMAKKTICSTFWQMNTEVTFLKNGAPMVVRIAGNAE
 Na-NLP-15    (1) ----------------MNSTAAPAMRVVLLIFTTMVVVSSAPSRQ
Hc-NLP-15.1   (1) -------------------------------------ILSAPSRQ
Hc-NLP-15.2   (1) ----------------MNSSAASALRIALFVCATVAIISSAPSRQ
Bx-NLP-15.1   (1) ------------------------------------------MDR
Bx-NLP-15.2   (1) ------------------------------MNTQFLRFSVILIVL
 Gp-NLP-15    (1) ----------------KRSFDSLTGPGFTGLDTRAAFDTDFTNYD
                  46                                         90
 Ce-NLP-15   (46) KRAFDSLAGSGFDNGFNKRAFDSLAGSGFGAFNK-----------
 Ts-NLP-15   (23) KRPFDRLEGAHFSG-FDKRTLDMLEGSSMSGFNKRYFDVPRLYIK
 Tm-NLP-15   (40) KYSFDRLDASGMSG-FNKRPFDRFESAQIGGLNK-----------
 As-NLP-15   (43) KRAFDSLTGAGFTG-FDKRAFDSLTGSGFTGFDK-----------
 Na-NLP-15   (30) KRAFDSFAAPGFDA-FNKRAFDTLAGQGFEAFNK-----------
Hc-NLP-15.1   (9) KRAFDTFAGS-FSG-FDKRAFDSLAGSGFSAFDK-----------
Hc-NLP-15.2  (30) KRAFDTFAGS-FSG-FDKRAFDSLVSFFFLLN-------------
Bx-NLP-15.1   (4) KRSFDSFTGAGFTG-MDKRSFDALTGSGFTGMDK-----------
Bx-NLP-15.2  (16) GCVRAALAGVANTG--EKRAFDALTARGFNGFDK-----------
 Gp-NLP-15   (30) KRSFDSFTGPGFTG-LDKR-FEPFDGYGFNGFEK-----------
                  91                                        135
 Ce-NLP-15   (80) RAFDSLAGSGFGAFNKRA--FDSLAGSGFSGFDKRAFDSLAGQGF
 Ts-NLP-15   (67) RNFDKFDSATLSGFRRKR-MLDNIDSAGFAFSNKTLLYQQLHFTF
 Tm-NLP-15   (73) KAFDMLENSKLGGFQKRP--FDSIETGDFKEGRAPV---------
 As-NLP-15   (76) RSLDSPANRGFTHFDKRA--FDSLTGAGFTGFDKRSFDSLNSRGF
 Na-NLP-15   (63) RAFDSLSGSGLTPFNKRA--FDSLAGNGFGGFDKRSPLVRRSYAI
Hc-NLP-15.1  (41) RAFDSLSGSGLTPFNKRA--FDSLAGSGFTGFDKRSPIALNAGYY
Hc-NLP-15.2  (60) -AFGNLLKPMIFPLLLG----------------------------
Bx-NLP-15.1  (37) RSFDSFTGAGFTGMDRKR---------ALARMMAQ----------
Bx-NLP-15.2  (48) RAFDSF---------------------------------------
 Gp-NLP-15   (62) RSFDSFMGPGFTGMDKKKRAFDSFTGPGFTGMDKK----------
                  136                                       180
 Ce-NLP-15  (123) TGFEKRAFDTVSTSGFDDFKL------------------------
 Ts-NLP-15  (111) RSLSWYEIINCYINLSSIILMRSIALA------------------
 Tm-NLP-15  (107) ---------------------------------------------
 As-NLP-15  (119) TGFDKRAFDSLVGHGFTGFD-------------------------
 Na-NLP-15  (106) VVLHIDLPSFIIRIIALVMLYLPDIKNYPERETNSVDQHKRISDV
Hc-NLP-15.1  (84) QPLREDALLL-----------------------------------
Hc-NLP-15.2  (76) ---------------------------------------------
Bx-NLP-15.1  (63) ----KRAFDSMTGRGFTGFD-------------------------
Bx-NLP-15.2  (54) ---------------------------------------------
 Gp-NLP-15   (97) ----KRAFDLFTGPGFTGMDRK-----------------------
                  181                                       225
 Ce-NLP-15  (144) ---------------------------------------------
 Ts-NLP-15  (138) ---------------------------------------------
 Tm-NLP-15  (107) ---------------------------------------------
 As-NLP-15  (139) ---------------------------------------------
 Na-NLP-15  (151) YALIPVTYKRWLLSLLYTNSKSTGITVGAECLSKETKHKTRRRHA
Hc-NLP-15.1  (94) ---------------------------------------------
Hc-NLP-15.2  (76) ---------------------------------------------
Bx-NLP-15.1  (79) ---------------------------------------------
Bx-NLP-15.2  (54) ---------------------------------------------
 Gp-NLP-15  (115) ---------------------------------------------
                  226     236
 Ce-NLP-15  (144) -----------
 Ts-NLP-15  (138) -----------
 Tm-NLP-15  (107) -----------
 As-NLP-15  (139) -----------
 Na-NLP-15  (196) ADTQQTHRLGC
Hc-NLP-15.1  (94) -----------
Hc-NLP-15.2  (76) -----------
Bx-NLP-15.1  (79) -----------
Bx-NLP-15.2  (54) -----------
 Gp-NLP-15  (115) -----------
```

NLP-17

```
                 1                                               50
Ce-NLP-17    (1) --------------------------------------------MFSKSI
As-NLP-17    (1) ------------------------------------------------MQ
Na-NLP-17    (1) ---------------------------------------------MRQSV
Hc-NLP-17    (1) ---------------------------------------------MRQCF
Bx-NLP-17    (1) MNLCFHCPPFHPFNLFFVVSRKSRVVAGRVIFAGQTYCNRERLTEHSFQL
Gp-NLP-17    (1) -----------------------------------------MLLNLLAVY
                 51                                             100
Ce-NLP-17    (7) LFCLLVLVFNVFGANFEN--DQDVMRPPFQALKRGSLSNMMRIGKRQ--M
As-NLP-17    (3) TVVVIITLALLTLPSSRCIPRYGRMIGTEQWTSNTSQDTVEQSSSDSDVM
Na-NLP-17    (6) VPILLVWFICLASAQFVDVDRPEILRPDVRFLKRNNLSNMMRLGKRG--L
Hc-NLP-17    (6) VLLLLWIVSAVCMQALDEGTREELMSPDVRFLKRNSLSIMMRLGKRVDAL
Bx-NLP-17   (51) GFSFSTTYNRDVSNLHNHKEMAVLVNSYVWQLIVLALVCLACTFATAQPV
Gp-NLP-17   (10) AVALMTTLYMPVQSAYTYQKMDELMPELRQYLGRTHLGTLKARKFNDDDN
                 101                                            150
Ce-NLP-17   (53) SRQQEYVQFPNEG------VVPCESCNLGTLMRIGRR-------------
As-NLP-17   (53) PLWWTHVR--------------ERKNILSNMMRIGRRSLSNP--------
Na-NLP-17   (54) RRQM-----------------LCEDCNLGNLMRLGRR-------------
Hc-NLP-17   (56) KEQE-----------------PCVDCSLGNLMRLGRR-------------
Bx-NLP-17  (101) GILGLETQGFNESPFEQIQLRERKSNNLRNLMRIGRRTP--------ADQ
Gp-NLP-17   (60) AFQQIR---------------ERKSNNLRNLMRIGKRNVPIADNFAREDE
                 151                       179
Ce-NLP-17   (84) -----------------------------
As-NLP-17   (81) -----------------------------
Na-NLP-17   (74) -----------------------------
Hc-NLP-17   (76) -----------------------------
Bx-NLP-17  (143) PIVPG-FEMRQAPLDYAQFFRPNAAIF--
Gp-NLP-17   (95) GGVGALSPVFYIPTDFLQAYRPQSVWLRR
```

NLP-18

```
                 1                                                         60
Ce-NLP-18    (1) -----------MNANVYSIVYFLSFLVLCISAQLHADSGATEVDG-----------IVDK
As-NLP-18    (1) ------------MRTFLALVLVGSLFAGVFCQELAEDFSPDKRGMSSFAKR--ARFNFAK
Bm-NLP-18    (1) -----------MASGFKVFVIFMSLLLTIIPCQEITDESLDRHDSSVFGKR--AKFAFAK
Di-NLP-18    (1) ----------------------------------MTDEVLNKHESSAFGKR--AKFAFAK
Na-NLP-18    (1) MCMVNKFQWVYIMQSMFTLAFVCTLTTMLLAQQVVDDVTAEKRVRS---------FAFAK
Hc-NLP-18    (1) ------------MQSTFTVVLVCTLASMILAQQMIDDVPAEKRMRS---------FAFAK
Bx-NLP-18    (1) ------------MHSTLGLIVCASLAVMVLSQELSEQDSMDKRARSFAFAKRARSFAFAK
                 61                                                       120
Ce-NLP-18   (39) RSP----YRAFAFAKRS--------DEENLDFLEKRARYGFAKRSPYRTFAFAKRASPYG
As-NLP-18   (47) RYP----SR-FAFAKRF--------DYDDWAEADKRAMRFAFAKRSMRNFAFAKRSP---
Bm-NLP-18   (48) RHP----SR-FAFAKRY--------DDEDVNYYDKKAMHFESPIHPTRNFAFAKRGP---
Di-NLP-18   (25) RYP----SR-FAFAKRY--------DDEDMYYYDKKAMHFEYPMHYMRNFAFAKRDP---
Na-NLP-18   (52) RSP----YRQFAFAKRS--------EDDDEEEVEKRARFAFAKRSPYRQFAFAKRASPYG
Hc-NLP-18   (40) RSP----YRQFAFAKRSDE------VVEDDGELEKRARFAFAKRSPYRQFAFAKRGSPYG
Bx-NLP-18   (49) KFDPYPFGRSFAFAKRSAEEPLLDVDSLEEDPMDKRARFAFAKRP--RNFAFAKRAR---
                 121       133
Ce-NLP-18   (87) FAFAKRGQFSSFA
As-NLP-18   (91) --------YSSFA
Bm-NLP-18   (92) --------YLSFA
Di-NLP-18   (69) --------YLLF-
Na-NLP-18  (100) FAFAKRG-YSSFA
Hc-NLP-18   (90) FAFAKRG-YSSFA
Bx-NLP-18  (104) ---------FAFA
```

NLP-19

```
                  1                                                 52
 Ce-NLP-19    (1) MLLRGVCLALLILVTIVQCQNDNDLKEEKRRIGLRLPNFLRFKDPDALMIHK
 As-NLP-19    (1) ---------------------KRRIGLRLPNILHMKVSIVVFQSYRHSSSQK
 Na-NLP-19    (1) -----------------------MPRFWYILFLLFIGSLP--IFSQTEDVYK
Hc-NLP-19.1   (1) ----------------------MISRIYCLLCLIFILSAVPNATSQEVELYK
Hc-NLP-19.2   (1) -MSSISPLHLGPLLILISAHFSMISRIYCLLCLIFILSAVPNATSQEVELYK
 Bx-NLP-19    (1) --------------------------------------MRPLNVESGERLQK
 Gp-NLP-19    (1) ----------------------------------SRLLNSPFRFCLSHIPDS
                  53                                               104
 Ce-NLP-19   (53) RRIGLRLPNMLKFKDSSNMYHLE-----------KRRMGMRLPNIIFLRNE-
 As-NLP-19   (32) RRFGLRLPNIIYIRRQRNLSFNYTI-------LQMACFGLRLPNIIYIRRQR
 Na-NLP-19   (28) RRIGMRLPNMLHYKPTAME---------------KRRIGMRLPNIIYLRSEP
Hc-NLP-19.1  (31) RRIGMRLPNMVHLKPLVR----------------KRRIGMRLPNIIYMRSEP
Hc-NLP-19.2  (52) RRIGMRLPNMVHLKPLVR----------------KRRIGMRLPNIIYMRSEP
 Bx-NLP-19   (15) RRLVMGMPNILHMGSKRMVKRMIERNAQRELVMNKRRISMHMPNLVYMRPQK
 Gp-NLP-19   (19) RKIGIRVPGFPIRQGYCS-----------------RKIGIRVPGFPIRQGYC
                  105                                              156
 Ce-NLP-19   (93) -KKNVLEY--------------------------------------------
 As-NLP-19   (77) ----------------------------------------------------
 Na-NLP-19   (65) EKKDNWEYDSF-----------------------------------------
Hc-NLP-19.1  (67) AKKNIWEYE-------------------------------------------
Hc-NLP-19.2  (88) AKKNIWEHQALLDRHDKNIISDTSTRDRRELLALFTPFWMTSRLRANPITIQ
 Bx-NLP-19   (67) ----------------------------------------------------
 Gp-NLP-19   (54) SRKIGIRVPGSDSAGIW-----------------------------------
                  157                           189
 Ce-NLP-19  (100) ---------------------------------
 As-NLP-19   (77) ---------------------------------
 Na-NLP-19   (76) ---------------------------------
Hc-NLP-19.1  (76) ---------------------------------
Hc-NLP-19.2 (140) TLFNNLYLNKFEWQHKNGKTSVLRKPWLGMTSQ
 Bx-NLP-19   (67) ---------------------------------
 Gp-NLP-19   (71) ---------------------------------
```

NLP-20

```
                 1                                     40
Ce-NLP-20    (1) ----------------------MQVTLLALLLLIVPFFAF
As-NLP-20    (1) ---------------------------------MSSTWSW
Na-NLP-20    (1) ----------------------KRMMCTLAVLILASIIAV
Hc-NLP-20    (1) MRTLKTRTARLALAQNTAKISSNKVMEAVPLFVILSTIVL
Bx-NLP-20    (1) -----------------------------MSPLLTTVTAL
                 41                                    80
Ce-NLP-20   (19) AASQYSDDDSELMSNNERFARDLELRKKFAFAFAKR----
As-NLP-20    (8) LCSFAFTPKKAGRFARHASGQQNE----LSFAFAKRAP--
Na-NLP-20   (19) SSGFAYDDEHNTDLARERFARELDQKQKFAFAFAKRGEDV
Hc-NLP-20   (41) STGFAYDDQENTDLTRDGFARDFA-KQRFAFAFANRYPEI
Bx-NLP-20   (12) FLFSSVYGQRQNAITDKVGQGNFAFARPFAFALTENGE--
                 81                                   120
Ce-NLP-20   (55) --------SAGD-------------ADVVIEAR---SGPQ
As-NLP-20   (42) --------FAKR------------EMQFAFEEPS-VDAER
Na-NLP-20   (59) PDAKQRFARDLENAKKFAFAFAKRSSDMVEESRSARFARD
Hc-NLP-20   (80) PEERSRFARDLDNSKKFSFAKK--SPDFSEESR--GFARE
Bx-NLP-20   (50) -----WYPIAKRSR----------TSTFAFKETPKEKERD
                 121                                  160
Ce-NLP-20   (71) AHEGAG------MRFAFAKRRAPKEFARFARASFA-----
As-NLP-20   (61) FARFAR------AAFAFAKRSPFAFA--------------
Na-NLP-20   (99) LDNDKK------FAFAFAKRLADDELRARYPRTPMEDQGF
Hc-NLP-20  (116) LEGNKK------FAFAFAKRFAGSHNEVRRQDNGRSNFAA
Bx-NLP-20   (75) SFAFAKRSEDPKFAYDFAKRAKFAFLKK------------
                 161               181
Ce-NLP-20  (100) ---------------------
As-NLP-20   (81) ---------------------
Na-NLP-20  (133) AREFAKRSDDRLMRFARPSFA
Hc-NLP-20  (150) PNFTNVHSVSEGV--------
Bx-NLP-20  (103) ---------------------
```

NLP-21

```
                  1                                     40
 Ce-NLP-21    (1) ----MRNSLFTTLFFGLAALVMVLN------AQYTSELEE
 As-NLP-21    (1) MIFIAKHQANSATFIGEKKCNQINTR-----WVNDCTFVS
 Na-NLP-21    (1) --------MLPPAAVSHADDDSIEK------RGGGRAFVP
Hc-NLP-21.1   (1) MRAVALVVIAAVAAISNAEEYSIEK------RGGGRAFVP
Hc-NLP-21.2   (1) -----------XAAISNAEEYSIEK------RGGGRAFVP
 Bx-NLP-21    (1) MDVTQKPMPNGERLVGTRPRGGVKKGREKVSELGFSIVAT
 Gp-NLP-21    (1) --RIVYIIPISSALFILPRIGVATS------ESDVDALAT
                  41                                    80
 Ce-NLP-21   (31) DEKRGGARAMLHK------RGGARAFSADVG---------
 As-NLP-21   (36) SNPLGHLKNFEKCS-------SIRLVVLLSEKQIYCMIAI
 Na-NLP-21   (27) IEERGGARAFFEDN----KRGGARVFPADKKGGAHA----
Hc-NLP-21.1  (35) IEERGGARAFFGDE----KRGGARAFVVDTRGGARA----
Hc-NLP-21.2  (24) IEERGGARAFFGDE----KRGGARAFVVDTRGGARA----
 Bx-NLP-21   (41) DRSWPGVVCKIAS----IMNLGVRTIDLFLWVSLFGILTQ
 Gp-NLP-21   (33) EQKRGGARAFNFFAPPDEKRGGARAFNFFAPDEKRGGARA
                  81                                   120
 Ce-NLP-21   (56) -----------------DDYKRGGARAFYDEKRGG-----
 As-NLP-21   (69) G----------------KRQLRIAQEPDVLEKRGG-----
 Na-NLP-21   (59) ----------------FRELKRGGARAFVPLEVRGGAVPS
Hc-NLP-21.1  (67) ----------------FREVKRGGGRAFLPVEERG-----
Hc-NLP-21.2  (56) ----------------FRDLKRGGGRAFLPVEERG-----
 Bx-NLP-21   (77) IVYGASFVQNLEDAELNEEDKRGGGRAFLNTDDKRG----
 Gp-NLP-21   (73) FNF------------FAPDEKRGGTRAFNFFVSDALPSSY
                  121                                  160
 Ce-NLP-21   (74) -------------ARAFLTEMKRGGARVFQGFEDE-----
 As-NLP-21   (88) -------------GRPFTLVDKRAGGRFFKIEDVDALE--
 Na-NLP-21   (83) Q-------LEKRGGGRAFVEPKRGGARAFYGIEQRGGARA
Hc-NLP-21.1  (86) ------------GARAFFKDDKRGGARAFVPVKEEDEGQS
Hc-NLP-21.2  (75) ------------GARAFFKDDKRGGARAFVPVKEEDEGQS
 Bx-NLP-21  (113) ------------GGRAFQFNDKRGGARAFAPVDVK-----
 Gp-NLP-21  (101) EKRSGIQTFRDDYDEKQAGELKRAGGRLFRMVDLPDGDDF
                  161                                  200
 Ce-NLP-21   (96) -----KRGGARAFMMD-------KRGGGRAFGDMM-----
 As-NLP-21  (113) -----KRGGGRPFYTQTDE----KRGGGRSFRLSSLGE--
 Na-NLP-21  (116) FFNDEKRGGARPFTDW-------KRGGARGFSTIE-----
Hc-NLP-21.1 (114) FTDLEKKGGARAFFVP-------KRGGGRAFLPIE-----
Hc-NLP-21.2 (103) FTDLEKKGGARAFFVP-------KRGGGRAFLPIE-----
 Bx-NLP-21  (136) ------RGGARAFAPID-----YKRGGGRAFSGFNGGN--
 Gp-NLP-21  (141) VPEGKKRGGARPFYGGGYMDGTWKRAGGRYFMRHFDDSPF
                  201                                  240
 Ce-NLP-21  (119) -----KRGGARAFVENSK----RDEDWVIR---PFEDDRL
 As-NLP-21  (142) -----KRMAAQKIAMPELDFYEGALAEIGNWP--------
 Na-NLP-21  (144) -----LRGGARAFFEDEK----RGGARAFK-RGGGRAFVL
Hc-NLP-21.1 (142) -----ERGGARAFFQEPK----RGGARAFPKRGGGRAFVP
Hc-NLP-21.2 (131) -----ERGGARAFFQEPK----RGGARAFPKRGGGRAFVP
 Bx-NLP-21  (163) YYLLNKRGGGRSFYGGFN----YGSLYAPSFPLSYEKKAD
 Gp-NLP-21  (181) AGWMAKRGGARAFFGDADGPFNSASYWAPRFVLPINKHPR


                  241                                  280
 Ce-NLP-21  (147) E---------------------------------------
 As-NLP-21  (169) --------------------------------------FI
 Na-NLP-21  (174) D---------------------------------------
Hc-NLP-21.1 (173) Q---------------------------------------
Hc-NLP-21.2 (162) Q---------------------------------------
 Bx-NLP-21  (199) YVP--------------------------------YYLTE
 Gp-NLP-21  (221) RSLLMGPLENVEDEAEQQRMDSSPSPFFTNNHNFDGRANG
                  281                                  320
 Ce-NLP-21  (148) -KRGGGRSFPVK-------------------P--------
 As-NLP-21  (171) EKRAGARSFPVLEDLLE-----------------------
 Na-NLP-21  (175) -KRGGARMFQNSKRDYTDDYPDWLMYADHEQSPFEVDENV
Hc-NLP-21.1 (174) -KRGGARIFQN----------------THKRP--------
Hc-NLP-21.2 (163) -KRGGARIFQN----------------THKRP--------
 Bx-NLP-21  (207) MKRAGGRAFQPKYYDP-------------YFWYVD-----
 Gp-NLP-21  (261) RKRGGARAFNGAEETLLNVANLAKRGGGRNFNEMEDTEEY
                  321                             355
 Ce-NLP-21  (160) -----GRLDD-------------------------
 As-NLP-21  (188) ------KRRIYDAFSRFP-----------------
 Na-NLP-21  (214) DVWQAGKRGGAHSFPVENSTKIRGNY---------
Hc-NLP-21.1 (189) -----WKRDDARSLPRTVESV--------------
Hc-NLP-21.2 (178) -----RKRDDARSLPRTVESI--------------
 Bx-NLP-21  (229) ----QTKRAGGRSFPISDESKEKRMLEEQADSRTI
 Gp-NLP-21  (301) K----KKRFGERIVSANPASCAMNFQPTQ------
```

NLP-35

```
                 1                                             48
Ce-NLP-35    (1) -MPRVSSFIVFLTFMVALLAVSHAAVVS-----GYDNIYQVLAPRFRR
As-NLP-35    (1) -MPRYASFVVLTAIILYVVESASSEQFG-------ES-YLPGFDELFG
Na-NLP-35    (1) --LRATLVILVVAVVLALCHALPIEENNPPVDYGSLDNFFYLSPRLRR
Hc-NLP-35    (1) MSLRAVLVILVVAVVMVLCHALPIDDTG--ADYGNLENFFYL-PRLRR
                 49                                            96
Ce-NLP-35   (43) ARMMSFDAEAPQYLQHLLQNLKPRFRRSV-------------------
As-NLP-35   (40) GRER--RDSKRAFLQQLMQRLKPRFRRDYAVNDHDITNFTRNRSELQL
Na-NLP-35   (47) DPLS--FGGQNAYLHQLLQNLKPKFRRSV-------------------
Hc-NLP-35   (46) DQLA--FSGQNAYLRQLLQNLKPKFRRSV-PLVPSIIANGLSLRDLSP
                 97           112
Ce-NLP-35   (72) ----------------
As-NLP-35   (86) S---------------
Na-NLP-35   (74) ----------------
Hc-NLP-35   (91) ALVDVTSMIFRAPVVY
```

NLP-36

```
                  1                                               50
 Ce-NLP-36    (1) ---------------MSVDLKQQLELADYLGALAVWCIFFGVLFILSVIF
Ts-NLP-36.1   (1) ----------------MVNLKQDVNVADYFGFIVVLVTFFVIIFVISTTC
Ts-NLP-36.2   (1) ----------------MVSLKQELQVLDYFGAFCVFISFFVIILIISVTC
Tm-NLP-36.1   (1) ----------------MVNLKQEVSIGDYFGAAIVFCLFFLIISVISFTC
Tm-NLP-36.2   (1) ----------------MVYLGQELSVLDYFSPLIVFGTFLTIVFIISATC
As-NLP-36.1   (1) ------------------MPQQEFEFIDYLGPLAVSICFAV-ALFLLSLV
As-NLP-36.2   (1) ----------------MVDLKQQLDVLDYFGPVAVWISFFAIVFIISVTC
Bm-NLP-36.1   (1) -------------------MKQIFGFTDYLGPLAASMCFFV-AIFLLSLI
Bm-NLP-36.2   (1) ------------------MPKQEFEFIDYLGPLAVSVCFVI-ALFILSAI
Bm-NLP-36.3   (1) ----------------MVSLKQEFEFYDYFGPVAVISLFAITILLISFFI
Bm-NLP-36.4   (1) ----------------MVDLKQQLDMLDYLGPIAVWISFFSITFIISVTC
Di-NLP-36.1   (1) ----------------MVSLKQEFEFYDYFGPLAVISIFAITVLLISFFI
Di-NLP-36.2   (1) ----------------MVDLKQQLDMLDYLGPIAVWISFFSIIFIISVTC
Na-NLP-36.1   (1) ---------------MSIDLKQNLDVIDYFGALVVWVIFFSILFIVSVTC
Na-NLP-36.2   (1) ------------------MAKQEFEPIDYFGPVVVAAIFAITLLLISFFV
Hc-NLP-36.1   (1) ------------------MAKQDFETIDYFGPVVVAAIFAVALFLISFFI
Hc-NLP-36.2   (1) ------------------MGKQTFTLLDYCGPLIVGAVFAS-ILFILSLM
Hc-NLP-36.3   (1) ---------------MSVDIKQQLDVIDYFGALVVWVIFFSVLFIVSVTC
Hc-NLP-36.4   (1) ---------------MSIDLKQHLDVIDYFGALVVWIIFFSFVFIVSVTV
Hc-NLP-36.5   (1) ------------------MAKQDFETIDYFGPVVVAAIFAVALFLISFFI
Bx-NLP-36.1   (1) ------------------MPKQDFLFTDYMGPVVAAVIFAICLLVVSFLL
Bx-NLP-36.2   (1) ------------------MPKQDFELIDYLGPVVVAIIFAICLLLISFTV
Bx-NLP-36.3   (1) MAETLKEDVRMSSDLLKKMPKQSLELIDYLGPIAAAVIFFSTMFILSVTV
Gp-NLP-36.1   (1) --------------------MGPTGTQGIGITKVLCRQFVLLVFLISFLF
Gp-NLP-36.2   (1) --------------------MGPTGTQGIGITKVLCRQFVLLVFLISFLF
Gp-NLP-36.3   (1) ------------------MPKQSLDLIDYFGPLAAWIVFFSIIFVVSVTC
                  51                                             100
 Ce-NLP-36   (36) N-FVCIKKDDDVTALERW--------------------------------
Ts-NLP-36.1  (35) ILWFCVDKTQEATVFDKWRG------------------------------
Ts-NLP-36.2  (35) ILWFCVSPTDDVTVFAKVRSLLTFL-------------------------
Tm-NLP-36.1  (35) ILWFCISKKDEATVVDKWRG------------------------------
Tm-NLP-36.2  (35) L-LCCCVEDVDSTVFSKFRR------------------------------
As-NLP-36.1  (32) INFIWITKKDDRTVFEKTERTHTE--------------------------
As-NLP-36.2  (35) ILWCCVSENDDSTVFAKWGM------------------------------
Bm-NLP-36.1  (31) INFIWISKDDERTIFEKFGSN-----------------------------
Bm-NLP-36.2  (32) INFIWITKNDDRTVFEKFGSTFDL--------------------------
Bm-NLP-36.3  (35) LNFFFISKHDEPTVFERIASKHNL--------------------------
Bm-NLP-36.4  (35) ILWCCVSETDDKTVFAKWGM------------------------------
Di-NLP-36.1  (35) LNFFFISKHDEPTVFERIASKHNL--------------------------
Di-NLP-36.2  (35) ILWCCVSETDDKTVFAKWGM------------------------------
Na-NLP-36.1  (36) INWCCIQKHDDITVLEEFSIRLPYGSCVSASAANSTILILEMKISLRKGS
Na-NLP-36.2  (33) INFFCITKYDDFTQFEKFGAKHN---------------------------
Hc-NLP-36.1  (33) INFFCITKYDDVTQFEKFGAKRN---------------------------
Hc-NLP-36.2  (32) MNFIFIRKRDEITSFEKLGAKYNL--------------------------
Hc-NLP-36.3  (36) INWCCIQKHDDITVLEEW--------------------------------
Hc-NLP-36.4  (36) INWCCIAKYDDVTVLEEW--------------------------------
Hc-NLP-36.5  (33) INFFCITKYDDVTQFEKFGAKRN---------------------------
Bx-NLP-36.1  (33) LNYCFVLKTDELTVFERFGSKHN---------------------------
Bx-NLP-36.2  (33) INWYCITHRDDLTVFEKLGRRADIR-------------------------
Bx-NLP-36.3  (51) ILWCFVTPNDDKTVFAKWGL------------------------------
Gp-NLP-36.1  (31) VNFFLITEEDDLTVFEKFGARHFG--------------------------
Gp-NLP-36.2  (31) VNFFLITEEDDLTVFEKFGARHFG--------------------------
Gp-NLP-36.3  (33) ILWCCVSETDDTTVFAKWGM------------------------------
                  101                                            150
 Ce-NLP-36   (53) ------------------------GYKKNIDMKLGPHRRSMVARQIPQTV
Ts-NLP-36.1  (55) ------------------------RRKGSVASRKSKHSMLISPTNY----
Ts-NLP-36.2  (60) ----------------------VLVVIVEQQLKKNLHNSICNNDSLLNAF
Tm-NLP-36.1  (55) ------------------------RRKESQTYPKS-KPLLAHGSNY----
Tm-NLP-36.2  (54) -------------------------SSRRRSYTMSSERTSSR--------
As-NLP-36.1  (56) -----------------------AAWGLRHPDAGFLLDSSENFRVIGFI-
As-NLP-36.2  (55) ------------------------GPRPRVSSQQQKYASGSPKAE-----
Bm-NLP-36.1  (52) -----------------------LCGVHRMRHRSKKGTKMMNKLMVNSYG
Bm-NLP-36.2  (56) -----------------------RCGVHRMRHRPNKSWKRVQLIDNQDV-
Bm-NLP-36.3  (59) -----------------------RLGPHSVKTVISRKQRYMKDNNSGDPD
Bm-NLP-36.4  (55) ------------------------GPRHQLTVAQQKYDSTSTHKAL----
Di-NLP-36.1  (59) -----------------------RLGPHSVKTVISRKQRYMKDSNSGDPD
Di-NLP-36.2  (55) ------------------------GPRHQLTVAQQKYDSTSTHKAL----
Na-NLP-36.1  (86) RADRKRVTTFTESIANVFDWFFEWGYNHKLHMRMGPHRPSVVARNVRHPL
Na-NLP-36.2  (56) ----------------------WRMGPHRLHIVKKGGFVADAYDDDDEEA
Hc-NLP-36.1  (56) ----------------------WRLGPHRMLHAKGGGLILKPEDDDDDTY
Hc-NLP-36.2  (56) -----------------------RVGPHRVSVVKRYVERPPLLEE-----
Hc-NLP-36.3  (54) ------------------------GYNHHVNMRLGPHRPSVVARNVRHSK
Hc-NLP-36.4  (54) ------------------------GYNHNLRMRMGPHRPSMVGRKLGHRT
Hc-NLP-36.5  (56) ----------------------WRLGPHRMLHAKGGGLILKPEDDDDDTY
Bx-NLP-36.1  (56) ----------------------VRLGPHSLDSIKRRSQRRLTEDEGCPIV
Bx-NLP-36.2  (58) ------------------------LGPHKMSVIRRGGYASTYAKDEEALM
Bx-NLP-36.3  (71) ------------------------GPQPK--NQQVVYRDGVASESLLVV-
Gp-NLP-36.1  (55) ----------------------VRLGPHSLAQIKSGGPSSVAAGQNVAPQ
Gp-NLP-36.2  (55) ----------------------VRLGPHSLAQIKSGGPSSVAAGQNVAPQ
Gp-NLP-36.3  (53) ------------------------GPQRRKRSGHHTLLNTPPQQQPKAE-
                  151                    176
 Ce-NLP-36   (79) VADH----------------------
Ts-NLP-36.1  (77) --------------------------
Ts-NLP-36.2  (88) LIVSSTPLFILLYYSVI---------
Tm-NLP-36.1  (76) --------------------------
Tm-NLP-36.2  (71) --------------------------
As-NLP-36.1  (82) --------------------------
As-NLP-36.2  (76) --------------------------
Bm-NLP-36.1  (79) PAVDTTVV------------------
Bm-NLP-36.2  (82) --------------------------
Bm-NLP-36.3  (86) GSVHESRKVVFTKEPEPEVHINSA--
Bm-NLP-36.4  (77) --------------------------
Di-NLP-36.1  (86) GSIQGSRKVVFTKEPEPEVHINSA--
Di-NLP-36.2  (77) --------------------------
Na-NLP-36.1 (136) KEDNAL--------------------
Na-NLP-36.2  (84) AYFA----------------------
Hc-NLP-36.1  (84) YA------------------------
Hc-NLP-36.2  (78) --------------------------
Hc-NLP-36.3  (80) VESDSL--------------------
Hc-NLP-36.4  (80) STTIPGHTGHP---------------
Hc-NLP-36.5  (84) YA------------------------
Bx-NLP-36.1  (84) TENTKVPIPQVQVDAASTS-------
Bx-NLP-36.2  (84) KKQSHAAQVALASEIA----------
Bx-NLP-36.3  (94) --------------------------
Gp-NLP-36.1  (83) KEQRKAQKDEKKAEEFEMKEHRTNVF
Gp-NLP-36.2  (83) KEQRKAQKDEKKAEEFEMKEHRTNVF
Gp-NLP-36.3  (78) --------------------------
```

NLP-37

```
                 1                                               50
Ce-NLP-37    (1) ---------------------------MSSRISVSLLLLAVVATMFFTAN
As-NLP-37    (1) ---------------------------MKFTALPALLAFAIIAVNANSEE
Bm-NLP-37    (1) --------------------------MKYFYCIVFFLPIFCCFNIANNKK
Di-NLP-37    (1) --------------------------------------------------
Na-NLP-37    (1) ------------------------------MVSRAVAPTALLLAILATTL
Hc-NLP-37    (1) MHIIWRWCVENRMRPSSITTLCPYRFHPLQTQMISRALFTVLFAVSAISS
Bx-NLP-37    (1) ----------MHAIIDYFRAVPSLLGKMQSLVSSFVVLMTITVTVMTSKS
                 51                                             100
Ce-NLP-37   (24) VVDATPRSQGNMMRYGNSLPAYAPHV-------------------LYRFY
As-NLP-37   (24) RRSLPAMGNRVIAQFLGRLPLYQGSSN------------KRFDAVMMPFD
Bm-NLP-37   (25) LLMLYPREQVMERLLESLPKRDDEHV-------------------VNTRN
Di-NLP-37    (1) ----------------IFFLKFTKRK-------------------LQIRN
Na-NLP-37   (21) AQQERIMGAMPSRYTFYHDRGYEYAP-------------------PLTLY
Hc-NLP-37   (51) ANQQRRMETTPSGYNSYQDKPYSRT--------------------LLSLY
Bx-NLP-37   (41) DALPFPPHTSLVIQDTQLIRQFLDRYQPAVLFHPMVRRYDVQQEDMDDGE
                 101                              136
Ce-NLP-37   (55) NSRQFAPINKRNNAEVVNHILKNFGALDRLGDVGK-
As-NLP-37   (62) TSSDDYTVQKRNNAEVVNHILKNFGALDRLGDVGRR
Bm-NLP-37   (56) DGISAYMVRKRNNAELVNHILKNLNEVDLLGEVGR-
Di-NLP-37   (16) DKISAYTMRKRNNAELVNHILKNLDEVDLLGDVGR-
Na-NLP-37   (52) RKVGARPVSKRNNAEVVNHILKNFGTLDRLGDVGK-
Hc-NLP-37   (81) RKLHDGLISKRNNAEVVNHILKNFGTLDRLGDVGK-
Bx-NLP-37   (91) ERLALRPMAKRNNAEVVNHILKNFGTLDRLGDVGK-
```

NLP-38

```
                 1                                               50
Ce-NLP-38    (1) --------------------------------------------------
As-NLP-38    (1) TNERRKRLYAYNIPYRTSYHDRREKWTRYQCSEAEYHSMMSSHSSFARHM
Bx-NLP-38    (1) --------------------------------------------------
Gp-NLP-38    (1) ----------------------------MSTQNSKSSAPVFSTKHCALGH
                 51                                             100
Ce-NLP-38    (1) -MQLIHFIVGLAMLISLSLAASDDRVLGWNKAHGLWGKR-----------
As-NLP-38   (51) LRTGIVLIATIVFLSAPALAREQDKRDNWNKALGLWGKRSI---------
Bx-NLP-38    (1) --MVQKSAWQMLILLLLAVFSTIQADRQWSAGVGLWGKRS----------
Gp-NLP-38   (23) RFALLVALLLFGLVTISVGAPADGQSHRWSNGVGLWGKRSDGLTNLALLD
                 101                                            150
Ce-NLP-38   (39) --SVQEASQDKRTPQNWNKLNSLWGKRSASSFDDDYTTENGDDDVTMLYK
As-NLP-38   (92) SPQFLDTNLEKRSQDQWNKLNSLWGKR------TSWQTANG------LWG
Bx-NLP-38   (39) ----YPLLRKRSPDFNPNGEAGLWDKR-----------------------
Gp-NLP-38   (73) GLFDEARSVRKRPEHSWHKLNALWGKR----------SSNS---------
                 151                                         197
Ce-NLP-38   (87) RSNLSPRFLGRMTFARIPKISPAQWQRANGLWGR-------------
As-NLP-38  (130) KRS-----------------S---WETANGLWGKRSLKSEQLDDH--
Bx-NLP-38   (62) ----------------------AQWQAANGLWGKRSVPPVPQYVPWN
Gp-NLP-38  (104) --N---------------------WQTANGLWGKR------------
```

NLP-40

```
                  1                                               50
 Ce-NLP-40    (1) --------------------------------------------------
 As-NLP-40    (1) --------------------------------------------------
 Na-NLP-40    (1) --------------------------------------------------
 Hc-NLP-40    (1) --------------------------------------------------
 Bx-NLP-40    (1) --------------------------------------------------
Gp-NLP-40.1   (1) --------------------------------------------------
Gp-NLP-40.2   (1) MASSSAYPILFFTQTWSGDDKLWIFYWLSSESKFGIASFPDTVYEMEADR
                  51                                             100
 Ce-NLP-40    (1) --------------------------------------------------
 As-NLP-40    (1) --------------------------------------------------
 Na-NLP-40    (1) -----------------------------------------MSLSAFQTR
 Hc-NLP-40    (1) --------------------------------------------------
 Bx-NLP-40    (1) --------------------------------------------------
Gp-NLP-40.1   (1) --------------------------------------------------
Gp-NLP-40.2  (51) DTRKAFLGNIAKSLVEEYWETMFEEGKTKIKQMNDESATVSYAFKHFLEL
                  101                                            150
 Ce-NLP-40    (1) --------------------------------------------MKLVIL
 As-NLP-40    (1) ------------------------------------------MYRMMRFT
 Na-NLP-40   (10) SASLKHFDKKACMDELVITKLFEWAEPHQLHTDTQHTKRTHTAIAPMRAL
 Hc-NLP-40    (1) ------------------MWAEHGKTSVLIRPRGWHTGHSRSPTSPMRPL
 Bx-NLP-40    (1) ----------------------------------------------MKSV
Gp-NLP-40.1   (1) ------------------MGHRRCCTACTCGAGVCMVNNNDEFCPKSSAS
Gp-NLP-40.2 (101) PQLDDPVKRRQRIDRLVKFFVDEALEKKRMEQGKQPLSLHNRNHPTNASS
                  151                                            200
 Ce-NLP-40    (7) LSFVATVAVFAAPSAPAG-LEEKLRALQEQLYSLE---KENGVDVK----
 As-NLP-40    (9) LFAVLLPAVLQAAAQPLSELEGKFKSLQQQLLNLQ---QEMQLQQQQDKS
 Na-NLP-40   (60) IIAVSIGLIFAAPIQDADSVQEKLKALQEQLNTLQ---KQLVVPAS----
 Hc-NLP-40   (33) VIALIFAAAYAAPVQDGEKIEEKLKHLQEDLAALQ---RQLTAPTT----
 Bx-NLP-40    (5) LLVFGVVLGVVLAAPAGEDLQKKIDAMEKHIQLLE---KALLQRRSPLQG
Gp-NLP-40.1  (33) RCGAAQSNSSNGRQDQPQQQINALRAQMDQLQYELGQLRQYAPAGADWME
Gp-NLP-40.2 (151) LRNNSAKRKQPTNTRKALRKLPTLGLNFDDVIEED---QQRGLQIADWME
                  201                                            250
 Ce-NLP-40   (49) -----QKEQP------------AAADTFLGFVP------QKRMVAWQPMK
 As-NLP-40   (56) VAVPAQEQQPQIHDEYMLPELMAQGIQYGDLRAKPSNTQQKRLVYWQPMK
 Na-NLP-40  (103) -----SQNED------------ARKFLFNDWQP------QKRMVAWQPMK
 Hc-NLP-40   (76) -----MEKEE------------ARKLLYNDWQP------QKRMVAWQPMK
 Bx-NLP-40   (52) SLIEGNQIMN------------PRATRALAFQP------MKRMVAWQPMK
Gp-NLP-40.1  (83) PVWPFSNNVDYHPKSMLVRRHMAWQPMRRGIENDV---------------
Gp-NLP-40.2 (198) PVWPFSNNVDYHPKSMLVRRHMAWQPMRRGIEKDER-SMTQRHIAWQPMK
                  251                                            300
 Ce-NLP-40   (76) RSM------------------------INEDSRAPLLHAIEARLAEVLRA
 As-NLP-40  (106) KS--------------------------VVSTKEQVLQAIESRLAEVIHA
 Na-NLP-40  (130) R---------------------------NLDERQPLLHAIEARLSEVLRA
 Hc-NLP-40  (103) R---------------------------NLDERQPLLHAIEARLSEVLRA
 Bx-NLP-40   (84) RSI------------------------AAEYNKDQVIRAIEEQLLEILHA
Gp-NLP-40.1 (118) --------------------------------------------------
Gp-NLP-40.2 (247) RQQQPEHGMGEQYDGEGPDVAAATVYAAGDGQKQQVMRTVEQELIEVLRA
                  301                        330
 Ce-NLP-40  (102) GERLGVNPEEVLADLRARNQFQ--------
 As-NLP-40  (130) GEKLGVSADELLAHLKMRNSSPQ-------
 Na-NLP-40  (153) GERLGVSPEEVLQDLRYRNQFKK-------
 Hc-NLP-40  (126) GERLGISPEEVLQDLRYRNQFKK-------
 Bx-NLP-40  (110) GETLGVNAEEVLGDLKKKNGDLM-------
Gp-NLP-40.1 (118) ------------------------------
Gp-NLP-40.2 (297) GERLGVSADEILAHLRARNNNNNAGGLGRR
```

NLP-42

```
                 1                                               50
Ce-NLP-42    (1) ------------------------------------------MRVQVVTL
As-NLP-42    (1) --------------------------------------------------
Na-NLP-42    (1) -----------------------------------------------MRS
Hc-NLP-42    (1) -------MQLTLAYGMKASDYLQGMPKGDIKPGWEDLGWAWGKRINMKGS
Bx-NLP-42    (1) -----------------------------------------------MSL
Gp-NLP-42    (1) MLRQCFLRPSCPFVGGGSSLLLLALLALPSAFAWNSRTFGGLDPNSGGGG
                 51                                             100
Ce-NLP-42    (9) LAVLLAVLQFTSAAG--NYYSGYPSDRTMKRSALLQPENNPEWNQLGWAW
As-NLP-42    (1) MCRRLHRLSVAFTSFIVIAFSHEALSLPSSAREREMKRNKPHWDDLGWSW
Na-NLP-42    (4) LQVIVLLIAIIQ------ATFTYNLKLADALQSMPKREIKPDWEDLGWAW
Hc-NLP-42   (44) SSVFLLFIAFMQ------LTLAYGMKASDYLQGMPKGDIKPGWEDLGWGK
Bx-NLP-42    (4) LTRILCTLTVVG------ITIAITHASPMDRSALKKASDNPQWEDLGWAW
Gp-NLP-42   (51) GTRRLYMLKKSESPQWEDLGWAWGKRSGGPNSAAPSPQNSANWDDLGWAW
                 101                                            150
Ce-NLP-42   (57) G---------------------KRSAGMEIP-HRAARALHPVKK-NPDWQ
As-NLP-42   (51) GKRALGGSVSPQWESSGWTWGEQTFPLQESHTQRALRVAMTKKN--PDWH
Na-NLP-42   (48) G---------------------KRSVPDSVYPQRFLRPMQLVKK-TPDWH
Hc-NLP-42   (88) ----------------------RSAVGELSYPRRFLRSMQLVKK-SVDWH
Bx-NLP-42   (48) GKRSVA---------------VELVDPEDVMLQRYVRSMKAIKK-NPDWH
Gp-NLP-42  (101) GKRSPAEGGEAARR-------AKAHREDPIVSMMSNRVVRSMTKQSPDWH
                 151
Ce-NLP-42   (84) DLGFAWGRK-
As-NLP-42   (99) DLGWTWGRK-
Na-NLP-42   (76) DLGWTWGRRK
Hc-NLP-42  (115) SLGWAWGRRK
Bx-NLP-42   (82) DLGWAWGK--
Gp-NLP-42  (144) DLGWAWGRRK
```

NLP-43

```
                 1                                               50
Ce-NLP-43    (1) ------MSLAQSTFYLLFVAFLAVVIAVTADKQQSAYDLETPISAYKRFY
Ts-NLP-43    (1) MSSVKSCICIFTAFCLLASFTSATLVDSPSSAKEVQMIPVFYSDTVGDHH
Tm-NLP-43    (1) ---------------MEIQSGFGESPKLPQGAQSWDMDKEFLQNLGPSAL
As-NLP-43    (1) ----------MHVQCAAAIFLLITVQLLISTVSAQDTPLVFKANQMKRFY
Na-NLP-43    (1) ----------MQRLLLVIATIFSVLANCQGEQQHSVENRYFTDDGITGDY
Hc-NLP-43    (1) --------------------MLAVLVSCQREQLSPEMSRYFMDDIGSSEY
                 51                                             100
Ce-NLP-43   (45) SWEDAKRAASSEEG----MRNKRKQFYAWAGKRSSAPVHYFEDSIAQEAG
Ts-NLP-43   (51) VIDPTVDNLEDVSELN---NLMKRKFYAWAGKRASRFYQWAG---KRNGE
Tm-NLP-43   (36) GTIRRPAYPDDPEDQMGHYQLLKRKFYAWAGKRSPAPRFYQWGGKRSDDV
As-NLP-43   (41) SWEETKRSPSEDVQPP--SDYTKRKFYAWAGKRS----------APISDS
Na-NLP-43   (41) IPYDMKGFVT-----------KRKQFYAWAGKRS---------------V
Hc-NLP-43   (31) VPSLMKGSDS-----------KRKQFYAWAGKRS---------------H
                 101                                            150
Ce-NLP-43   (91) PSMEKRKQFYAWAGK-----------------------------------
Ts-NLP-43   (95) KVEKVRRKFYPWAAKCNKKTYEVAKDIQNRTTILFIFAITYLYCTLTVQN
Tm-NLP-43   (86) SDEKVRRKFYAWAG------------------------------------
As-NLP-43   (79) VAYNKRK-FYQWAGK-----------------------------------
Na-NLP-43   (65) PVIDKRKQFYAWAG------------------------------------
Hc-NLP-43   (55) PPYEKRKQFYAWAGK-----------------------------------
                 151     161
Ce-NLP-43  (106) -----------
Ts-NLP-43  (145) LQKACKILRIR
Tm-NLP-43  (100) -----------
As-NLP-43   (93) -----------
Na-NLP-43   (79) -----------
Hc-NLP-43   (70) -----------
```

NLP-44

```
                 1                                               50
Ce-NLP-44    (1) ----------MLLWIVATLLIFSLPVSTALDYNDFSLQRIARAPHPSSAL
As-NLP-44    (1) --------------MATLAVTFFALLLTVFVSGNYPKQRRNDRPQSSAAV
Na-NLP-44    (1) MEALLDQDSRQTQEERGMKNRSTPTTQSAENLDVFPLQRISRSQLALSTL
Hc-NLP-44    (1) ---------MKLTLTLLLVIIFFRVNAEHSDVDMFPLHRISRSTLPLSSL
Bx-NLP-44    (1) --------MKNAAVAVLLCLCLVYVSSQELFSERFAVPKPIGVPVEASQL
Gp-NLP-44    (1) ---------------MSVVAGRKVLIDPYEWPRVLAARKAQRPPTDSWNY
                 51                                             100
Ce-NLP-44   (41) LVP----YPRVGKR-SNILNNNSESQNS-----VQKRLYMARVGKRA-FF
As-NLP-44   (37) LVP----YPRVGKRFDTIAATYPYSSDE-----NSQSSSSPRAKQPV-RL
Na-NLP-44   (51) LVP----YPRVGKRSTNGTDHQVSAPIL-----HAKRLFAT-VGKRT-YM
Hc-NLP-44   (42) LVP----YPRVGKRSFFTGDRNVYPPTS-----IRKRLYVARVGKRR-YL
Bx-NLP-44   (43) RSRGSDVYVTPGKYPSRVYGLFKRAADQPAPEFQVKRMYVARIGKR--QV
Gp-NLP-44   (36) DPPWYDDIFTRGEGMHKRMSSMDGGGGEGEDGSMGKRLFVARIGKRMAHG
                 101                            134
Ce-NLP-44   (80) YTPRIGK---------------------------
As-NLP-44   (77) EARSPHRWVGGGCTYPFTDLSSVHNAAPLNVASD
Na-NLP-44   (90) YAARVGK---------------------------
Hc-NLP-44   (82) YTARVGK---------------------------
Bx-NLP-44   (91) YRARVGK---------------------------
Gp-NLP-44   (86) YWARVG----------------------------
```

NLP-46

```
                 1                                               50
Ce-NLP-46    (1) -----------------------------MLSIRTFVLVLLVLVGLAAAL
As-NLP-46    (1) -------MQYGYSIKVLLTTTLIALVISLPVHDADYSSKDQYRALEETPI
Bm-NLP-46    (1) -----------------MVDVSYTLVLLLATFIKFAQGLPIFHETNYEPF
Na-NLP-46    (1) ------MLLVPYKDYGATESAKTITVKSYMTTVIGYVRSLVFFLLIGLSL
Bx-NLP-46    (1) MEKCPMVDDKVRADFGALASAAYCWVKIRRKFVWTMSQALEAALLCVVLI
                 51                                             100
Ce-NLP-46   (22) PYFRLNYDYDP------------------------VDLDNDEKNAARQFL
As-NLP-46   (44) LYPANILFLRG-----------------------------DEYADFISPR
Bm-NLP-46   (34) ANYLNSLDFDQP------------------------KLSMYEPLPLPILD
Na-NLP-46   (45) ALPLLWLDVEG------------------------GDDQQYQQWRGIGPL
Bx-NLP-46   (51) SVHSLPILMNGQPMEMMYIDGPHDMFLPALDTEGPESAWSWFDQTQSSHP
                 101             119
Ce-NLP-46   (48) PYMKRNIAIGRGDGLRPGK
As-NLP-46   (65) AG-KRNIAIGRGDGFRPGK
Bm-NLP-46   (60) VMVKKNIAIGRADGFRPGK
Na-NLP-46   (71) TMVKRNIAIGRGDGLRPGK
Bx-NLP-46  (101) VAQKRNIAIGRGDGFRPGK
```

NLP-47

```
                  1                                               50
 Ce-NLP-47    (1) ----------------MQLYVVLCFLVLLGLSAGQMTFTDQWTKKR---A
 Ts-NLP-47    (1) -------MQFIFKEAVTVSWCTLLLAFLVSQTASQLTFSSGWGKRTTAEP
 Tm-NLP-47    (1) -------------MSHTSLRVTILLVFVICGAVAQLTFSQGWGKRTTDAL
 As-NLP-47    (1) ------------MFLTRHNLILIAIVELLRMGTAQITFTDQWTKRS-GIS
 Bm-NLP-47    (1) ------------------MRRITLLPAIIYIVTAQMTFTDNWDKRSLSNL
 Di-NLP-47    (1) ------------------MRRVVLLLTVIYIVTAQMTFTDNWDKRTFPSL
Na-NLP-47.1   (1) ---------------MRSVLVSLIVLLLVPSSFSQMTFTDQWSKK--SEP
Na-NLP-47.2   (1) ------------------MRALLVLLAFLALTFAQMTFTDQWNKRSAPYE
 Hc-NLP-47    (1) MAQNHSIQRRKHGSNMRFVLVTIIVALCIGISLSQMTFSDHWSKKSLPGP
 Bx-NLP-47    (1) ------MDLVCRSLTNTMQTIMILLLVSVTMTSAQMTFSDGWGKRSSPLK
 Gp-NLP-47    (1) -------------MPSLVFPTLCFFVSAVVLANSQMTFSDGWGKRS-AVA
                  51                                             100
 Ce-NLP-47   (32) TLHKQLPVVTPE------------------------EPICPSDRVQAVFE
 Ts-NLP-47   (44) IGKASYCAHLDQD----------------------ILQLIQQIITVHISF
 Tm-NLP-47   (38) ILKNANLDCESIN---------------------EEAMKIVRRLVEVSFF
 As-NLP-47   (38) PIPLLKDLSQRG------------FASEAIPNATAFHMSFHAEYTANCSI
 Bm-NLP-47   (33) FRGWQQDDLMVN-------------------------RACSWDTIAEITR
 Di-NLP-47   (33) FRGWQRDELTIN-------------------------EACSSDVIIEIIK
Na-NLP-47.1  (34) PFPRYKPLLRQD--------------------------------------
Na-NLP-47.2  (33) PLASLKTTGRSS--------------------------MCGHRAIDEMLL
 Hc-NLP-47   (51) AFPRYKPRESESSSQQKPAQ---------------QQQFCQEEKVTDALA
 Bx-NLP-47   (45) PFSAGDDSGVNS-------------------------NPLLDKCNTEYSQ
 Gp-NLP-47   (37) PFALLADRHMTEPTKRQQKSKTNANFFVSPDEDTLTMSSEMEACHAGYAQ
                  101                             135
 Ce-NLP-47   (58) QLDQLQKAQQRLTEYLASCAYPVEVPQKAEKM---
 Ts-NLP-47   (72) ENFFIQFIYSYFSFLFILFSFSIRNITFNIKNA--
 Tm-NLP-47   (67) RGIILLCMQLMLQDYRRSYRICQQQHMRRLFGRK-
 As-NLP-47   (76) TLESSFITQIFIDQQVMRMKDAKRPFSSQWWLS--
 Bm-NLP-47   (58) ELKQIYKKQNELINLIDRCSIIERPVFVKEG----
 Di-NLP-47   (58) ELKHIYKRQNELINVIDRCSIINRPVFIKEGKQKF
Na-NLP-47.1  (46) -----------------------------------
Na-NLP-47.2  (57) QLVQLQSAEKEIIAELNACLPAKRQ----------
 Hc-NLP-47   (86) QLAQLGEAQNMLTSYLNTCYQSRPRV---------
 Bx-NLP-47   (70) TLVQLHHRIMKLYDTYEQCQTRALNGVATHRQQ--
 Gp-NLP-47   (87) RLIQLHEQMLSLFSTYQQCQTKAAIETKKSMRQS-
```

NLP-48

```
                 1                                               50
Ce-NLP-48    (1) MMSSRRIVYTLLLITIALCAYSTAYAPRNIRGVGDVPSMFFSPFRMMGKR
Na-NLP-48    (1) MSRSRVLVFTFMALFILAAVVDSATIVR-----RSIPEVFNSPFHVLGKR
Hc-NLP-48    (1) MLALRVFVLTIMLAMVFNVIAESDRVIR----SGPLYNAFNVPFRVIGKR

                 51                                             100
Ce-NLP-48   (51) NQFYGLFKKIRSNEVVDY--------------------------------
Na-NLP-48   (46) NQFYGLFKPFCERRVNDFYKKKKLLSKLFEVNYSQRMSCTHGLLGVLVMT
Hc-NLP-48   (47) NQFYGLFKKLNYESSKPF--------------------------------

                 101                         131
Ce-NLP-48   (69) -------------------------------
Na-NLP-48   (96) FFFLNITLTPLGAVPGTAPGVACGGLLMTRT
Hc-NLP-48   (65) -------------------------------
```

NLP-49

```
                 1                                               50
Ce-NLP-49    (1) ------------------------------MMKWLLLAVFCIA----AYA
Ts-NLP-49    (1) MGNNQRILSANDPKTEDIHWLNSNQLAELQMNITVIALVLTTLCVIVVHS
Tm-NLP-49    (1) --------------------------MTLFKNLALLSLLLSTLYTSSVLS
As-NLP-49    (1) ---------------------------------------------MQFYR
Bm-NLP-49    (1) ------------------------------MKLSVTLLIVLVTDCIAWPK
Di-NLP-49    (1) ------------------------------MKLSIALLIVFITDCIAWPK
Na-NLP-49    (1) ------------------------------MMKWTLLSVLALVCILNAFA
Hc-NLP-49    (1) ------------------------------MMKWTMFAVLVLACIMNAFA
Bx-NLP-49    (1) ---------------------------MMKLWVSVLALLAIVSAVLAYPQ
Gp-NLP-49    (1) -----------------------VAPMRRSLPSFLPFRFLLLPLLFLIVS
                 51                                             100
Ce-NLP-49   (17) WADGTEDSNIDQLMSRVYRTVLLKSKRS----------------------
Ts-NLP-49   (51) EPNLYDWYQNSGSLERFLQELREKRKRSRISKYSQNQESIRHRGNVLSST
Tm-NLP-49   (25) QRNPYEWIQPSQSFEKMIEQLKEKRK------------------------
As-NLP-49    (6) KTTLKHNDDEEAKRASCYSVSTFWLSHIFIA-------------------
Bm-NLP-49   (21) SIDRTGNDLPIEQLSRIYQAMLLKSKRS----------------------
Di-NLP-49   (21) NIDRTDNDLPIEQLSRAYQPILLKSKRS----------------------
Na-NLP-49   (21) WADASEDSNMDQLMARVYRTVFLKSKRS----------------------
Hc-NLP-49   (21) WADGVEDANIDQLMSRVYRTVLLKSKRS----------------------
Bx-NLP-49   (24) TPSEFDAEFARQLIGRAYSALAKTKRSRE---------------------
Gp-NLP-49   (28) NSAAPTIDSDLIPPARFYRTLWSFNPKRS---------------------
                 101                                      144
Ce-NLP-49   (45) ------PSMGLSLAEYMASP-QGGDNFHFMP-SGRK--------
Ts-NLP-49  (101) VHLERLPSIGLSLAEYMADP-ERSQDFHFLR-GRKKRMIQFFNS
Tm-NLP-49   (51) ------PSIGLSLAEYVADP-TRADNFHFLN-GRRKRSTESGS-
As-NLP-49   (37) ------ASMGLSLAEYMASP-QGQDNFHFIP-SGRK--------
Bm-NLP-49   (49) ------PSMGLSLAEYMASP-QSQDNFHFIP-IIRR--------
Di-NLP-49   (49) ------PSMGLSLAEYMASP-QGQDNFHFIP-IIRR--------
Na-NLP-49   (49) ------PSMGLSLAEYMANP-QGGDNFHFIP-SGRK--------
Hc-NLP-49   (49) ------PSMGLSLAEYMASP-QGGDNFHFIP-SGRK--------
Bx-NLP-49   (53) ----FKTSMGLSLAEYMASP-QGGDNFHFIP-SGRK--------
Gp-NLP-49   (57) ------PSIGLSLAEYMAAGNQGQEQFHFIPGSGRRK-------
```

NLP-50

```
                1                                               50
Ce-NLP-50   (1) --MRFSVFVALFALLAVSYGFMYEMED-SYPVSGELPKRSLENFWRNIHL
Na-NLP-50   (1) MRALRELLVLLLAILATTQAFMIDGDEYSGLVDYPLPKRNFQHIWRNIQM
                51                       77
Ce-NLP-50  (48) QMPSGRGGQKRTEGLSRASANAYYRLG
Na-NLP-50  (51) QMP---GSQKRTESLSRARANAYYRLG
```

NLP-51

```
                1                                               50
Ce-NLP-51   (1) ------------MRFLILALLVLFAITQAYPSSDYQPRYRKS------QT
As-NLP-51   (1) MRMPQVRIIAVSLVLLVLVDALPSLQYPYYSIQQLQPRYRKSS--SANQP
Hc-NLP-51   (1) ------------MRILLFCLALLLSVCGFH--SEAQTRLRKS------HA
Bx-NLP-51   (1) MSRLTVLLLIAFMCTMLMVSVRPSPIHQQPLNTDVYLRFRRNAQNSRPTR
                51                                   89
Ce-NLP-51  (33) QEANIQPFIRFRKSTQPQHNWMFRPDVAPYFE-------
As-NLP-51  (49) HDSMVQPFIRFRKALSNGFHSWFDEVTPSLAYLYAESRN
Hc-NLP-51  (31) QDANIQPFIRFRKASYGSWIDPLNAYH------------
Bx-NLP-51  (51) ADEYVQPFIRFRKNYQLPYDVSFDYVPYMTRI-------
```

NLP-52

```
                1                                               50
Ce-NLP-52   (1) MFRTLLICFFLMAAVSITSAYILDYDVPERQTRGDVKSVFFSPFRMVGKR
Na-NLP-52   (1) ----MSRTSFVIIVAFICLVLIAHVQSRNLAMTGGVSSLFNSPFRVIGKR
                51                      76
Ce-NLP-52  (51) SQFYGLYKQLGHSKRETSLE------
Na-NLP-52  (47) SQLYGYYRQGNLDFPQVPLAIDQFDY
```

NLP-54

```
                 1                                               50
Ce-NLP-54    (1) MMNYSVVLLLFALCSIVAASPMWFDE---EEPMNLRAFRVMPNQLDSIDA
As-NLP-54    (1) ---MNNKARCNDITGRELFGKRSLDASTNEYDALDAVHSVEGSTSISSSY
Na-NLP-54    (1) MQLLLLFTMVALIVTVVSAEPRLRFSNSAESGLLRRAGLFFRKRGSAWDG
Hc-NLP-54    (1) -MNLLLLLCTLGLVALTAAEPRLRLS-SAEEDLVRRAKVLLKKKGNMWGT
Bx-NLP-54    (1) MHAIISFALFATIAAGMSAQDMELNGDFSQHDYRMLPVQREMMRENVGET
Gp-NLP-54    (1) MPILFVLIGAKIVIGGLDVEEWDFPTDRLEAFQKRKVRELFGKR----ST
                 51                                             100
Ce-NLP-54   (48) SRRLLKRASMFN---------------KRRGRELFGKRSLPIEYP-----
As-NLP-54   (48) TTPSEQQQRLFEFVPSMADDAHG----AGINLEVNRERMPAVEFLKLLAA
Na-NLP-54   (51) VSAPFTFTNLAE---------------KRRVRELFGKRSAPAVETPDVNV
Hc-NLP-54   (49) PSKSYGYTNLAE---------------KRRVRELFGKRSAISATPENE--
Bx-NLP-54   (51) LPTLHFLRSIRRSRELFGKRSPSVVEEKRRSRELFGKRSAKSEE------
Gp-NLP-54   (47) PPAAIPYERHSR-----------------RTRELFGKRADPFAESNDEPL
                 101                                            150
Ce-NLP-54   (78) -------------------------------------------DETVMYE
As-NLP-54   (94) TSGPK----------------------------------IVLQNRAALGM
Na-NLP-54   (86) LELPD-------------------------------DQYPSLYDSMLELH
Hc-NLP-54   (82) -ALSD-------------------------------EQYQPVDDLLMELH
Bx-NLP-54   (95) -EIPF-------------------------------NSQEVLALATLLTE
Gp-NLP-54   (80) MAVDLPEEDEHRIKMPRTVFARLRTFQNRANAVEEEQEAPFSTEGLWLLQ
                 151       163
Ce-NLP-54   (85) SRFRRRANELFG-
As-NLP-54  (110) RHRRGRSRELFG-
Na-NLP-54  (105) RDRRGRVRELFG-
Hc-NLP-54  (100) RDRRGRVRELFG-
Bx-NLP-54  (113) RPRRARGTELFGK
Gp-NLP-54  (130) RPRRGVHRELFG-
```

NLP-55

```
                 1                                               50
Ce-NLP-55    (1) --------------------------------------------------
As-NLP-55    (1) --------------------------------------------------
Na-NLP-55    (1) MTTNELATCDASHSALGHYSNQDFLSMEPVYLLVYHDVKKKRCNYVVETD
Hc-NLP-55    (1) --------------------------------------------------
Bx-NLP-55    (1) --------------------------------------------------
Gp-NLP-55    (1) --------------------------------------------------
                 51                                             100
Ce-NLP-55    (1) --------------------------------------------------
As-NLP-55    (1) -----------------------------------------MKAPWIIYI
Na-NLP-55   (51) EERISTRITRDHTEARQRLVLRWYCVKSCVRFGCGSQGVELIPGSAVYGT
Hc-NLP-55    (1) --------------------------------------------------
Bx-NLP-55    (1) ------------------------------------MVSVSVIILQLITA
Gp-NLP-55    (1) -------------------MATLSEDFRRHNQCSIRNMHRTWLLLLCATV
                 101                                            150
Ce-NLP-55    (1) --------MSCISMLFILLVACLLVSNAMYINPDYY---YVEQLPTMKKS
As-NLP-55   (10) TVLAISTGRINGHLMERFMGNRANLNDATFILPDDS-PSSSHSILFTKKS
Na-NLP-55  (101) APHAMNGSLSAIIVRVLLLCALVALSSGYYLAYKPS-----KNGLLPIKK
Hc-NLP-55    (1) ----MNSSLSAAIVRVLLLCALVALCNSYYLAYEPNGASGYEGAWRAAKK
Bx-NLP-55   (15) FLLVVDGYIIDRWSTLEDPARLYPLSDSEYIRYPVRRSVDTEEASMDRTF
Gp-NLP-55   (32) FFMSIASNQVHGYLIDGRSLQKMMAVQRRWAQRPAKLEEFSKFVPASKKS
                 151                                            200
Ce-NLP-55   (40) GQLRA---------------LAGSRNCFFSPVNCIITHDINSYRRLAKGS
As-NLP-55   (59) LRQFQ---------------GSGSRNCFFSPIQCVLQHDLSKYRKLVDST
Na-NLP-55  (146) SQIGG---------------LVGSRNCYFSPVSCVITRDVSAYRKLAGPS
Hc-NLP-55   (47) SQLRA---------------LAGSRNCFFSPINCVITHDVSTYRKLAGTR
Bx-NLP-55   (65) SQGYAKKSQNLSYKALKKFSGSGSRNCFFTPIQCMIQHDMSKYKKLVDGS
Gp-NLP-55   (82) AGYGVLSKFG----------GSGARNCYFTPVSCMIQHDMSKYKKLIESS
                 201
Ce-NLP-55   (75) SYA------
As-NLP-55   (94) NYKLNV---
Na-NLP-55  (181) ---------
Hc-NLP-55   (82) ---------
Bx-NLP-55  (115) KVIQN----
Gp-NLP-55  (122) RLDVFANLR
```

NLP-56

```
                  1                                                  53
Ce-NLP-56A    (1) -------------------------MPSPSSLLGSLLLVCAVLTITSRASS--
Ce-NLP-56B    (1) -------------------------MPSPSSLLGSLLLVCAVLTITSRASS--
Ce-NLP-56C    (1) -------------------------MPSPSSLLGSLLLVCAVLTITSRASS--
 As-NLP-56    (1) MPGKLKEDFRERPEVLSTSYLSAFLFLRLHYKYHLIKLLLSYVHVVEDLMESG
 Na-NLP-56    (1) ---MFLAEKPQGVDIRDANQKKFEAKGKYVMRLWLNTLLAAMLLVIVTALEP-
 Bx-NLP-56    (1) -------------------------MCSRFSWALLIVGMALAIHLASGVPSP-
 Gp-NLP-56    (1) ---------------------------TEMAEGYVISSYQTKLLPPKGFFPS-
                  54                                                106
Ce-NLP-56A   (27) -----------------IMTDDVEPPQLLTRQLRSFPYSVSFYRMLG------
Ce-NLP-56B   (27) -----------------IMTDDVEPPQLLTRQLRSFPYSVSFYRMLG------
Ce-NLP-56C   (27) -----------------IMTDDVEPPQLLTRQLRSFPYSVSFYRMLG------
 As-NLP-56   (54) VMYWLGLCSVLLWLHASVFSTVQSSEDINFRTARGFPQTTSFVQLLQPDEG--
 Na-NLP-56   (50) ----------------------FEDEFLKERPTRAFPFLLRITPLIG------
 Bx-NLP-56   (28) ----------------KAKSDKQKLFRVARMTFQSFPYSLAYMRMMRPAPSSI
 Gp-NLP-56   (26) ----------------STIAYYRSMKTLLQRPNKIVPNHVGDSEGHRP-----
                  107                                               159
Ce-NLP-56A   (57) ---HDRQLRPYYGVNDEVAALIDSMNSDNVANEDVFPTRPR-RSDGLRGSFYW
Ce-NLP-56B   (57) ---HDRQLRPYYGVNDEVAALIDSMNSDNVANEDVFPTRPR-RSDGLRGYACR
Ce-NLP-56C   (57) ---HDRQLRPYYGVNDEVAALIDSMNSDNVANEDVFPTRPR-RSDGLRGSFYW
 As-NLP-56  (105) RKWRSYNGAVAVRLSSDLNDLINQIEAEQMMGRRRRSKDIG----KYRRYACR
 Na-NLP-56   (75) -----YDRRYGSKVDKVPLSTEQIDSTDGVIARMRRDGHML------KRYTCR
 Bx-NLP-56   (65) RPLASYQWLHKAEADNELAELIDSIDGQAIGDRERRSTGP-----VYKRYACR
 Gp-NLP-56   (58) ---NGREMTEWVKVFNARAQKGRDRRSEGEAIKREFSISFKNFKIRNLGYTCR
                  160      171
Ce-NLP-56A  (106) ARIRLPLQIL--
Ce-NLP-56B  (106) FKFCRIYDA---
Ce-NLP-56C  (106) ARFVVCQPVLVI
 As-NLP-56  (154) FKFCRIFD----
 Na-NLP-56  (117) FKFCRIFDE---
 Bx-NLP-56  (113) FKFCRIFDA---
 Gp-NLP-56  (108) FKFCRIFSPTDE
```

NLP-57

```
                 1                                               50
Ce-NLP-57    (1) -------------------MRFILGLLIAIVAFVASSPIHGIWNNLPAPP
As-NLP-57    (1) ---------------------FLEGKRNSQAVLINEMTERSSTSKSELFQ
Bm-NLP-57    (1) -----------------MKQIFSFALISLLLNLLHASPIASIWGNLPDSA
Na-NLP-57    (1) MMNIQVWTFQRRIREAVGKVLFLRDIGSPRGYRTNGKERQDGNDMVLKEF
Bx-NLP-57    (1) ---------------MSRSLALLFALFMAAVIMVHCAPIDS----LPEME
                 51                                             100
Ce-NLP-57   (32) QK----RVYGFYNYLPKEEDDRDKRNTILLLTPNEDYVE-----------
As-NLP-57   (30) DK--K-SIYGLYRHLPGYYDDDTVEKKSAFFGIPLNEME-----------
Bm-NLP-57   (34) AQSMKRSIYGLYHHLPGYAYNEQPEPEIYVLPPAYRSWTN----------
Na-NLP-57   (51) EATIQKKKYGYYVRLPWKKDSADLPDNKAMSFRRLQTIVAKLQKNPELMQ
Bx-NLP-57   (32) KK----SIYGLYQYIPKASGVKS-KKFSSDPGPMIGQLPDGSYVIFGY--
                 101
Ce-NLP-57   (67) ----------
As-NLP-57   (66) ----------
Bm-NLP-57   (74) ----------
Na-NLP-57  (101) QYHNTFMNQL
Bx-NLP-57   (75) ----------
```

NLP-58

```
                           1                                               50
Ce-NLP-58    (1) -------------------------------------MISKCSVMGLLLL
Ts-NLP-58    (1) ----------------------------MLVLLGVVMVGFSELQVAPLLE
Tm-NLP-58    (1) ----------------------------------FLQQSLSERDNDVAYL
Na-NLP-58    (1) MVERRARVGLGENCHKELLEYHTKQPFKQAFLSSGIHAAFLADLKNLKKK
Hc-NLP-58    (1) ------------------------------------------------IV
                 51                                             100
Ce-NLP-58   (14) LFVHITTAQFETHFDDAQVYWPSQYKRSGPSSAS----EGEAYAFPGLRG
Ts-NLP-58   (23) NEYDLSSSSSSSSWPGLAKSFKILYSNDNSNMVPNQLWTIVDRRSVPFFG
Tm-NLP-58   (17) DQTYPSWNFGFPRWFDGETYQPVQLSTINR-------------RSVGFFG
Na-NLP-58   (51) VAREQTAFCGRSQQYPYLDLVPEYVKRSDF--------VDAGPTFRGLRG
Hc-NLP-58    (3) IAVEIETTCIFFQQYPYLDLVPEVYKRSLYG-------VDDGFTFKGFRG
                 101                                          148
Ce-NLP-58   (60) LRGKRDPTYHKRVPMMSLKGLRGKRARIFDGQEEQ-------------
Ts-NLP-58   (73) VRGK-------KVPFFGARGKRLRPVVIGAGEDERYKKLSMDFFGPRG
Tm-NLP-58   (54) SRGK-------KVPFFGSRGKRLQMDNVHESYQGDGKNKKLDFFGVRG
Na-NLP-58   (93) LRGK-------RMPYMNLKGLRGKRML---------------------
Hc-NLP-58   (46) LRGK-------RMPYMNLKGLRGKRTF---------------------
```

NLP-59

```
                 1                                               50
Ce-NLP-59    (1) --------------MSALHIARLFLLIAIMSTVVSTYAVLPS-------G
As-NLP-59    (1) MVIAQILFALAISSPVMCFLEPLPSLIDTSEGNAATSTTLQQPKFFSIAP
Na-NLP-59    (1) -----MAYCSTPGTMFVQLAVIWCVLIAQAFSQPSSSDVLLQRLHYINTP
Hc-NLP-59    (1) --------------MIVHVASVWCALVAQALSLTIPSDELEKRLQYMEVP
                 51                                             100
Ce-NLP-59   (30) EILFRRPAS--------------------------LFDRAADYSDDVAKF
As-NLP-59   (51) YLRYVNAFRENQLEHLNDNLGNMNARVIYSIPKHFDDDPTWKMMDIDQQQ
Na-NLP-59   (46) VVRYIAEN--------------------------EGLRKSLRGLNDAMRQ
Hc-NLP-59   (37) IVQFVEGREALKENFRRMHPEYMEVPIVQFVEGREALKENFRRMHPELRQ
                 101                              136
Ce-NLP-59   (54) DESRDKRELDGIDTQLSKRTNLKRLVILSARGFGKK
As-NLP-59  (101) MNINRRSLSRFIGASNEKRNALQRLAMNGARGFGRK
Na-NLP-59   (70) KRFEGVQDYTSLEDPYMKRTSLKRLAILSARGFGRK
Hc-NLP-59   (87) KRFEGLADYVALEDPNAKRSSLKRLAILSARGFGRK
```

NLP-60

```
                 1                                               50
Ce-NLP-60    (1) --------------------------------------------------
Ts-NLP-60    (1) MLAKYVQVILISRAGTALASLRQRASITQPQWTTAIASIIRIKSRNVGGH
Hc-NLP-60    (1) --------------------------------------------------
                 51                                             100
Ce-NLP-60    (1) -MSSSSSTFISIFIITMVLVAFCSANLATGRASLRPSAGKRAAVVGRLPA
Ts-NLP-60   (51) SRMSVMFILLSRFDRFSSLLGPANLNLATPIAGCRPSTGRREQLLKTPEQ
Hc-NLP-60    (1) ------MYMHSQCIILCVLVACGSSNLFVGRMSTRPSAGKRSTGG-----
                 101                                            150
Ce-NLP-60   (50) HYYQLASAVRSPKQYENEFTCNENTMSVINARYQKLQEELNGLVELMEAC
Ts-NLP-60  (101) AG-KQADGLKSLLECKGGRQISALKHLSMLAGYERRSKTR----------
Hc-NLP-60   (40) -----LEPFRRIFQNSGQFCISGIVLNYALTRF-----------------
                 151
Ce-NLP-60  (100) QTLQSTISL
Ts-NLP-60  (140) ---------
Hc-NLP-60   (68) ---------
```

NLP-61

```
                  1                                               50
 Ce-NLP-61    (1) --------------------------------------------MSHWIA
 As-NLP-61    (1) ------------------------------MGFAVQKTFGTSFIPSRKLL
Bm-NLP-61.1   (1) MEQTATVCILKSSNVGTYVSIVQVKRNFFSLLFEEITLSKFCMLLAMKVA
Bm-NLP-61.2   (1) ------------------------------------------MLLAMKVA
 Di-NLP-61    (1) ----------------------------------------------MKMT
 Na-NLP-61    (1) ----------------------------------MRNVIQMKSLMGVLFV
 Hc-NLP-61    (1) -----------------------------------------------VIS
 Bx-NLP-61    (1) --------------------------------------MFTVTAFRTMFL
Gp-NLP-61.1   (1) ----------------------------MLSIFREKNNGGSSHLSTAFIG
Gp-NLP-61.2   (1) -----------------------------MATSKNDSSRQITPLAAFAFL
                  51                                             100
 Ce-NLP-61    (7) TLLAFILIANAFAALSTGADVGASDDVRSYVMPFYG--------------
 As-NLP-61   (21) QFSTITIMLSNDTHVKFLFAGSPSRAVPVAAFERYPLLTIEDIGFHP---
Bm-NLP-61.1  (51) SSLFLIFVFLWTSVVSKSIG---QIEAPLMEPMR----------------
Bm-NLP-61.2   (9) YSLFLTFVILWTSVVPKSIG---EIEAPLMEPMR----------------
 Di-NLP-61    (5) YTLVFVFVFLWASVIPQSMEKSNQIEIPIMELNR----------------
 Na-NLP-61   (17) TALLVLFGLATPKPIDINEECGPFVGSRKIKFTKS---------------
 Hc-NLP-61    (4) SSSMEVFASTFNHIYNLDGS----HISRILNFS-----------------
 Bx-NLP-61   (13) VVFVVFVCISFSSALPAPAGYGYDLAPSAFEFPQLN--------------
Gp-NLP-61.1  (23) LFLAALFVARTAGQSDSSTAIEYEIVLPVSAADMSIGPRAWQGEWVR---
Gp-NLP-61.2  (22) AIMAALMSSAETAAMPQSHGVSRMVLVGEDQQPSLFLLLGPHGWVQQQPM
                  101                                            150
 Ce-NLP-61   (43) -------------------------------------MEKKWSRREPSIR
 As-NLP-61   (68) ----------LELAR-----------------------SKKWTRLEPSIR
Bm-NLP-61.1  (82) --------------------------------------YKKWTKLEPSVR
Bm-NLP-61.2  (40) --------------------------------------YKKWTRLEPSVR
 Di-NLP-61   (39) --------------------------------------YKRWTRLEPSVR
 Na-NLP-61   (52) --------------------------------------QKKWSKLEPSIS
 Hc-NLP-61   (33) ---------------------------------------KKWSSLEPSIR
 Bx-NLP-61   (49) -------------------------------------MAKKWSRLEPSIR
Gp-NLP-61.1  (70) ----------RFSPRVVQNGWKYRTAPRNDFGAHRIVMEKKWSPLEPSIR
Gp-NLP-61.2  (72) VFGGSAAPRNRFLPPLSSGTSYRGNKVGEDSGVAEGKVEKKWSRFEPSIR
                  151                                            200
 Ce-NLP-61   (56) FFKRNGGGQDLPPSRFLWDY------------------------------
 As-NLP-61   (85) FIGGSNPYFI----------------------------------------
Bm-NLP-61.1  (94) FLTNDEANERKPSRKAKHIDTNTLYNRTYISSIYACQPALLNKVLFNNLT
Bm-NLP-61.2  (52) FFDKRWTRLEPSVRFFDK--------------------------------
 Di-NLP-61   (51) FFDKRWTRLEPSVRFFDKR-------------------------------
 Na-NLP-61   (64) REGVQRASHISRRCCPV---------------------------------
 Hc-NLP-61   (44) --------------------------------------------------
 Bx-NLP-61   (62) FFKRSQFDSDFDN-------------------------------------
Gp-NLP-61.1 (110) FL------------------------------------------------
Gp-NLP-61.2 (122) FF------------------------------------------------
```

NLP-62

```
                1                                               50
Ce-NLP-62   (1) MVKFFVFLLFFAFLCSFTSAIPLRSLFLRSYDDINQESIARGYFAPQPVD
As-NLP-62   (1) -------MKSNTTVCSKKYWLHQEAIFTPVERANEGKRAES--DRQMRLY
Na-NLP-62   (1) -------MWPLRVLCLCLLAIGSIHGFELFQPYWGYNPRTLIRGSDPDFN
Hc-NLP-62   (1) -------MWPLRVLCFCVLLLVSAHAFEGYKPHLAYYRQ-----SEPEVY
                51                               85
Ce-NLP-62  (51) DDNADNRPKRGIDLLKRRVEIIERNRCFFNPITCY
As-NLP-62  (42) LHRASRLPPRIRKHVT--RSQVAKMRCFFNPISC-
Na-NLP-62  (44) LENVRPLWKRQSAEQSKRREPLSRSRCFFNPITC-
Hc-NLP-62  (39) LEENHPLLKRHVIEVK-RKEALARARCFFNPITC-
```

NLP-63

```
               1                                               50
Ce-NLP-63   (1) ------------MLVSLLICLALVAVASAMPFYASSNDGDVVRQFPQQGV
Na-NLP-63   (1) -----------------------YYSQNFPCFQMVDSNDSAGLRVPLQDD
Hc-NLP-63   (1) MPIRRLPLLVTVILLSLCTSAAAQHRLFYMLGNAVDGTSAAGLHVPRQDD
                51                              84
Ce-NLP-63  (39) LENMVNEIMRVQSVEQMNNHHVRRQIFSRKFW--
Na-NLP-63  (28) LQSIVSQLLTRVNDNNSEARHVRKQIFSRKFWKR
Hc-NLP-63  (51) LQSMVSQLLTRVNENNSDALHVRKQMFSRKFWKR
```

NLP-64

```
                 1                                               50
Ce-NLP-64    (1) -----------------------------MMAQKTLIIAVMLVCSILQPM
As-NLP-64    (1) --------------------------------------------------
Bm-NLP-64    (1) --------------------------------------------------
Di-NLP-64    (1) --------------------------------------------------
Na-NLP-64    (1) -------------------------MSRSLCLFLALVSLTHSFSLGNYYV
Hc-NLP-64    (1) -------------------------MSRTLCLVLAFIAVTNAFNLGNYYV
Bx-NLP-64    (1) MQPALVICLVASVVGTAHAFYALPPSFQRYMSDRSMLRPSAPLPQLVPFL
Gp-NLP-64    (1) ------MRSTNRTIPFTGVLLLLLVLFMAKFCHCDTNFRLSNFLPSYNRL
                 51                                             100
Ce-NLP-64   (22) LALGSLTPSAAFRANMQQRERSPNTLFYMDG-ASKQYGDEIKDPIYKRFK
As-NLP-64    (1) ---------MEHVKQYEEPLQKYNSKVVDDNAGILKRATYLRQSPLKRLR
Bm-NLP-64    (1) ------------FDNIKLNFRKKDETIYKDKYGYLSMKDFGEQRPLKRLR
Di-NLP-64    (1) --------------------XRNDPTIYKDDYGSLMLNDSG----MKRLR
Na-NLP-64   (26) GALPYRFMDSPYENRIAFGQWRGGRDFPVG--DSALQKDKKDTLPMKRFK
Hc-NLP-64   (26) GSMPYRYLDAQYDGRLGFGPLRGGREFGIDNPTPELQKDKKDAAPMKRFK
Bx-NLP-64   (51) PPVPIALEEEPIAAEPHPNNAKGYEEVQVPR--RSLRQFIGETHQMKRMR
Gp-NLP-64   (45) AAINPDDDQQLPVDEDEAVVANAGRLFADDDDEQPKQMHTMNQQVVLPER
                 101                                            150
Ce-NLP-64   (71) PCYYSPIQCLIKRK------------------------------------
As-NLP-64   (42) PCFYSPIQCLMKRANVHGISANLAP-------------------------
Bm-NLP-64   (39) PCFYSPIQCLMKKRTARIENFMDNYR------------------------
Di-NLP-64   (27) PCMYNPIQCIVKRLKLRRR-------------------------------
Na-NLP-64   (74) PCYYSPIQCLIKRR------------------------------------
Hc-NLP-64   (76) PCYYSPIQCLIKRR------------------------------------
Bx-NLP-64   (99) PCFYSPIQCLMKKRSQP---------------------------------
Gp-NLP-64   (95) PCYFSPIQCLLRTPGQHEVFENHPKFNVFRADLQRPTIDQSVIGASRQHQ
                 151                                         197
Ce-NLP-64   (85) -----------------------------------------------
As-NLP-64   (67) -----------------------------------------------
Bm-NLP-64   (65) -----------------------------------------------
Di-NLP-64   (46) -----------------------------------------------
Na-NLP-64   (88) -----------------------------------------------
Hc-NLP-64   (90) -----------------------------------------------
Bx-NLP-64  (116) -----------------------------------------------
Gp-NLP-64  (145) QVRRHLLGLMNSKSLVVTPSSSARHKLQASEIGRQRLFGGNPWKLGL
```

NLP-66

```
                1                                               50
Ce-NLP-66   (1) MSSGKLFQFFIVFLATLLLADAIPMVSSRDEDDQIIQKRLSNDALIRLLM
As-NLP-66   (1) ------MRPITSANFISNGRSPFAEMKFAEHDTRRLQRRSP--SIFPVLD
Na-NLP-66   (1) ----MSFLKYIVVLSLSSFLIIHALAMFNENEVPTIKKRLSNDALIRLIM
Hc-NLP-66   (1) -----TMIYFAPVYFLLLLFIPTDNVAMPEEDLLEITKRLSNDALIRLLI
                51                                            98
Ce-NLP-66  (51) RNRGTQTQLGLKRGLVKKAEVERRSIDEDFSNCFLSPVQCMLPSSRK-
As-NLP-66  (43) R-------R------------GFDTGDADFQNCFLSPVQCMLPRMRRR
Na-NLP-66  (47) RN---QHSMFTSKRDEKRADFDRRSVDDDFSNCFLSPVQCMLPRSRR-
Hc-NLP-66  (46) R-----FAMQRKR-HDKRAEVDRRSVDDDFSNCFLSPVQCMLPRMKR-
```

NLP-67

```
                 1                                               50
Ce-NLP-67    (1) ---------------MRVLTFLLVTLFALANVMQAQRYDRAIYEALLNDL
Na-NLP-67    (1) ---------------MKSVTLLFVLLLTVMHLSLSISHDRSIYEALLKNM
Hc-NLP-67    (1) ---------------MKSLALLFVALLALVQVSIAISYDRLVYEALLKNM
Bx-NLP-67    (1) MSLSSAMLTALRESRGVLMFSLLTVLLISTVTAKPLSQDRRSLQILLRAI
                 51                                             100
Ce-NLP-67   (36) EREFVERELAQHVLEK-RELLRQDRQELDRVRRASEKKSYPRNCYFSPIQ
Na-NLP-67   (36) EQELAERELLRIIS---RRERAIVAESEPRERRSSEKKSYPRNCYFSPIQ
Hc-NLP-67   (36) EQEMAERELVRELS---RRVRAVPVEVEQRERRSSEKKSYPRNCYFSPIQ
Bx-NLP-67   (51) DNELESPYSDDLMPSRSRRAVASELPQDKRELLAYDKKSYPRICYFSPIQ
                 101
Ce-NLP-67   (85) CLFTRN----
Na-NLP-67   (83) CLFTRS----
Hc-NLP-67   (83) CLFTRS----
Bx-NLP-67  (101) CLFTRMSLKK
```

NLP-68

```
                 1                                               50
Ce-NLP-68    (1) -MLLVLLFSLFSVGFGMRVMRRELLKDQPTVLLLGPLEVLTSSSSE-DDT
Na-NLP-68    (1) -MYLAVLLAFLSISCRASFVGHSPPNSPLTGRHALGESIEYNGSGESSKE
Hc-NLP-68    (1) MYSVTVLSALLCISSGATFYGNS-LNSDFSGDQALYYVNEYPTSEEPQKR
                 51                                             100
Ce-NLP-68   (49) PDFPSLRDKRGVDPMSIPRLIKEPPLKKRSGDFKRGDVVYPSAAKQTVPL
Na-NLP-68   (50) PVLESIVTKRGVDPLSIPQLIREPPLKRG---DKRDSYVYPSTSSKLSDR
Hc-NLP-68   (50) LVREPPLIKRGIDPFSIPTLIKEPPLKRG---DKRDSYVYPSTNSQLSDR
                 101                                            150
Ce-NLP-68   (99) VRVREPPLKR--G----------QMFLEELLQFNDDAHRQFMDPLRRIRY
Na-NLP-68   (97) VPLREPPLKR-------------QEYFELSNLFSPSEKIVEWQDTTDENS
Hc-NLP-68   (97) IPLREPPLKRSQSLDDKVFVLVPEERHEPNGLANVYRRMLLLDQSTNHAA
                 151                172
Ce-NLP-68  (137) G-----------PNRLIYTW--
Na-NLP-68  (134) G-----EGKSNGKHVLLRLDQ-
Hc-NLP-68  (147) DSEAVPHPKLRSQRFLSRIWWH
```

NLP-69

```
                  1                                               50
 Ce-NLP-69    (1) MHFFPILLLSILLILISTCSSTLVNNSPTAAFDTNAYSELNAKAKASMRR
 As-NLP-69    (1) --------------------------------------------------
 Na-NLP-69    (1) --------------------------------------------------
Hc-NLP-69.1   (1) ------------------------------------MCTVFARPSKNFEM
Hc-NLP-69.2   (1) -------------------------------------------MRQLEPL
 Bx-NLP-69    (1) --------------------------------------------------
 Gp-NLP-69    (1) --------------------------------------------------
                  51                                             100
 Ce-NLP-69   (51) LADILDFEMYQRRLSAAPDNSYYYQISPPHHQRISSLPLTIFPLPDKRLH
 As-NLP-69    (1) --------------------------------QQTLQPEDDR-PLVKRYN
 Na-NLP-69    (1) -----------TLRLNHEIEREIYRRLSGGAKPRLRRKINSMEIPIKRIT
Hc-NLP-69.1  (15) FLPADYRQLGNTLQRVDDTDQEMFQGYYDRPMARLQRKVNTFALPIKRIP
Hc-NLP-69.2   (8) GTLFQDAPQCVDAPAGDVHDQEVFQGYYDRPMARLQRKVNTFALPIKRIP
 Bx-NLP-69    (1) -----------------RGKQIHGTFEFSALSATVIQFQDDRDQRHKRYG
 Gp-NLP-69    (1) --------------------------------------AGRSPTVNKRFN
                  101      112
 Ce-NLP-69  (101) RIGGNIVMGK--
 As-NLP-69   (18) RIGGMIVMGRRK
 Na-NLP-69   (40) RIGGNIVMGRKK
Hc-NLP-69.1  (65) RIGGTIVMG---
Hc-NLP-69.2  (58) RIGGTIVMG---
 Bx-NLP-69   (34) RIGGTIVMGR--
 Gp-NLP-69   (13) RIGGNIMMGRK-
```

NLP-70

```
                 1                                               50
Ce-NLP-70    (1) --------------MSVSRPLNLAILSILVGLLYLSACTCAPSVSAHHLG
As-NLP-70    (1) MLDDELRFLPRRMRQKPFADSSSGAITKRTASRRSISISVYVTIIISDQP
                 51                                             100
Ce-NLP-70   (37) LRLKKWYEWNN-DMEITKKWYDWQNVPHALQQ-----------KRQPFET
As-NLP-70   (51) LRPKKWYDWNDDETRLHKRWYDWQALPLSDSIDLMKKNRLSHLERQLHPG
                 101
Ce-NLP-70   (75) SDYIIE
As-NLP-70  (101) NDWLLG
```

NLP-71

```
                  1                                                    55
 Ce-NLP-71    (1) -------------------------------------------------------
 As-NLP-71    (1) -------------------------------------------------------
 Bm-NLP-71    (1) -------------------------------------------------------
 Di-NLP-71    (1) -------------------------------------------------------
 Na-NLP-71    (1) MNTEEGSSTDEKEIVDVLTHHAGEHEWMQPGRTDVETGRASNVDDRKERGRVPLW
Hc-NLP-71.1   (1) -------------------------------------------------------
Hc-NLP-71.2   (1) -------------------------------------------------------
 Bx-NLP-71    (1) -------------------------------------------------------
                  56                                                  110
 Ce-NLP-71    (1) -------------------------------------------------------
 As-NLP-71    (1) -------------------------------------------------------
 Bm-NLP-71    (1) -------------------------------------------------------
 Di-NLP-71    (1) -------------------------------------------------------
 Na-NLP-71   (56) DGQLNATLRTKMNNLQTPECREQMNRGGGYITCRSSAKANAKDGKSGIMSDRNLT
Hc-NLP-71.1   (1) -------------------------------------------------------
Hc-NLP-71.2   (1) -------------------------------------------------------
 Bx-NLP-71    (1) -------------------------------------------------------
                  111                                                 165
 Ce-NLP-71    (1) ----------MKSSSVLSVALIVLVIVQLISASLASVPSSS---AVSDGQIDFDA
 As-NLP-71    (1) -------MTSCCRLALVFAVIVFHSQWFTNATSVGWDYSKE-----NS-LQRQGG
 Bm-NLP-71    (1) -----MAVFLGKIFEKSKIPFESVKLICSDVFCILKIFSRE----NLAMSQLFPW
 Di-NLP-71    (1) ----------MLKTSKTTGIISSVSRTNSAAISLWDEESRNNAWGNLATSQLFPW
 Na-NLP-71  (111) HNSEVLERLECEDVKRRESLYLFLMMSDVEDVTPHLHFRKI-----ASAKTEQGF
Hc-NLP-71.1   (1) ------------MQTAIAYLLAFILFVQCLGSSVGWAGERQ-----DDSEQVADP
Hc-NLP-71.2   (1) ------------MQPTVTYLLALILFIRCFGPSVGWEAER------KVDGQALDP
 Bx-NLP-71    (1) -------------------MQNGTFLANGSGPPLLYGMVAQ-----ESQDPLAYF
                  166                                              217
 Ce-NLP-71   (43) LAAKIEMLRPNRYWKR--AHNIDTRALNQFKNCYFSPIQCVLMERRRK----
 As-NLP-71   (43) ISRPIASWEAALLEQKRAVHNVDARSISQFKNCYFSPIQCVLVEKRK-----
 Bm-NLP-71   (47) HTISHEQTRHLEHQQQQPKHHIVLRSTNHFKNCYFSPIQCVLVEEQ------
 Di-NLP-71   (46) RTISHKQTRHYEQRK--PEHHIVLRSANHFKNCYFSPIQCVLVEEQ------
 Na-NLP-71  (161) GRNVLRQIGAYRLWKR--AHNIEPRALSQFKNCYFSPIQCVLMERRRR----
Hc-NLP-71.1  (39) RTAALRQMVPYRLWKR--AHIIEPRALNQFKNCYFSPIQCVLMERRRR----
Hc-NLP-71.2  (38) RTIALRELYPYRLWKR--SPYAGRRAAKRFKNCYFSPIQCVLMDRRRR----
 Bx-NLP-71   (32) VQNPPNEVFNAAAKRN--IHRVQARAVNTFKNCYFSPVQCVLLERRRRSVQV
```

NLP-72

```
                 1                                               50
Ce-NLP-72    (1) -------------------------------------MLTRVPVLILAVI
Ts-NLP-72    (1) ------------------------------------------MDLKIKIR
Tm-NLP-72    (1) MALYDRIKPGDVFRDYNGELSNRQSEAASLNQSTVALGKRLRRIMSLKVV
As-NLP-72    (1) --------------------------------MSISTQLFVSLCILTLMV
Na-NLP-72    (1) ---------------------------------MATSTMHSWLRFVLVCC
Hc-NLP-72    (1) --------------------------------------MQFWLRLTLVFC
Bx-NLP-72    (1) ------------------------MSLWAAKGSSRRAWTSWLTCFCLIIC
Gp-NLP-72    (1) --------------------------------------MFFALFLSTAVQ
                 51                                             100
Ce-NLP-72   (14) VMLALCQEPEKPEKRP----ALLSRYGRA-VLPRYGKRSGN--LMESSQ-
Ts-NLP-72    (9) IIFFLIGMALFSEAKP----ASLARYGRA-ALPRYGKRAELISGLDAMR-
Tm-NLP-72   (51) AFVILLVSVLAEAKGP----LAMARYGRA-ALPRYGKRMEISNGPEYFR-
As-NLP-72   (19) TAEFATNDILEPTKRA----ALLSRYGRA-VLSRYGKRSLPQSGYSSFG-
Na-NLP-72   (18) FAVLIIASDQVPEKRP----ALLSRYGRA-VLSRYGKRSPLPKVAAPENE
Hc-NLP-72   (13) FVVLLIADNQNLEKRP----AHLSRYGRA-LLSRYGKRSVISPNNGKYF-
Bx-NLP-72   (27) YLMVMVTAEFEKAEQPKRGNALLSRYGRA-VLSRYGKRSAPNFFESDVP-
Gp-NLP-72   (13) IRSHILPPQGANEAAG--GTILLSRYGRAAVLSRYGKRDGGAERREFNQS
                 101                                            150
Ce-NLP-72   (56) ------------NSLTEE-SSDVVCQLIDGKYICLPVDAVRFR---PFFL
Ts-NLP-72   (53) ------------DEVTEFKNSPCVYTGYEDLYRCSSMT------------
Tm-NLP-72   (95) ------------DDILEYRNNMCVYTGYEDLYRCSPVT------------
As-NLP-72   (63) ---EQTDVSPRIKTFALVLFEYCTERHIEGEYLRSCLNGVVFRRGEGVRK
Na-NLP-72   (63) TEIRQYRYRKENLSLYSSNHKTGEKKDFTELLLCRSVNGEFLCVPYTDEV
Hc-NLP-72   (57) ------------FLKVLSKRPSIKHRQTKSRSVVFHVHRIFFFNSSPVFY
Bx-NLP-72   (75) -------------FDGIAEEWLLCRRAPNDMLQCLPTPLYHY--------
Gp-NLP-72   (61) FGEGERAVHPPVAGPGPQCTHRLQGRARSAPTGCRAGPAVHPPVAGPGPQ
                 151                       179
Ce-NLP-72   (90) -----------------------------
Ts-NLP-72   (79) -----------------------------
Tm-NLP-72  (121) -----------------------------
As-NLP-72  (110) GEKGGMNDRIGEREIDIYYAVAKCLLYHI
Na-NLP-72  (113) RK---------------------------
Hc-NLP-72   (95) -----------------------------
Bx-NLP-72  (104) -----------------------------
Gp-NLP-72  (111) CTHRLQGRPPPPDPK--------------
```

NLP-73

```
                 1                                                    55
Ce-NLP-73    (1) MSCSSSSMLFLVLIATTVLIAESRVFYNRFDGGLSSDRFMEQKRDGAEASYDYDA
As-NLP-73    (1) -------MLTKYARVLFMLVVALKFTSAMITPSYTSIDYLNRRLQQANAYHEPII
Na-NLP-73    (1) --------MKFCLALLALTVIAATIEARLFGYEDVLVDPTQDRDSSSQNEWG--Y
Hc-NLP-73    (1) -------MNSTVSLCFVLVCFAAVIEARLFQLGDVLVDPSKDRDPTWIADW----
                 56                                           103
Ce-NLP-73   (56) NQVIRNTMKRNRQCLLNAGLSQGCDFSDLLHAQTQARKFMSFAGPGK-
As-NLP-73   (49) DDQSAPIMKRSRLCILNAGLSQGCDLSDMLFAKLQANKFSSFAGPGRR
Na-NLP-73   (46) KYSPARPSKRNRMCLINAGLSQGCDLSDILMAKQQASKFLSFAGPGRR
Hc-NLP-73   (45) EYALERPAKRNRMCLINAGLSQGCDLSDILMAKQQASKFMSFAGPGRR
```

NLP-74

```
                 1                                               50
Ce-NLP-74    (1) MIQTFFFLPNSNFSISFSLYFSQISVIFSPRVPGRFKSGRVAPSWCAGRR
Ts-NLP-74    (1) --------------------------------------------------
Tm-NLP-74    (1) --------------------------------------------------
As-NLP-74    (1) --------------------------------------------------
Bm-NLP-74    (1) --------------------------------------------------
Di-NLP-74    (1) --------------------------------------------------
Na-NLP-74    (1) --------------------------------------------------
Hc-NLP-74    (1) --------------------------------------------------
Bx-NLP-74    (1) --------------------------------------------------
Gp-NLP-74    (1) --------------------------------------------------
                 51                                             100
Ce-NLP-74   (51) EAESKTREICLSLGCAHSFYSTCASQPSEQAFPALPLPSSLNVNMNRFII
Ts-NLP-74    (1) -----------------------MRATCASVVTIRQHKREAYSIPSLRLL
Tm-NLP-74    (1) --------------------------------------------------
As-NLP-74    (1) --------------------------------------------------
Bm-NLP-74    (1) ------------------------------------------MVSNISKS
Di-NLP-74    (1) --------------------------------------------------
Na-NLP-74    (1) --------------------------------------------MRSFFS
Hc-NLP-74    (1) --------------------------------------------MRSFFA
Bx-NLP-74    (1) --------------------------------------------MKVSGL
Gp-NLP-74    (1) ------------------------------------------MSSGLSPA
                 101                                            150
Ce-NLP-74  (101) SMIALLAVFCAVSTASPLLYRAPQYQMY----------------------
Ts-NLP-74   (28) FSTIFVYLFLKAVEKYPIRLQIMNGKLIFGIIFLYATIFTYITAVPVVYE
Tm-NLP-74    (1) --------------------------------------------------
As-NLP-74    (1) -------MINAITIRESRLDNFG---------------------------
Bm-NLP-74    (9) TLRTFLLLILTVTIASTFEFGELMYGDN----------------------
Di-NLP-74    (1) --------------------------------------------------
Na-NLP-74    (7) VILALVALVFAVASANPLLYRVP--QMY----------------------
Hc-NLP-74    (7) IFLTLLVSVVVISTANPLLYRAP--QMY----------------------
Bx-NLP-74    (7) VQALFVLLVIALAQAQPLVYRPYGLHNVG------------------IPV
Gp-NLP-74    (9) QLTTFGEDNVNIGFSTANNSSANGTPTYTACTS------------FDEPH
                 151                                            200
Ce-NLP-74  (129) DDVQFVKRSNAELINGLIGMDLGKLSAVGKRS-------NAELINGLLSM
Ts-NLP-74   (78) ADLTAQEKEILQSLAEKIGRTYGSRFDSGRTIKR----ANAELVNGLLGM
Tm-NLP-74    (1) ----------LEALLERAGRRYLTRMASGNYIKR----ANAELVNGLLGM
As-NLP-74   (17) DGELLTKRSNAELINGLIGMDLNKLSAIGKRS-------NAELINGLLGM
Bm-NLP-74   (37) ELQGAWKRSNAELINGLIGMDLGKLAKAGKRSRTSSMITNDENINNDIRF
Di-NLP-74    (1) EMRGVWKRSNAELINGLIGMDLGKLTRAGKRS-VSPLITNDDNTNSN---
Na-NLP-74   (33) DDTQFVKRSNAELINGLIGMDLGKLSAVGKRS-------NAELINGLLGM
Hc-NLP-74   (33) DDVQFVKRSNAELINGLIGMDLGKLSAVGKRS-------NAELINGLLGM
Bx-NLP-74   (39) STYRHAKRSNAELINGLIGMDLNTLQAVGKRS-------NAELINGLIGM
Gp-NLP-74   (47) EFQAVEKRRNAEFINGLLTMERLNAIGKRSPP-----VPDVDDVQQLFWR
                 201        214
Ce-NLP-74  (172) NLNKLSGAGRR---
Ts-NLP-74  (124) RYGDLMRAG-----
Tm-NLP-74   (37) RYGDLMRAG-----
As-NLP-74   (60) NLNRLSSAGRR---
Bm-NLP-74   (87) KMNLKKLLSNQRHY
Di-NLP-74   (47) --------------
Na-NLP-74   (76) NLNKLSSAGRR---
Hc-NLP-74   (76) NLNKLSSAGRR---
Bx-NLP-74   (82) DLNRLSSIGRRR--
Gp-NLP-74   (92) PQHPFFWTE-----
```

NLP-75

```
                 1                                                         60
Ce-NLP-75    (1) ------------------------MGSSPILLVLAISIGLASACFLNSCPYRRYGRTIR-
As-NLP-75    (1) ------------------------------CAGRRIRTAILIDFFINQCPYRHYGRTQCA
Na-NLP-75    (1) ----------------------------QSLAFVIFSLSIVNGCFLNSCPYRRYGRTLR-
Hc-NLP-75    (1) SVKMDYPLRAAGGSFRRGRDELFNIMNWLCFAFLIVSTIIIDGCFLNSCPYRRYGRTIR-
Bx-NLP-75    (1) ----------------------MGDRTRTLALIICLLTMHCDGCFLNSCPYRRIGRSNQP
                 61                                                       120
Ce-NLP-75   (36) -----------------CSSCGIENEGVCISEGRCCTNEECFMSTECSYSAVCPELFCKI
As-NLP-75   (31) ------------------------------------------------------------
Na-NLP-75   (32) -----------------CALCAPSREGICHGDGQCCTHTHCFPSKDCGTRDSCPENFCRF
Hc-NLP-75   (60) -----------------CESCGEFMNFHR-------------------------------
Bx-NLP-75   (39) ANAVIPSGPSVDITVETCGICGPKNNGICVNERVCCTVEECKEDTGCQNVHACPLPLCII
                 121                             155
Ce-NLP-75   (79) GHHPGYCMKKGYCCTQGGCQTSAMC----------
As-NLP-75   (31) -----------------------------------
Na-NLP-75   (75) DSHSGVCITKLLCCTSSHCIRSMQCA---------
Hc-NLP-75   (72) -----------------------------------
Bx-NLP-75   (99) DNGPGFCATNTLCCADKFCQRNLQCMVVASKQLNL
```

NLP-76

```
            1                                               50
 Ce-NLP-76    (1) --------------------------------------------------
 As-NLP-76    (1) -----------------------------------------------MRY
Bm-NLP-76.1   (1) --------------------------------------------------
Bm-NLP-76.2   (1) --------------------------------------------------
Di-NLP-76.1   (1) --------------------------------------------------
Di-NLP-76.2   (1) --------------------------------------------------
Na-NLP-76.1   (1) --------------------------------------------------
Na-NLP-76.2   (1) MDIRGGDCFNSQGKAVEKRLEGGLAITVRLILFEAPVKTCIENLAWVGYF
 Hc-NLP-76    (1) --------------------------------------------------
 Bx-NLP-76    (1) --------------------------------------------------
 Gp-NLP-76    (1) -------------------------------------------------F
                  51                                             100
 Ce-NLP-76    (1) --MSRKFLLVFIAICVLSQNAMALRGALFRSGRSLIHR-------QSQNE
 As-NLP-76    (4) NRSMNAALLCSIVLLITLPTADALRGALFRSGRSMQRISRQSTDAKFPLD
Bm-NLP-76.1   (1) ----MKFSLSFILFFLMITTSMTLRGALFRSRKAALES-------IIRAN
Bm-NLP-76.2   (1) -MSATYGITCVFVGLALFTATNALRGALFRTGKSFWSSG------KYDPY
Di-NLP-76.1   (1) ------------VFFLLITTSMALRGALFRSRRASLDS-------IIRAH
Di-NLP-76.2   (1) -------YSGVFICLALFTATNALRGALFRTGKRLWSSD------EYDPY
Na-NLP-76.1   (1) ----MRAFIYVLAVLATTSSVFGIRGALYRTGRSATPAYG---KQIPMIK
Na-NLP-76.2  (51) QSFFKAHTIEALYMVLLTTSVLSLRGALYRSGRSFRFRQPTVVEHNALVE
 Hc-NLP-76    (1) ---MLRSLLFLVVLTAIPQTAHGVRGALFRTGRADLVPF------FDNVN
 Bx-NLP-76    (1) -----MFPKSFLVILLMPLMVECIRGALFRSGRSPGLQKTRGSQAIKMLY
 Gp-NLP-76    (2) KQKDVLLLLFVLLCCSAPPSAMAIRGALFRSGRSMAIPLWR-PPAKNTGA
                  101                                   141
 Ce-NLP-76   (42) EAVLNQPHLLPAEKRSPIVELYPVVDSGNVEPEAFPAYFRF
 As-NLP-76   (54) SSAFEFQHTMPFYPFGYRKRFDVDQELHDFRNE--------
Bm-NLP-76.1  (40) TGRQDFLRFGRTNEKYNDRHLIASALLTPVDLK--------
Bm-NLP-76.2  (44) TNLFELGKSLKAIDLMDNGNRNVISTNRDAKKTIYLQ----
Di-NLP-76.1  (32) ASRVRYLHKFKNLSLFE------------------------
Di-NLP-76.2  (38) MSLFKPSRSVEMVSLI-------------------------
Na-NLP-76.1  (44) NPQVEMPYGSQPVMVG-------------------------
Na-NLP-76.2 (101) TKRNVVVAYPPNSMESLDNQEANLRVFPNFLL---------
 Hc-NLP-76   (42) FISIK------------------------------------
 Bx-NLP-76   (46) SNLFRPDNGPLLRPQISYNNVIDNEQIFSRLRTLLMPFNAE
 Gp-NLP-76   (51) ASGYSAGLASTSEKLANYARPSVIVDLRQVVQVLVADSKS-
```

NLP-77

```
                 1                                               50
Ce-NLP-77    (1) -----------------------------------------MCRLAVFAL
As-NLP-77    (1) ----------------------------------------------MKVL
Bm-NLP-77    (1) ------------------------------------------MKMMKFVL
Di-NLP-77    (1) --------------------------------------------------
Na-NLP-77    (1) ----------------------------------------------MLKL
Hc-NLP-77    (1) --------------------------------------------------
Bx-NLP-77    (1) ---------------MWNVPCGHDRRETVNPEAPLKPQATIMSSLSCVLL
Gp-NLP-77    (1) MSEGVRTLKFRPSRRKHNNFSSTFTLDHSIPDMTSSSALLLVASLAMTVL
                 51                                             100
Ce-NLP-77   (10) LAVAAVSVYGQPAGGQD---VPPFLRNATPAQLQSFQQLIQANGHLTETA
As-NLP-77    (5) IIFVAIVVIAFAQGPQG---PPPFLVGAPANVVAEFKQIITGAPDKTDAE
Bm-NLP-77    (9) LAFPVLLLVCPVFGEEQQFMIPPFLLGAPESAVREMKELFQKYANKPDSQ
Di-NLP-77    (1) ----------MILEQLE---VPPFLVGAPQSVIKQFYDLLKADETKTDAQ
Na-NLP-77    (5) VALVCLVAICFAQGPQG---PPPFLQSAPAAVQQDFDKLFVNAGSKTDAE
Hc-NLP-77    (1) ---------------------------------------MANAGSLTDAQ
Bx-NLP-77   (36) VALFGLLALSQAQNIP----PPPFLAGESPDVIQSFQRVLERGNDLTDAQ
Gp-NLP-77   (51) ALCRAQLSAPSSQGAGQ---APPFLAEAGADKINEFYQFVASLGHLKDAE
                 101                                            150
Ce-NLP-77   (57) LDGKVQAWVNQQGGKVAADWADFQKFIKGQQGQAEAAHQAAVSNFSPAAK
As-NLP-77   (52) IDRDIENWVSRQGPKIKTEFNKFKTQMQQGKARAEAAHRASIAKFSPAAK
Bm-NLP-77   (59) LEAAVEEWVNGKGGMIKEKYNQFKANMQLMHEKTALLRKAMAENLSPEAK
Di-NLP-77   (38) TEADVEAFINRLGGTYKTRFDQFKQEIKQGKAAYERLHQQAVAKFSKEAR
Na-NLP-77   (52) IDKMVQDWVGKQDASIKTAFDAFVKEVKAAQAQGEAAHQAAIAKFSAEAK
Hc-NLP-77   (12) IDQKVQDWVNQQDPAIKTAFDAFVQQVKQAQAQSEAAHQAAIANFSPAAK
Bx-NLP-77   (82) LDREVDAWVQTQSAAVKNKYAQFKSDLAKHQAQAEAAHKAAVSRFSSVAR
Gp-NLP-77   (98) IDAKVKEWVGKQSDDVQKKYAGFEQEREKAKAQGEAAHEEAIKHFSEGAK
                 151                                          198
Ce-NLP-77  (107) KADADLTAISNDSSLSVQAKGQKIQAYLNSLPANVKAELEKAQGQ---
As-NLP-77  (102) AADAQLTAIADNPNLKGREKQQKITSLLQSLPAAVQAELQKEMQG---
Bm-NLP-77  (109) KADVDLSSIGKDKTLSEQQKREKFKEYLRNLSPAVRNELQAVFYAKF-
Di-NLP-77   (88) EADAKMSAIADSPSLTTQQKTQQIQAIMD-------------------
Na-NLP-77  (102) AADAKLSAIANDRSKTNAQKGAEIDSVLKGLPPNVRTEIENAMKG---
Hc-NLP-77   (62) EADAKLSAIAADPNKTNQQKAQEIQATLASLSPEVRNEIETAMKGGK-
Bx-NLP-77  (132) EADAKLSTIANNPSLTSAQKNAQIEKFVSQLPPAVRKEIESAMQG---
Gp-NLP-77  (148) KADKELSEIANNVNLSGQDKAKQINDKVQSLPKDVRDEIEGNVGGGRN
```

NLP-79

```
                   1                                               50
  Ce-NLP-79    (1) --------------------MSTRWFVFVALMALVLLSAHTTEAM-----
  As-NLP-79    (1) MKRQVAEIARMDRTTYAKRALATSLFVMCALCWLANATPLIDGNLRKQST
  Na-NLP-79.1  (1) ---------MNLRSFVSFRMQSSTRWSFLAIISLLLVVQSFANQG-----
  Hc-NLP-79.1  (1) -----------MDSLQVMMHDNKRTLLLVIVGLLLLVEQSYASLR-----
  Na-NLP-79.2  (1) ------------------MRYSVCWVLLAAFTIMSMTSARIHER------
  Hc-NLP-79.2  (1) ------------------MRYSICWVVLATACSLIFLRITQAELS-----
                   51                                             100
  Ce-NLP-79   (26) ------PRPGQMSLSNTRNCFFSPMGCVFVPKMSRIRKLSADFRNEIPPP
  As-NLP-79   (51) LSFVGNGMQQYENSRYDRNCFFSPMNCVFYEHPMDAPVRRFTFGEEALRR
  Na-NLP-79.1 (37) ------EIDRLLGARVERNCFFSPMGCMFLPGKKLSRVRRLPELADQDPQ
  Hc-NLP-79.1 (35) ------ELDRVLGARIERNCFFSPMGCMFLPNKKFARVRRLQESEDES--
  Na-NLP-79.2 (27) ------YADYAITLRSSRNCFFSPIGCVFFSSGVEFPRARHKKRHVNRRH
  Hc-NLP-79.2 (28) ------ALDYRIAARADRNCFFSPIGCVFLTGNKQIRARRHKNVDYPWSN
                   101
  Ce-NLP-79   (70) DYI------
  As-NLP-79  (101) ---------
  Na-NLP-79.1 (81) SERSLLWFI
  Hc-NLP-79.1 (77) ---SPLQFL
  Na-NLP-79.2 (71) ---------
  Hc-NLP-79.2 (72) ---------
```

NLP-81

```
                 1                                               50
Ce-NLP-81    (1) -------------------------MLKSSIFSLLIVLLMVCS-------
As-NLP-81    (1) ----------------------MCRPAQTAACALLLFLSLFVS-------
Bm-NLP-81    (1) ------------------MIYGTFLLLLHLPLLFAIAILNLHC-------
Di-NLP-81    (1) -------------------------XXXXXXWIFFLILLFFFL-------
Na-NLP-81    (1) -----------------------------MHAILGLLLAMVAA-------
Hc-NLP-81    (1) XDETHECLGILFVAIQLWSISAHRSSTWKMHVILRLLLAMVAA-------
Bx-NLP-81    (1) -----------------------MSPLSTVILALFMLLVVQSG-------
Gp-NLP-81    (1) -------MNISKSQNLSSSSMNYVPMFLRFVLLIACFLLLMSGPFAMAND
                 51                                             100
Ce-NLP-81   (19) -----FGNVSANDFFLRSAKWVSKMNPSGGALVSGRGGFRPGFVS--RDW
As-NLP-81   (22) -----STQTKPSDMFIRSPKWARLA-PSGGSLVSGRGNFRPGFVSSAADT
Bm-NLP-81   (26) -----QAVDATNDYFFRSPKWVNLG-PSGGSLVSGRGNFRPGFSSSVSDS
Di-NLP-81   (19) -----QIVDAENEYFAQPPKTVYLR-PSGGLLVSGRGNFRPGFLPSNIDN
Na-NLP-81   (15) -----VAAFSNSDFFIRSAKWIPSG---GAGLVSGRGNFRPGFTG--RDW
Hc-NLP-81   (44) -----AAAFTNGDFFVRSAKWIPSG---GAGLVSGRGNFRPGFTG--RDW
Bx-NLP-81   (21) -----KADDMDSYYFRRNAKWISLAP-SGGSLVSGRGNFRPGFYGR-GGS
Gp-NLP-81   (44) DVANELAMRGTQQFFRRNAKWTNLP-SGGGSLVSGRGNFRPGFHSS-LRY
                 101                                     143
Ce-NLP-81   (62) RHAMAEPNFVKRSYNDY--------------------------
As-NLP-81   (66) RMLLSEPLLIFKKQVLFQAKHISALSTVYQLVSSISADNRYVK
Bm-NLP-81   (70) RFYLSEPLFAFKRT-----------------------------
Di-NLP-81   (63) PFYFFEPLFALRV------------------------------
Na-NLP-81   (55) RSAMAEPNFNKRSAQPWFSEDY---------------------
Hc-NLP-81   (84) RSAIAEPNFRKRSVESWYSDEY---------------------
Bx-NLP-81   (64) AAYMGDPVLFKRSQPADLE------------------------
Gp-NLP-81   (92) LAQLSEPSFAFKRSPQHFPMTMDELELADNFLGTQ--------
```

NLP-82

```
                 1                                                         60
Ce-NLP-82    (1) -------------------------MPSYHTVIIILLISIISTTSFILQDLPLERFERSG
Na-NLP-82    (1) ------------------------FQMGFVRSVLVWCAIVCLTLSFVLQDLPLERYERS-
Hc-NLP-82    (1) MGGSFGPIIPLTTSYVRSLPYEVLSGMSFVRSLLLWCAVACLTLGFVLQDLPLERFERG-
                 61                                                       120
Ce-NLP-82   (36) HGSDELLGGGSQFDRHIRSGLSYRPMPNLMARYRMMSG----------------------
Na-NLP-82   (36) -------QREAYFERPYRALPGPAFIYNRFM-----------------------------
Hc-NLP-82   (60) -------YPEISFDRASQAECVDSELVSIVIIIVVIVMFITGLQAVVRIGNSLHSITVVL
                 121
Ce-NLP-82   (74) ---
Na-NLP-82   (60) ---
Hc-NLP-82  (113) FFE
```

NLP-83

```
                 1                                                         60
Ce-NLP-83    (1) ----------------MARFTPLLMILLALVPLYYSLDCRKFSFAPACRGIMLKRSGGHP
Ts-NLP-83    (1) --------------MCTSVLPFITIIVVSIQLSSCNINCRNYPFAPKCRGIILKKSVLS-
Tm-NLP-83    (1) MQFSVVPQTTTAYLTAHRSIVVIFCLSSSLTAYYAFEFFRKYPYAPPCRGIVLKRSPIYE
As-NLP-83    (1) -------------------MRIVIFALFALVPLVSALDCRKFSFAPACRGIMLKRSEGS-
Na-NLP-83    (1) -------------------MRIILFVFLSLIAIALSLDCRKFSFAPACRGIMLKRSETTE
Hc-NLP-83    (1) -------------------MRSLLFLIVAIIPLAIALDCRKFSFAPACRGIMLKRSEVAE
                 61                                                       120
Ce-NLP-83   (45) MIAEQQPMIDNAKAREVQMMELLIRNIEDEALIANSDCVSMSWLRDRLVNAKNEMPQ---
Ts-NLP-83   (46) -------------------LKANREIVEEDENPIIK------------------------
Tm-NLP-83   (61) P--------------LP--KTHMTHIATLK------------------------------
As-NLP-83   (41) ---------AVAFDTDMLPLIAVCWHFQELKLQKVRSRTLLFGK----------------
Na-NLP-83   (42) RQQ------SLELYQLAQNLEMLSLLIREAEAEGIKECIPISWIRNKLATKASDAPPPPL
Hc-NLP-83   (42) VES------PLDLDQLAQNLETLSWILRLAEADGIKDCVPLSWIKHKLTSSSWLTSRS--
                 121                142
Ce-NLP-83  (102) ----------------------
Ts-NLP-83   (63) ----------------------
Tm-NLP-83   (75) ----------------------
As-NLP-83   (76) ----------------------
Na-NLP-83   (96) FTVSRDSRDRNEAHAQNIRTHK
Hc-NLP-83   (94) ----------------------
```

NLP-85

```
                 1                                               50
Bm-NLP-85    (1) MSSFRLSTIYFCRTLLGTLICCYLLTDHVTAITDDSNIAKIPAYRTFQET
Di-NLP-85    (1) MPSFRPWINHFHKTLLSTLICYYLLADHATGINDGSNFMTYPAYR--QD-
                 51                                             100
Bm-NLP-85   (51) GYGLNEASHLRSLRSALGMDNIG---QITYPLTSRSAAMISGRGFRPGKR
Di-NLP-85   (48) LYGMNDASHLRSLRSDLGMNNIGHVEDFNYPLASRSAAMISGRGFRPGKR
                 101
Bm-NLP-85   (98) YDAEVISFM
Di-NLP-85   (98) YDAGIISFI
```

NLP-85 Pan-phylum

Note: additional species *Acanthocheilonema viteae, Brugia pahangi, Brugia timori, Ditylenchus destructor, Elaeophora elaphi, Enterobius vermicularis, Gongylonema pulchrum, Loa loa, Onchocerca flexuosa, Onchocerca volvulus, Onchocerca ochengi, Syphacia muris, Thelazia callipaeda, Wuchereria bancrofti.*

```
                            1                                               50
    A. viteae NLP-85    (1) --------------------------------------------------
    B. malayi NLP-85    (1) --------------------------------------------------
   B. pahangi NLP-85    (1) --------------------------------------------------
   B.  timori NLP-85    (1) --------------------------------------------------
D. destructor NLP-85    (1) MSGLHLPLLNSKPKMTSKTILYLMFGVFGMCLCQEGSAFEPTRTLSIATG
   D. immitis NLP-85    (1) --------------------------------------------------
    E. elaphi NLP-85    (1) --------------------------------------------------
     E. vermi NLP-85    (1) --------------------------------------------------
  G. pulchrum NLP-85    (1) --------------------------------------------------
       L. loa NLP-85    (1) --------------------------------------------------
  O. flexuosa NLP-85    (1) --------------------------------------------------
  O. volvulus NLP-85    (1) --------------------------------------------------
   O. ochengi NLP-85    (1) --------------------------------------------------
     S. muris NLP-85    (1) --------------------------------------------------
T. callipaeda NLP-85    (1) --------------------------------------------------
 W. bancrofti NLP-85    (1) --------------------------------------------------
                            51                                             100
    A. viteae NLP-85    (1) --------------------------------------------------
    B. malayi NLP-85    (1) ----------------------------------------------MSSF
   B. pahangi NLP-85    (1) ----------------------------------------------MSSF
   B.  timori NLP-85    (1) --------------------------------------------------
D. destructor NLP-85   (51) RGNIFRPSYAIGIPRFFYNGFQPYQFMNKRSMSSAVPAKFMSLAGEDSEL
   D. immitis NLP-85    (1) ----------------------------------------------MPSF
    E. elaphi NLP-85    (1) ----------------------------MNVQFAKKKFSATAKFQVMPSL
     E. vermi NLP-85    (1) --------------------------------------------------
  G. pulchrum NLP-85    (1) --------------------------------------------------
       L. loa NLP-85    (1) ----------------------------------------------MSPS
  O. flexuosa NLP-85    (1) ----------------------------------------------MSPF
  O. volvulus NLP-85    (1) ----------------------------------------------MSSF
   O. ochengi NLP-85    (1) ----------------------------------------------MSSF
     S. muris NLP-85    (1) --------------------------------------------------
T. callipaeda NLP-85    (1) ------------------------------------------------MV
 W. bancrofti NLP-85    (1) ----------------------------------------------MSSF
                            101                                            150
    A. viteae NLP-85    (1) --------------------------------------------------
    B. malayi NLP-85    (5) RLSTIYFCRTLLGTLICCYLLTDHVTAITDDSNIAKIPAYRTFQETGYGL
   B. pahangi NLP-85    (5) RLSTIYFCRTLLGTLICCYMLTDHVTAITDDSNFAKIPAYRTFQETGYGL
   B.  timori NLP-85    (1) --------------------------------------------------
D. destructor NLP-85  (101) DNDLQFMKRSAALGRTRFRPGKRSLVTMEELEDMGMLPGFSLLTEAQMGK
   D. immitis NLP-85    (5) RPWINHFHKTLLSTLICYYLLADHATGINDGSNFMTYP-AY---RQDLYG
    E. elaphi NLP-85   (23) RPSIIHFHKTLLMLLSCS-LLADHATGILDNSNYGRLPSSF---RLGRFG
     E. vermi NLP-85    (1) --------------------------------------------------
  G. pulchrum NLP-85    (1) --------------------------------------------------
       L. loa NLP-85    (5) RSSSIHFNRTLLGTLICCYLSADHVTGIIDDSNTAKFPVYR---QTGCGS
  O. flexuosa NLP-85    (5) RPWIIHFHRNLLATLICCYLLTSHVTGVFDDSSFK------------A--
  O. volvulus NLP-85    (5) RLWIIHYHRNLLGTLIYCYLLASSVTGIIDDSNFIRYP-AY---KKTAHD
   O. ochengi NLP-85    (5) RPWIIHFHRNLLGTLIYCYLLASSVTGIIDDSNFIRYP-AY---KKTAHD
     S. muris NLP-85    (1) --------------------------------------------------
T. callipaeda NLP-85    (3) RSQLLHWQ---IIYLICSCFWTDRVIANLYYPNFSRLSENKVDNERTPCG
 W. bancrofti NLP-85    (5) RSSTIYFCKTLLGTLICCYFLTDHVTAITDDSNFAKFPAYRTFQETGYGL
                            151                                            200
    A. viteae NLP-85    (1) --------------------------YPLASRSAVMISGRGFRPGKRYDE
    B. malayi NLP-85   (55) NEAS-HLRSLRSALGMDNIG---QITYPLTSRSAAMISGRGFRPGKRYDA
   B. pahangi NLP-85   (55) NEAS-HLRSLRSALGMDNIG---QITYPLTSRSAAMISGRGFRPGKRYDA
   B.  timori NLP-85    (1) --------------------------YPLTSRSAAMISGRGFRP------
D. destructor NLP-85  (151) RSMAVGRVGFRPGKRSIATGRAGALFWNPNYAKAAVISQRGFRPGKRSLT
   D. immitis NLP-85   (51) MNDASHLRSLRSDLGMNNIGHVEDFNYPLASRSAAMISGRGFRPGKRYDA
    E. elaphi NLP-85   (69) LNEVAHLRSLRSALGMENIGHVESFTYPLASRSAAMISGRGFRPGKRYDT
     E. vermi NLP-85    (1) -----------------------SFQIPILSRSATMISGRGFRPGKRAQS
  G. pulchrum NLP-85    (1) -----------------------KFQYPLNSRSAAMISGRGFRPGKRSDL
       L. loa NLP-85   (52) NEGSSHLRSLRSVFGMDSVG---QVVYPLTSRSAAMISGRGFRPGKRYDT
  O. flexuosa NLP-85   (41) IHELEFLRSLRSALERKSNGPLKNFNYPIKSRSAAMISGRGFRPGKRYDA
  O. volvulus NLP-85   (51) LNKASFLRSLRSVLGIDGTGYIEGFTYPLTSRSAAMISGRGFRPGKRYDA
   O. ochengi NLP-85   (51) LNKASFLRSLRSVLGIDGTGYIEGFTYPLTSRSAAMISGRGFRPGKRYDA
     S. muris NLP-85    (1) ------------------KRSLEYQEIPILSRSATMISGRGFRPGKR---
T. callipaeda NLP-85   (50) TNEVPYFRSIRSPILINNEDY-DSFAYPLSSRSAAMISGRGFRPGKRFNQ
 W. bancrofti NLP-85   (55) NEAS-HLRSLRSALGMDNIG---QITYPLTSRSAAMISGRGFRPGKRYDA
                            201                                  240
    A. viteae NLP-85   (25) GFISFM----------------------------------
    B. malayi NLP-85  (101) EVISFM----------------------------------
   B. pahangi NLP-85  (101) EVISFM----------------------------------
   B.  timori NLP-85   (19) ----------------------------------------
D. destructor NLP-85  (201) LSENADSWEQMNMVERLTDN--------------------
   D. immitis NLP-85  (101) GIISFI----------------------------------
    E. elaphi NLP-85  (119) GIISFM----------------------------------
     E. vermi NLP-85   (28) TVSYPIVLYRINVSPLSFPFFSLFSSSTTFCIIDIQIYPV
  G. pulchrum NLP-85   (28) RNIAFM----------------------------------
       L. loa NLP-85   (99) EIISFM----------------------------------
  O. flexuosa NLP-85   (91) PILSFM----------------------------------
  O. volvulus NLP-85  (101) EIISFM----------------------------------
   O. ochengi NLP-85  (101) EIISFM----------------------------------
     S. muris NLP-85   (30) ----------------------------------------
T. callipaeda NLP-85   (99) NTAMM-----------------------------------
 W. bancrofti NLP-85  (101) EIIPFM----------------------------------
```

NLP-86

```
                 1                                               50
As-NLP-86    (1) MLDCVESAFVGGACCLYTNQRYSIPQQHSFSCIRCRDALVACGALRFDYR
Bx-NLP-86    (1) -----MRLLLALLFTLLAVGFADFLEPNELDQMEDIDVPRERRSSLASGR
Gp-NLP-86    (1) ---MTSLPLLTLLSVLLALQLIGQLFADKFEQVEDQDQLNQQQLTIGSNR
                 51                                             100
As-NLP-86   (51) GRCVSFSIRSLASGRWGLRPGKRS-QDVPIYTDELGLDGNSPLYDALRQS
Bx-NLP-86   (46) WGLRPGKRSSLASGRWGLRPGKRSIVDLVLEDDIEDESDKTRERRNLASG
Gp-NLP-86   (48) DIR------SLANGRWQLRPGKR--ASLIDFRPMDGLEERARRFRNLYAL
                 101                                      144
As-NLP-86  (100) HFVYVVRK------------------------------------
Bx-NLP-86   (96) RWGLRPGKRSLANGRWGLRPGKRSMGMMPTQARHNPIMLLIPQL
Gp-NLP-86   (90) LPPLNNWN------------------------------------
```

NLP-86 Pan-phylum

Note: additional species *Ascaris lumbricoides, Globodera rostochiensis, Meloidogyne arenaria, Meloidogyne incognita, Meloidogyne javanica, Panagrellus redivivus, Parascaris univalens, Syphacia muris, Toxocara canis*.

```
                               1                                               50
 A. lumbricoides NLP-86    (1) --------------------------------------------------
         A. suum NLP-86    (1) --------------------------------------------------
   B. xylophilus NLP-86    (1) --------------------------------------------------
      G. pallida NLP-86    (1) --------------------------------------------------
 G.rostochiensis NLP-86    (1) --------------------------------------------------
     M. arenaria NLP-86    (1) --------------------------------------------------
  M. floridensis NLP-86    (1) --------------------------------------------------
        M. hapla NLP-86    (1) --------------------------------------------------
    M. incognita NLP-86    (1) --------------------------------------------------
     M. javanica NLP-86    (1) --------------------------------------------------
    P. redivivus NLP-86    (1) MNKCNGASEPVGAHQDRFSSRRSKKPKAVYPQTVTSTKHFITLKNNGLHH
    P. univalens NLP-86    (1) --------------------------------------------------
        S. muris NLP-86    (1) --------------------------------------------------
        T. canis NLP-86    (1) --------------------------------------------------
                               51                                             100
 A. lumbricoides NLP-86    (1) --------------------------------------------------
         A. suum NLP-86    (1) -------------------------------------------MLDCVES
   B. xylophilus NLP-86    (1) --------------------------------------------------
      G. pallida NLP-86    (1) --------------------------------------------------
 G.rostochiensis NLP-86    (1) --------------------------------------------------
     M. arenaria NLP-86    (1) --------------------------------------------------
  M. floridensis NLP-86    (1) --------------------------------------------------
        M. hapla NLP-86    (1) --------------------------------------------------
    M. incognita NLP-86    (1) --------------------------------------------------
     M. javanica NLP-86    (1) --------------------------------------------------
    P. redivivus NLP-86   (51) PGGQLSLTTLLAKIENQRLGKSTSIASQNEVNKNPATSKNLTFEDSPTQL
    P. univalens NLP-86    (1) -------------------------------------LKVRLWEEHVASA
        S. muris NLP-86    (1) --------------------------------------------------
        T. canis NLP-86    (1) --------------------------------------------------
                               101                                            150
 A. lumbricoides NLP-86    (1) -------------------MRSLLAVLFVSIIVDVVYPSPLYVAGPSISK
         A. suum NLP-86    (8) AFVGGACCLYTNQRYSIPQQHSFSCIRCRDALVACGALRFDYRGRCVSFS
   B. xylophilus NLP-86    (1) ----MRLLLALLFTLLAVGFADFLEPNELDQMEDIDVPRERRSSLASGRW
      G. pallida NLP-86    (1) --MTSLPLLTLLSVLLALQLIGQLFADKFEQVEDQDQLNQQQLTIGSNRD
 G.rostochiensis NLP-86    (1) --MTSLPLLTLFSVLLALQLIGQSFADEFEQVEDQDLLNPQQLTIDSNRN
     M. arenaria NLP-86    (1) --------MRFSIALIVFAFILVIFN--CAESKDEGGNQEEIEQQYQNRR
  M. floridensis NLP-86    (1) --------MRFSIALTIFAFILVIFN--CAESKDEGGNQEEIEQQYQNRR
        M. hapla NLP-86    (1) --------MRFSIAFLVLTFILVISTMFVVESRDEGGNQEEIEQQYQNRK
    M. incognita NLP-86    (1) --------MRFSIALIVFAFILVIFN--CAESKDEGGNQEEIEQQYQNRR
     M. javanica NLP-86    (1) --------MRFSIALTIFAFILVIFN--CAESKDEGGNQEEIEQQYQNRR
    P. redivivus NLP-86  (101) INMVAARSAQTSSAFNKMLVAMVLVTILAAANAHPFVYGSDIQAPASFNI
    P. univalens NLP-86   (14) RISGIQFQSSTHSVAYGAEMRSWLAVLVVSIIVDVVYPSPLYVAGPSISK
        S. muris NLP-86    (1) -----MRLLLTLLLVSSFCVLISAIPLLDGLKEDSGYVDKR---------
        T. canis NLP-86    (1) -MDPAMLFLESLCEYDAAKRAVSAVNESIENVEFHAHFYCAFADYLKELF
                               151                                            200
 A. lumbricoides NLP-86   (32) VS----------EFASGRWGLRPGKRSQDVPIYTDELGLDGN---SPLYD
         A. suum NLP-86   (58) IR----------SLASGRWGLRPGKRSQDVPIYTDELGLDGN---SPLYD
   B. xylophilus NLP-86   (47) GLRPGKRS----SLASGRWGLRPGKRS--IVDLVLEDDIEDE---SDKTR
       C. sinica NLP-86  (148) VENDNMNNNQKRSLANGRSQWRPGKKS----SINSRVNLGNGMDLIVDAN
      G. pallida NLP-86   (49) IR----------SLANGRWQLRPGKR----ASLIDFRPMDGL---EERAR
 G.rostochiensis NLP-86   (49) IR----------SLANGRWQLRPGKR----ASLIDFRPMDGL---GERAR
     M. arenaria NLP-86   (41) LR----------SLANGRWQLRPGKRFVP-ENYYYQMMLDN---------
  M. floridensis NLP-86   (41) LR----------SLANGRWQLSSPPHLKK-GSHQEKDSCQKT---ITIK-
        M. hapla NLP-86   (43) LR----------SLANGRWQLRPGKRFVP-ESYYYQMMLDN---------
    M. incognita NLP-86   (41) LR----------SLANGRWQLRPGKRFVP-ENYYYQMMLDN---------
     M. javanica NLP-86   (41) LR----------SLANGRWQLRPGKRFVP-ENYYYQMMLDN---------
    P. redivivus NLP-86  (151) MP---KR-----SLASGRWGLRPGKRS-------YDADLDDEDSHFAARF
    P. univalens NLP-86   (64) R-----------SLASGRWGLRPGKRSQDAPNYVDELGFDGN---SPLYD
        S. muris NLP-86   (37) ------------SLASGRWGLRPGKRSLGPLTPSDMFGGNGDGQFLLLPD
        T. canis NLP-86   (50) QR----------SLASGRWGLRPGKRS-SDMAMSYVDDADGI---APLYD
                               201                                            250
 A. lumbricoides NLP-86   (69) ALRQSHFVYVVRK-------------------------------------
         A. suum NLP-86   (95) ALRQSHFVYVVRK-------------------------------------
   B. xylophilus NLP-86   (88) ERRNLASGRWGLRPGKRSLANGRWGLRPGKRSMGMMPTQARHNPIMLLIP
      G. pallida NLP-86   (82) RFRNLYALLPPLNNWN----------------------------------
 G.rostochiensis NLP-86   (82) RFRNLYALLPPLNNWN----------------------------------
     M. arenaria NLP-86   (71) --------------------------------------------------
  M. floridensis NLP-86   (76) --------------------------------------------------
        M. hapla NLP-86   (73) --------------------------------------------------
    M. incognita NLP-86   (71) --------------------------------------------------
     M. javanica NLP-86   (71) --------------------------------------------------
    P. redivivus NLP-86  (186) PYRFVLVD------------------------------------------
    P. univalens NLP-86  (100) ALRQPHFVYLVRK-------------------------------------
        S. muris NLP-86   (75) AYDRPQHLYWWLG-------------------------------------
        T. canis NLP-86   (86) TFRQQPVVYVVSKRK-----------------------------------
                               251
 A. lumbricoides NLP-86   (82) --
         A. suum NLP-86  (108) --
   B. xylophilus NLP-86  (138) QL
      G. pallida NLP-86   (98) --
 G.rostochiensis NLP-86   (98) --
     M. arenaria NLP-86   (71) --
  M. floridensis NLP-86   (76) --
        M. hapla NLP-86   (73) --
    M. incognita NLP-86   (71) --
     M. javanica NLP-86   (71) --
    P. redivivus NLP-86  (194) --
    P. univalens NLP-86  (113) --
        S. muris NLP-86   (88) --
        T. canis NLP-86  (101) --
```

NLP-87 Pan-phylum

Note: additional species *Ascaris lumbricoides, Plectus sambesi, Steinernema carpocapsae, Steinernema glaseri, Steinernema monticolum, Steinernema scapterisci, Toxocara canis.*

```
                              1                                               50
A. lumbricoides NLP-87    (1) --------------------------------------------------
         A.suum NLP-87    (1) --------------------------------------------------
     P. sambesi NLP-87    (1) MQVAHEWDEDGEIALFYFCVYDVFESRRPPRRADDNDAQALIICLQQTLP
 S. carpocapsae NLP-87    (1) --------------------------------------------------
     S. glaseri NLP-87    (1) ------------------------------MLRKEWEGRHAAFLDRVCEC
  S. monticolum NLP-87    (1) --------------------------------------------------
 S. scapterisci NLP-87    (1) --------------------------------------------------
       T. canis NLP-87    (1) --------------------------------------------------
                              51                                             100
A. lumbricoides NLP-87    (1) --------------------------------------------------
         A.suum NLP-87    (1) ---------------------------------------MMCTWKCVLLV
     P. sambesi NLP-87   (51) STRRINIAFRLQKAQFHARTARRFAYDDDDDQDNNMAQTLALKLICVAMV
 S. carpocapsae NLP-87    (1) ------------------------------------------------MT
     S. glaseri NLP-87   (21) RTLSERTEAACVSTEGAYNHASYSLLFGGKSGPGDEWYFQCNGEPIETMS
  S. monticolum NLP-87    (1) -----------------------------------------------MSP
 S. scapterisci NLP-87    (1) ------------------------------------------------MT
       T. canis NLP-87    (1) ---------------------------------------MMCIWKCVLVA
                              101                                            150
A. lumbricoides NLP-87    (1) -MVSVLSCDANLAVGRSSLRPSGKRSIALGRFSLRPGKR-----SIESTA
         A.suum NLP-87   (12) LMVSVLSCDANLAVGRSSLRPSGKRSIALGRFSLRPGKR-----SIGSFP
     P. sambesi NLP-87  (101) VALVTEETAANVAIGRSGSVRPGKRSLALGRFNLRPGKR--------SII
 S. carpocapsae NLP-87    (3) RSVLRFWLVLSFFALAFLLVTAEKRSIALGRLSLRPGKRADYSSTAQDIS
     S. glaseri NLP-87   (71) PSLAVFRIIVFLLLVVYVCATYNKRSIALGRLSLRPGKRS-PFSSSDSLE
  S. monticolum NLP-87    (4) SAMTMFQLFVTMLFVTCILANQNKRSIALGRLSLRPGKR--DSTIAPTME
 S. scapterisci NLP-87    (3) RCVLIVRLVLSLLALTLLLVTAEKRSIALGRLSLRPGKRSGYSSTAQDLN
       T. canis NLP-87   (12) LIVAVMFSEANLAVGRSSLRPSGKRSIALGRFSLRPGKR-----SVESAV
                              151                                            200
A. lumbricoides NLP-87   (45) QSLQTILGDKSGWDCSTQQLAMISDVMEGLASLLEDYTSYLGRCSALADI
         A.suum NLP-87   (57) FK------------------------------------------------
     P. sambesi NLP-87  (143) DSINAIDVDTNS--CNPSDIADVTNVMENLVPHLEKYVVYLERCTDGNSD
 S. carpocapsae NLP-87   (53) DVLQFIRVPSVAPFCSGALVEAVTTNLEAVIRLLDKYSTYLERCNELGYD
     S. glaseri NLP-87  (120) DLLNVVRLPAVAAQCSNAHVETITGGLKNLARLLDEYSFYLEKCRDLGYS
  S. monticolum NLP-87   (52) DLIQVVRVPTIVSQCSNVHMEAVANTLDNLIRLMEQYSFYLEKCSDLGYS
 S. scapterisci NLP-87   (53) DMLHFIRVPSVAPVCSNAQVEAVTRNLEAVIHLMDKYSTYLERCYELGYD
       T. canis NLP-87   (57) QSLERMLQEKTEWECSTEQLGMMNNLMERAASLLEEYTSFLNACSLLSDI
                              201
A. lumbricoides NLP-87   (95) AGNFR
         A.suum NLP-87   (59) -----
     P. sambesi NLP-87  (191) K----
 S. carpocapsae NLP-87  (103) IPI--
     S. glaseri NLP-87  (170) T----
  S. monticolum NLP-87  (102) V----
 S. scapterisci NLP-87  (103) YFM--
       T. canis NLP-87  (107) GSQL-
```

NLP-88

```
                 1                                               50
Ce-NLP-88    (1) -------------------------------------------MSILLIL
As-NLP-88    (1) ---------------------------------------MKPTDKCLLFI
Na-NLP-88    (1) -----------------------------------------MLGPLLLSV
Hc-NLP-88    (1) MGPFSPPFSRDWNPPTGGAVPEENSFPGLLNFPIAKKQEGKMISPLLLSL
Bx-NLP-88    (1) -----------------------------------MKGLDMQIKHVLLFF
Gp-NLP-88    (1) --MLSLPALSLVSFSENGRFRNFCLLLATLCLISGGRSLPLDRTNDRLFF
                 51                                             100
Ce-NLP-88    (8) LFCVVATSALVQLP--------RFSRYGIITNYDSPSRE---EMIVER--
As-NLP-88   (12) VCLIAIIGAGNATP--------TDEINLSYLSSSLNGLHPLATWDMKNER
Na-NLP-88   (10) AAFVPVLSSIISSS--------QDSQLMNLLASTRPASEGYPEWILQSSR
Hc-NLP-88   (51) ATLVPIVSSILSSS--------QDSQLMNLLASSRGASDGYPEWILQSSK
Bx-NLP-88   (16) LITMASVRTVQSASGSIYYYRDPPSAVLNPLSRFFYAEE---PRHLSPAK
Gp-NLP-88   (49) STVLRNPHSVNTLIPIG-----GVSPMFRFVGTNQFVGLKKRRERVGPIQ
                 101                                            150
Ce-NLP-88   (45) ----KRYDRNCFFSPVQCMLTYNDDSR------LPLVVTRKSSKRSIPLF
As-NLP-88   (54) RPIGKKFDRNCFFSPVQCMLSLNSNPA------DDMIWVRRK--------
Na-NLP-88   (52) A--VKKYDRNCFFSPVQCMLSFNQGQE------QPIVFTSKG-KR----L
Hc-NLP-88   (93) A--VKKYDRNCFFSPVQCMLSFNQGQE------QPIVFTSKG-KR----L
Bx-NLP-88   (63) R--DKKYDRNCFFSPVQCMLSYNTNNN------DALYHPRGR-K------
Gp-NLP-88   (94) PFLAKKYDRNCFFSPVQCMLSFKDGSNSMGSNSEVLHFEAGRRRRR----
                 151
Ce-NLP-88   (85) SDFRNLK
As-NLP-88   (90) -------
Na-NLP-88   (89) FYYRK--
Hc-NLP-88  (130) YYYRK--
Bx-NLP-88   (98) -------
Gp-NLP-88  (140) -------
```

NLP-92

```
                1                                               50
As-NLP-92   (1) ----------------------------IIALKLYCDFFIEATNCRPYAP
Bm-NLP-92   (1) MVLALRKQTSPIFLSAVFLLCFPICNHATLLTHIIHGRLFDIDPSSLFAR
Di-NLP-92   (1) MVLTSWKKIFTYSPISAISTMLSHLQSCNIPNTYCSQQIFDTDPLSLIGR
                51                                            98
As-NLP-92  (23) YPPVAPLTKLRSGA-LRNCFFSPVQCLLPFNNKSMRRLDEKRTAFNE-
Bm-NLP-92  (51) ELKLIHLKKQRSNN-LRNCFFSPVQCLLPFDDGRSVESEAKRRS----
Di-NLP-92  (51) ELKLTYQKKLRSNNNLRKCFFSPVQCLLPFDCSRSMESDNKKRIGSYD
```

NLP-92 Pan-phylum

Note: additional species *Ascaris lumbricoides, Anisakis simplex, Acanthocheilonema viteae, Brugia phangai, Brugia timori, Elaeophora elaphi, Gongylonema pulchrum, Onchocerca flexuosa, Onchocerca ochengi, Onchocerca volvulus, Toxocara canis, Wuchereria bancrofti*.

```
                              1                                               50
A. lumbricoides NLP-92    (1) MKAMSSWITGSMLWLAVLLLLDGQHAESLPTVDFEDARFIDGTEFNARNA
     A. simplex NLP-92    (1) --------------------------------------------------
        A. suum NLP-92    (1) --------------------------------------------------
      A. viteae NLP-92    (1) --------------------------------------------------
      B. malayi NLP-92    (1) ----------------------------------------MVLALRKQTS
     B. phangai NLP-92    (1) --------------------------------------------------
      B. timori-NLP-92    (1) --------------------------------------------------
     D. immitis NLP-92    (1) ----------------------------------------MVLTSWKKIF
      E. elaphi NLP-92    (1) --------------------------------------------------
    G. pulchrum NLP-92    (1) --------------------------------------------------
    O. flexuosa NLP-92    (1) ----------------------------------------MVLTSWKQSS
     O. ochengi NLP-92    (1) --------------------------------------------------
    O. volvulus NLP-92    (1) ----------------------------------------MVLTLWKQSS
       T. canis NLP-92    (1) -----------------------------MNNDNPKVAGRAGHSNNSK-V
    W. bancrofi NLP-92    (1) --------------------------------------------------
                              51                                             100
A. lumbricoides NLP-92   (51) NLTHTEGANREGNTSKRVSILYCDFFIEATNCRPYAPYPPVAP---LTKL
     A. simplex NLP-92    (1) -----------------MLSVVCSFLEKRLYIGSYRSYGPRFYP-LLNKR
        A. suum NLP-92    (1) ---------------IIALKLYCDFFIEATNCRPYAPYPPVAP---LTKL
      A. viteae NLP-92    (1) ----------------MLCDKTPKTFYRLFDIDPSSLFTKEFKLIQSTKQ
      B. malayi NLP-92   (11) PIFLSAVFLLCFPICNHATLLTHIIHGRLFDIDPSSLFARELKLIHLKKQ
     B. phangai NLP-92    (1) ------------------------MFYRLFDIDPSSLFARELKLIHLKKQ
      B. timori-NLP-92    (1) ---------------------------RLFDIDPSSLFARELKLIHLKKQ
     D. immitis NLP-92   (11) TYSPISAISTMLSHLQSCNIPNTYCSQQIFDTDPLSLIGRELKLTYQKKL
      E. elaphi NLP-92    (1) ------------------------------------------------KQ
    G. pulchrum NLP-92    (1) ---------------------YNLICYRTFGYYPSLPVD---DLEYVVKR
    O. flexuosa NLP-92   (11) PLLLSALLLICFPICNHATLLTHIVRGRSFDVDPLSIVGRELKVIQPEKQ
     O. ochengi NLP-92    (1) ---------------------------RSFDVDPLSIVGRELKVIHPEKQ
    O. volvulus NLP-92   (11) PLLLSAVLLICFPICSHATLLTHIVRGRSFDVDPLSIVGRELKVIHPEKQ
       T. canis NLP-92   (21) MSSWILGSVVWTAMLLILTDQQTNAFPTMILDDPRPYMPPNLDGLIVPKR
    W. bancrofi NLP-92    (1) ---------------------------RLFDIDPSSLFARELKLIHLKKQ
                              101                                138
A. lumbricoides NLP-92   (98) RS-GALRNCFFSPVQCLLPFNN-KSMRRLDEKRTAFNE
     A. simplex NLP-92   (33) RN-AALRNCFFSPVQCLLPFDN-RSLRRFTDQSITDFD
        A. suum NLP-92   (33) RS-GALRNCFFSPVQCLLPFNN-KSMRRLDEKRTAFNE
      A. viteae NLP-92   (35) RSKNNLRNCFFSPVQCLLPFDDSRLMQSNSKRRTEFPD
      B. malayi NLP-92   (61) RS-NNLRNCFFSPVQCLLPFDDGRSVESEAKRRS----
     B. phangai NLP-92   (27) RS-NNLRNCFFSPVQCLLPFDDGRSVESDAKRRS----
      B. timori-NLP-92   (24) RS-NNLRNCFFSPVQCLLPFDDGRSVESDAKRRS----
     D. immitis NLP-92   (61) RSNNNLRKCFFSPVQCLLPFDCSRSMESDNKKRIGSYD
      E. elaphi NLP-92    (3) QNMNALRNCFLSPVQCLLPFD-----------------
    G. pulchrum NLP-92   (27) RSKNSLRNCFFSPVQCLLPFDNSRSMRRIGEKRAENYD
    O. flexuosa NLP-92   (61) RSSNNLRNCFFSPVQCLLPFDSSRSMENDIKKRIEFFY
     O. ochengi NLP-92   (24) RS-NNLRNCFFSPVQCLLPFDSSRSMENDIKKR-----
    O. volvulus NLP-92   (61) RS-NNLRNCFFSPVQCLLPFDSSRSMENDIKKRIEFFY
       T. canis NLP-92   (71) RD-GGLRNCFFSPVQCLLPFDN-KSLRRFGEKRTIGFQ
    W. bancrofi NLP-92   (24) RS-NNLRNCFFSPVQCLLPFGDGRSVESNTKRRS----
```

NLP-93

```
                 1                                               50
As-NLP-93    (1) ---------------------------------LYALRLDGNLSK-----
Na-NLP-93    (1) -----------------------------MSHIKRVTMCCFRFRK-----
Hc-NLP-93    (1) --------MLSLRSLATIFFVLVVVASSQRDHLSYYSQPDLRSSS-----
Bx-NLP-93    (1) MQMLVILLLLNFISANPALRSMPQLNPVMRYYRLKSQPADENSLPNERPV
                 51                                             100
As-NLP-93   (13) RFRKQSRNSRNCFFSPIQCHLPVAAINDMETLSKRRFRSKPW--------
Na-NLP-93   (17) --ESDSQKYRNCYFSPVQCQLPVLINIDPLASAIKPDLQQDIYTRFRRYR
Hc-NLP-93   (38) EYTKRRAALRNCFLSPMQCLLPVDNLRFNKFHKRFFFAK-----------
Bx-NLP-93   (51) RATPLASSGRNCFFSPVQCRLPVSKVRSMDSSGQRLFLNDQPGLGIFDRM
                 101
As-NLP-93   (55) ------
Na-NLP-93   (65) K-----
Hc-NLP-93   (77) ------
Bx-NLP-93  (101) RRWNKY
```

NLP-93 Pan-phylum

Note: additional species *Ascaris lumbricoides, Anisakis simplex, Ditylenchus destructor, Haemonchus placei, Heligmosomoides polygyrus, Oscheius tipulae, Pristionchus exspectatus, Pristionchus pacificus, Plectus sambesii, Steinernema glaseri, Toxocara canis*.

```
                              1                                                         60
A. lumbricoides NLP-93    (1) --------------------------------MAQSLMMGYERLFLLSIFISAVICNESS
     A. simplex NLP-93    (1) ------------------------------------------------------------
        A. suum NLP-93    (1) --------------------------------------------------LYALRLDG--
  B. xylophilus NLP-93    (1) --------------------------MQMLVILLLLNFISANPALRSMPQLNPVMRYYRL
  D. destructor NLP-93    (1) -------MALSSKMLWDSVRWSHVITAIFVLTTFIVICPTTSVASPLARYQLGRSSTDFA
   H. contortus NLP-93    (1) ------------------------------MLSLRSLATIFFVLVVVASSQRDHLSYYSQ
     H. placeii NLP-93    (1) -----------------------------------------------------FSRLNSP
   H. polygyrus NLP-93    (1) --------------------------MKFVQLIALLCFLSPLYSRVIQGDANYYYKVLRM
  N. americanus NLP-93    (1) ------------------------------------------------------MSHIKR
     O. tipulae NLP-93    (1) -----------------------------MRKDLEGCDIRVHLTEVCFRYWAQNRLRRKL
 P. exspectatus NLP-93    (1) ------------------------------------------------------------
   P. pacificus NLP-93    (1) ---------------------MSSSSSSTISSLLFVLISLSLLLSTAQAYPSYVYADDGR
     P.sambesii NLP-93    (1) MTANSERVFILLAVVAVVGVFAEIQTSDTQQRLNNYDQRDMTRLYRSMSKIGAIKDINEW
     S. glaseri NLP-93    (1) ---MSLDRLGNVGPNWLAVLLMISLLHEGLSTSPKNTASLIESMNPIVRYRFTAHLKNRL
       T. canis NLP-93    (1) ------------------------------------------------------------
                              61                                                       120
A. lumbricoides NLP-93   (29) DQFEDMNPILRKQ----------SRNSRNCFFSPIQCHLPVAAIND-------METLSKR
     A. simplex NLP-93    (1) -----MLKSFRKQ----------SKNSRNCFFSPVQCQLPVPAIDQ-------MENLSKR
        A. suum NLP-93    (9) ----NLSKRFRKQ----------SRNSRNCFFSPIQCHLPVAAIND-------METLSKR
  B. xylophilus NLP-93   (35) KSQPADENSLPNERPV--RATPLASSGRNCFFSPVQCRLPVSKVRSMDSSGQRLFLNDQP
  D. destructor NLP-93   (54) PTWDVILVKRAKNNMSPSQIAPSIRNNRNCYFSPVQCLLPVNTDVLR------SFSSDRY
   H. contortus NLP-93   (31) PDLRSSSEYTKRR-----------AALRNCFLSPMQCLLPVDNLRFN--------KFHKR
     H. placeii NLP-93    (8) PSPPSHSEAASRL----------LSLDRNCFFSPVQCMLPHNSMQQ-------TI-----
   H. polygyrus NLP-93   (35) TNPGASHSIASRA----------PSLDRNCFFSPVQCMLPVTMQHP-------I------
  N. americanus NLP-93    (7) VTMCCFRFRKESD----------SQKYRNCYFSPVQCQLPVLINIDPLASAIKPDLQQDI
     O. tipulae NLP-93   (32) PLLDAMAAKNRQEG-------GDSVKYRNCFFSPVQCQLPVTVDKDPLVTALHIPNQAAF
 P. exspectatus NLP-93    (1) ---YFRMTTPRKP----------EFLRRNCFFSPVQCQFPVSQGVN-------ELRFRRK
   P. pacificus NLP-93   (40) SDFNVPVVRRSPS----------HSAMRNCFFSPVQCMMPMQSVNFN--------GFHKR
     P.sambesii NLP-93   (61) KTKTANAPNKNLPPLQPSLADLSQGNIRNCYFSPVQCLLPVGKRSDAFPKLDFSTDLSRF
     S. glaseri NLP-93   (58) QQLDSNSADRQKP----------NRNSRNCFFSPVQCQLPVEVASQQRPDNLYVVNKFRK
       T. canis NLP-93    (1) ------NFNFRKQ----------SLNSRNCFFSPVQCQLPVTAIDD-------METLSKR
                              121           137
A. lumbricoides NLP-93   (72) RFRSKPW----------
     A. simplex NLP-93   (39) WQWMRPS----------
        A. suum NLP-93   (48) RFRSKPW----------
  B. xylophilus NLP-93   (93) GLGIFDRMRRWNKY---
  D. destructor NLP-93  (108) -----------------
   H. contortus NLP-93   (72) FFFAK------------
     H. placeii NLP-93   (46) -----------------
   H. polygyrus NLP-93   (72) -----------------
  N. americanus NLP-93   (57) YTRFRRYRK--------
     O. tipulae NLP-93   (85) DRFRRLGWGRK------
 P. exspectatus NLP-93   (41) L----------------
   P. pacificus NLP-93   (82) LN---------------
     P.sambesii NLP-93  (121) LKFLGARNSIDSE----
     S. glaseri NLP-93  (108) RLSSLPFFSASALGLVD
       T. canis NLP-93   (38) RLRAKPW----------
```

NLP-94 Pan-phylum

Note: additional species *Ascaris lumbricoides, Anisakis simplex, Toxocara canis*.

```
                             1                                               50
A. lumbricoides NLP-94   (1) ---------MHGNNRIGVTVAFVICSHIFGN-AMLITHENVMKPFAIGAI
     A. simplex NLP-94   (1) --------MYYNKQQIKIILAFIICAIGYGTNALPLTSMYMLRPLELSEM
        A. suum NLP-94   (1) MNECMHCRFMHGNNRIGVTVAFVMCSHIFGN-AMLITHENVMKPFAIGAI
       T. canis NLP-94   (1) -----------------MSITLLLKTQLHAI-TQRKVCFNKRATLGINNM
                             51                                           97
A. lumbricoides NLP-94  (41) EEQPMNLAYEDNSES---MNAEENAFRRNCFFSPVQCMFRRSTT---
     A. simplex NLP-94  (43) NDNAVSLDYNEDELTNSSLEGASLTFKRNCFFSPVQCMFRRSSSK--
        A. suum NLP-94  (50) EEQPMNLAYEDNSES---MNAEENAFRRNCFFSPVQCMFRRSTT---
       T. canis NLP-94  (33) EERPMHIAYDDDLTP---FYVKSDSFRRNCFFSPVQCMFRRSVDYSM
```

NLP-95

```
                 1                                                         60
As-NLP-95    (1) ------------------------------------------------------------
Bx-NLP-95    (1) ------------------------------------------------------------
Gp-NLP-95    (1) MIRRNSSPISAHFLANFGTSMAAFLLIVVVACSLASHPFTTGTYRYVPKHNFGALYDGGR
                 61                                                       120
As-NLP-95    (1) ---------------------------------------------------------GAL
Bx-NLP-95    (1) ---MLTYCRFVILIVGVNVISLSAQSRAATWKSALNFGPDFEKDWIQMGESSEKREPNAP
Gp-NLP-95   (61) PHLPRSASTFFASSQIPDRTVSSLFGAVPDYLRHAGGVSPSDLSGVQLFVPETIAADEMP
                 121                                                      180
As-NLP-95    (4) IPFQNILQEMFSDNEGEIVEESSRLGNKNCFFSLVQCSFYYR------------------
Bx-NLP-95   (58) RPKPEIAKESLRGMMSDEPIGSGKLASKNCFFSPVQCSFFYKRNYDPFPYSRFV------
Gp-NLP-95  (121) QIGGAELKQNDLPSTDLHGDQKGQLARKNCFFSPLQCSFYYKKRSHHGRNAPGGAIFAPG
                 181
As-NLP-95   (46) --------
Bx-NLP-95  (112) --------
Gp-NLP-95  (181) TVIRSDLF
```

NLP-95 Pan-phylum

Note: additional species *Ascaris lumbricoides, Anisakis simplex, Ditylenchus destructor, Globodera rostochiensis, Meloidogyne arenaria, Meloidogyne floridensis, Meloidogyne hapla, Meloidogyne incognita, Meloidogyne javanica, Parascaris equorum, Panagrellus redivivus, Rhabditophanes spp., Steinernema monticolum, Steinernema feltiae, Toxocara canis*.

```
                                 1                                               50
   A. lumbricoides NLP-95    (1) --------------------------------------------------
        A. simplex NLP-95    (1) --------------------------------------------------
           A. suum NLP-95    (1) --------------------------------------------------
     B. xylophilus NLP-95    (1) --------------------------------------------------
     D. destructor NLP-95    (1) --------------------------------------------------
        G. pallida NLP-95    (1) ------MIRRNSSPISAHFLANFGTSMAAFLLIVVVACSLASHPFTTGTY
  G. rostochiensis NLP-95    (1) ------MNRRNSSPISAHFLANFGTSITAFFLIVVVACSMASHPFTTGTY
       M. arenaria NLP-95    (1) MSTKQQQQKSFSSSSINYSTQIFFYFRIIFIYYFLAIQKLDAYPNFYNNE
    M. floridensis NLP-95    (1) MSTKQQQQKAFSSSSINYSTQIFFYFRIIFIYYFLTIQKLDAYPNFYNNE
          M. hapla NLP-95    (1) --------------------------------------------------
      M. incognita NLP-95    (1) MSTKQQQQKSFSSSSINYSTQIFFYFRIIFIYYFLAIQKLDAYPNFYNNE
       M. javanica NLP-95    (1) MSTKQQQQ-FSSSSSIKYSTQIFFYFRIIFIYYFLAIQKLDAYPNFYNNE
        P. equorum NLP-95    (1) --------------------------------------------------
      P. redivivus NLP-95    (1) --------------------------------------------------
Rhabditophanes sp. NLP-95    (1) --------------------------------------------------
     S. monticolum NLP-95    (1) --------------------------------------------------
        S. feltiae NLP-95    (1) --------------------------------------------------
          T. canis NLP-95    (1) --------------------------------------------------
                                 51                                             100
   A. lumbricoides NLP-95    (1) ----------------------------------------------MFQQ
        A. simplex NLP-95    (1) --------------------------------------------------
           A. suum NLP-95    (1) --------------------------------------------------
     B. xylophilus NLP-95    (1) ----------------------------------MLTYCRFVILIVGVNV
     D. destructor NLP-95    (1) --------------------------------------------------
        G. pallida NLP-95   (45) RYVPKHNFGALYDGG------------RPHLPRSASTFFASSQIPDRTVS
  G. rostochiensis NLP-95   (45) RYVPKHNFGALYDGG------------RPHLPRSASTFFASSQTPYRTVN
       M. arenaria NLP-95   (51) LLPMQNMRKFLSQGGGEEKNEGENNGNLNNLKEENKLNDGNHPVFFGRKI
    M. floridensis NLP-95   (51) LP-IQNVRKFLSQGG-EEKNEGENNGNFNNLKEENKLNDGNHPVFFGRKI
          M. hapla NLP-95    (1) -----------------------------------------------XPI
      M. incognita NLP-95   (51) LLPMQNMRKFLSQGGGEEKNEGENNGNLNNLKEENKLNDGNHPVFFGRKI
       M. javanica NLP-95   (50) LLPIQNVRKFLSQGG-EEKNEGENNGNLNNLKEENKLNDGNHPVFFGRKV
        P. equorum NLP-95    (1) --------------------------------------------------
      P. redivivus NLP-95    (1) -------------------------------MQQYVLLIAAAFIFVVASE
Rhabditophanes sp. NLP-95    (1) ----------------------MLCGIKMIVTMMFFSLMGSALTINHSRL
     S. monticolum NLP-95    (1) --------------MSPNQLVQFALVLVGFVCLTTLAQIDESDSSRVAID
        S. feltiae NLP-95    (1) --------------------------------------------------
          T. canis NLP-95    (1) -----------------------------MLYHAAVMLPLIASLLSLSEQ
                                 101                                            150
   A. lumbricoides NLP-95    (5) INFDKCIKRAKKEAILPVISCSNLYKKRTGLQNEINRFQ----------D
        A. simplex NLP-95    (1) ------------------------------MGMSEWDDK----------N
           A. suum NLP-95    (1) --------------------------------GALIPFQ----------N
     B. xylophilus NLP-95   (17) ISLSAQSRAATWKSALNFGPDFEKDWIQMGESSEKREPN-----APRPKP
     D. destructor NLP-95    (1) -------------------------------------------------M
        G. pallida NLP-95   (83) SLFGAVPDYLRHAGGVSPSDLSGVQLFVPETIAADEMP--------QIGG
  G. rostochiensis NLP-95   (83) SLFGAVPDYLRHAGSVSPSDLSGVQLFVPETIPADEMP--------QIGG
       M. arenaria NLP-95  (101) SSLGDLQQTFTEGLLETPSSGFVRSAANKLIKKDKEEN----------GI
    M. floridensis NLP-95   (99) SSLGDLQQTFTE--LETPSSGFVRSAANKLIKKDKEEN----------GI
          M. hapla NLP-95    (4) SSLGDLQQTFTEGLLETPSSGFVRSAANKLMKKAEKEE---KENKKENEI
      M. incognita NLP-95  (101) SSLGDLQQTFTEGLLETPSSGFVRSAANKLIKKDKEEN----------GI
       M. javanica NLP-95   (99) SSLGDLQQTFTE--LETPSSTFVRSAANKLIKKDKEEIE-EKENKKENGI
        P. equorum NLP-95    (1) --------------------------------------------------
      P. redivivus NLP-95   (20) SVQNSPVDFRPYRRFNDLESQFEENLLAPLVKRPFERRDGAVIGARRPSF
Rhabditophanes sp. NLP-95   (29) NRMHIRGIIEQEARIEQLRDSLSNNYLSQYNSNSAPRSDP--PKRTMSLD
     S. monticolum NLP-95   (37) PAALARYRIYAYGNLLKRSDEGSLG-TSKLSFFTDKRRA----------D
        S. feltiae NLP-95    (1) --RHQNPLTNDYRNLLKRSAEGQLGSPAKLSLFTDKRRA----------D
          T. canis NLP-95   (22) KSIAMTEIRNSEPLFNSLSYLSSLYNKRTGIQSEASRVQ----------Q
                                 151                                            200
   A. lumbricoides NLP-95   (45) ILRGSKRKDNEGEIVEESSRLGNKNCFFSLVQCSFYYR------------
        A. simplex NLP-95   (11) GRCIAHISENDAEIMESSSRFGNKNCFFSLVQCSFYY-------------
           A. suum NLP-95    (9) ILQ-EMFSDNEGEIVEESSRLGNKNCFFSLVQCSFYYR------------
     B. xylophilus NLP-95   (62) EIAKESLRGMMSDEPIGSGKLASKNCFFSPVQCSFFYKRNYDPFPYSRFV
     D. destructor NLP-95    (2) PEFIARDSFLRDETIKPTSRLARKNCFMSPLQCSFYYKRSNSFF------
        G. pallida NLP-95  (125) AELKQNDLPSTDLHGDQKGQLARKNCFFSPLQCSFYYKKRSHHGRNAPGG
  G. rostochiensis NLP-95  (125) AELKQNDLPLTDLHGDQKGQLARKNCFFSPLQCSFYYKKRSHHRRSAPGG
       M. arenaria NLP-95  (141) INTIITNNRQPRAN-NYNSKLARKNCFFSPLQCSFYYKSNSNFANMLMRG
    M. floridensis NLP-95  (137) INTLITNNRQPRANSNYNSKLARKNCFFSPLQCSFYYKSNSNFANMLMRG
          M. hapla NLP-95   (51) INTIITNNRQPRAN---NAKLARKNCFFSPLQCSFYYKSNSNFANMLIRG
      M. incognita NLP-95  (141) INTIITNNRQPRAN-NYNSKLARKNCFFSPLQCSFYYKSNSNFANMLMRG
       M. javanica NLP-95  (146) INTIITNNRQPRAN-NYNSKLARKNCFFSPLQCSFYYKSNSNFANMLMRG
        P. equorum NLP-95    (1) ----EMFSDNEGEIADDSSRLGSKNCFFSLVQCSFYYR------------
      P. redivivus NLP-95   (70) VKSGIVRDIVNDEAGKSSSRFARKNCFFSPVQCSFYYKRSVNLF------
Rhabditophanes sp. NLP-95   (77) AFSAYHDLSQGPYVQPSGSKYSTKNCFLSPVQCSFFYYRK----------
     S. monticolum NLP-95   (76) GRSSAGLQPGSIRSFSGTPKFSSKNCFFSPVQCSFYYRR-----------
        S. feltiae NLP-95   (39) GRGGAGLQPGSIRSFSGTPKFSSKNCFFSPVQCSFYYRR-----------
          T. canis NLP-95   (62) ILSGAKRKDSESEIVEGRTRFGNKNCFFSLVQCSFYYR------------
                                 201        214
   A. lumbricoides NLP-95   (83) --------------
        A. simplex NLP-95   (48) --------------
           A. suum NLP-95   (46) --------------
     B. xylophilus NLP-95  (112) --------------
     D. destructor NLP-95   (46) --------------
        G. pallida NLP-95  (175) AIFAPGTVIRSDLF
  G. rostochiensis NLP-95  (175) AIFTPGTVISSNLF
       M. arenaria NLP-95  (190) DE------------
    M. floridensis NLP-95  (187) DE------------
          M. hapla NLP-95   (98) DE------------
      M. incognita NLP-95  (190) DE------------
       M. javanica NLP-95  (195) DE------------
        P. equorum NLP-95   (35) --------------
      P. redivivus NLP-95  (114) --------------
Rhabditophanes sp. NLP-95  (117) --------------
     S. monticolum NLP-95  (115) --------------
        S. feltiae NLP-95   (78) --------------
          T. canis NLP-95  (100) --------------
```

NLP-96 Pan-phylum

Note: additional species *Ancylostoma caninum, Angiostrongylus cantonensis, Ancylostoma ceylanicum, Angiostrongylus costaricensis, Ancylostoma duodenale, Dictyocaulus viviparus, Heterorhabditis bacteriophora, Haemonchus placei, Heligmosomoides polygyrus, Nippostrongylus brasiliensis, Oesophagostomum dentatum, Strongylus vulgaris, Teladorsagia circumcincta.*

```
                               1                                               50
      A. caninum NLP-96    (1) --------------------------------------------------
  A. cantonensis NLP-96    (1) --------------------------------------------------
   A. ceylanicum NLP-96    (1) --------------------------------------------------
A. costaricensis NLP-96    (1) --------------------------------------------------
    A. duodenale NLP-96    (1) --------------------------------------------------
    D. viviparus NLP-96    (1) --------------------------------------------------
H. bacteriophora NLP-96    (1) --------------------------------------------------
    H. contortus NLP-96    (1) --------------------------------------------------
       H. placei NLP-96    (1) --------------------------------------------------
    H. polygyrus NLP-96    (1) --------------------------------------------------
 N. brasiliensis NLP-96    (1) MEARTEKTVVGSREEFRNNSDIGEAKPSLDHNVVYFVVVQSTWRIRRSSI
     O. dentatum NLP-96    (1) -------------------------GSKIHKIKILLELQLPFCVYRRSPC
     S. vulgaris NLP-96    (1) --------------------------------------------------
 T. circumcincta NLP-96    (1) --------------------------------------------------
                               51                                             100
      A. caninum NLP-96    (1) -------------MLVVPLILLFCVFSQLPARVVVGDANYYYKVLRLANP
  A. cantonensis NLP-96    (1) ---------------------------------------RNWKLFRLASS
   A. ceylanicum NLP-96    (1) -------------MLVVPLIVLLCVFSQLHARVVVGDANYYYKVLRLANP
A. costaricensis NLP-96    (1) -------------------------------------PLVRNDYALGYCS
    A. duodenale NLP-96    (1) ----------------------ALKEQRVGLHRAQFNNFNDSTRFRLANI
    D. viviparus NLP-96    (1) -----------------------------------------YVCFRLTSP
H. bacteriophora NLP-96    (1) ---------------------------------------NLIYLFRMHSP
    H. contortus NLP-96    (1) -------------MRVVQAIAFFCLVSQLYSRVVVGDANYYFKVLRLNSP
       H. placei NLP-96    (1) -------------------------------------------FSRLNSP
    H. polygyrus NLP-96    (1) -------------MKFVQLIALLCFLSPLYSRVIQGDANYYYKVLRMTNP
 N. brasiliensis NLP-96   (51) EDGNSLELQHTKVMSIAQVVLLFCLVSQLSSRVIQGDANYYFKVLRMANP
     O. dentatum NLP-96   (26) SSETQALLETLRRFNIRQYIRIGHRSKLVCAVRFKCLVAYLILHFRMPSP
     S. vulgaris NLP-96    (1) -------------------------------LILSCAQINEFLFFRLADS
 T. circumcincta NLP-96    (1) ---------------------------------------NVRFYRLADPT
                               101                                         147
      A. caninum NLP-96   (38) G---ASHSIASRTPSLDRNCFFSPVQCMLPASGTIPQAI--------
  A. cantonensis NLP-96   (12) N---TPHFIESRNPSLDRNCFFSPVQCMLVN----RSP---------
   A. ceylanicum NLP-96   (38) G---ASHSIASRTPSLDRNCFFSPVQCMLPASGTVPQAI--------
A. costaricensis NLP-96   (14) N---TPHFIESRNPSLDRNCFFSPVQCMLTPPQKHKQPWVIDSGRRL
    A. duodenale NLP-96   (29) G---ASHSIASRTPSLDRNCFFSPVQCMLPASGPIQAI---------
    D. viviparus NLP-96   (10) N---VSHLVASRNPSLDRNCFFSPVQCMFSMSRTP------------
H. bacteriophora NLP-96   (12) G---KAHTIASRTPKLDRNCFFSPVQCMLSSNNALNSF---------
    H. contortus NLP-96   (38) PSPPSHSEPASRLLSLDRNCFFSPVQCMLPHNSMQQTI---------
       H. placei NLP-96    (8) PSPPSHSEAASRLLSLDRNCFFSPVQCMLPHNSMQQTI---------
    H. polygyrus NLP-96   (38) G---ASHSIASRAPSLDRNCFFSPVQCMLPVTMQHPI----------
 N. brasiliensis NLP-96  (101) G---ASHSIASRAPSLDRNCFFSPVQCMLPLNSMQQPL---------
     O. dentatum NLP-96   (76) S---TSHLIASRAPSFDRNCFFSPVQCMLPSNNRLQQAV--------
     S. vulgaris NLP-96   (20) G---LSHYVASRAPSFDRNCFFSPVQCMLPFNRIHRAE---------
 T. circumcincta NLP-96   (12) T---QAHYVASRAPSLDRNCFFSPVQCMLSNGKQAQEI---------
```

NLP-97

```
                 1                                               50
As-NLP-97    (1) -----------MCESGCSGKHHVKAYLARATPLVMFGVYVSK------RS
Bm-NLP-97    (1) ----------------------MCHYHIISYHFIIISIIIIP------TI
Di-NLP-97    (1) -------MLYLMNPNIGNVKMDRYRYSIPFYSIIIISTIIIA------TT
Na-NLP-97    (1) ---------------------------MSSLSFLFVLLLVYT------IS
Hc-NLP-97    (1) ---------------------------MSPLSLLCALLVVYT------AL
Bx-NLP-97    (1) MEHVNHPRVERRGEARGWATPFGGNGGLFARSLIVNCYLDKGRQVGNYST
Gp-NLP-97    (1) -------MSSFTFTSSTTGFIFSALFVSASFAFPPFVGQKAEQDKRGFPV
                 51                                             100
As-NLP-97   (34) RNPYSWMNEEKRMDNPYAWTNVNENMLGLD----LDELEKRARNPYSWMN
Bm-NLP-97   (23) ANPAIGNIYEKPS-KYFTYKNEPDNYGFMP----EEYFNTAKRSRNSYSW
Di-NLP-97   (38) ASPTN-DIYDKSSNHFALYKNEPNNDDSML----DEYFVADKRFQN----
Na-NLP-97   (18) ADAVN----------EIQGTNEDENAIDIN----EEKRVHS-RNPYSWMS
Hc-NLP-97   (18) ADVND----------EAEQHDRDLLQLNDD----GISSVKRARNPYSWMV
Bx-NLP-97   (51) RDSVNGFGDMALLRPSIVVALTLVVVYGLDSFETTKRRLGRPMNPYSWQN
Gp-NLP-97   (44) RNPYSWMAQLQKRSVPSGMDNTSPSDGELPNLLLEEPSKRFPSPAASFGG
                 101                                            150
As-NLP-97   (80) YKEKRALRNPYSWMAFG-----------EKRAPLNPYSWMYTLQKRAPFG
Bm-NLP-97   (68) MSNSAIKSQPRNMLIVL-----------EKPAPLNPYSWQYMMKKKNQI-
Di-NLP-97   (79) --DKGIKLPP--VLIAF-----------EKPVPMNPYSWQYMVKKKSGK-
Na-NLP-97   (53) --QEKRARNPYSWMVS------------EGKRSRNPYSWMAMEKEKYYLN
Hc-NLP-97   (54) VDQSKRSRNPYSWMVHSKGKRSLRYFDDAYRQPRNPYSWLYTMFDKRAR-
Bx-NLP-97  (101) -GGYKIKRSLENELETETGALPE---EKRAPMPRNPYSWLARSQKEGPYQ
Gp-NLP-97   (94) ----RRPQNPYSWMSFGDADQSNMDWLFTRLRPFSELVSKRLNALGMGMG
                 151                                          198
As-NLP-97  (119) --YFRNPYSWLNSPASW-------------------------------
Bm-NLP-97  (106) --ILRNPYSWMNNHNN--------------------------------
Di-NLP-97  (113) --IARNPYSWMNNIY---------------------------------
Na-NLP-97   (89) --YRLLSGQTCASFFR--------------------------------
Hc-NLP-97  (103) ----N-PYSWMNE-----------------------------------
Bx-NLP-97  (147) QKRTRDAYFWNNQDFLLDSLFRSETKKSPYVGFNGRRARNPYSWMNFV
Gp-NLP-97  (140) FRRPKNPYLWAMNRQLLAE-----------------------------
```

NLP-97 Pan-phylum

Note: additional species *Ancylostoma caninum, Ancylostoma ceylanicum, Angiostrongylus costaricensis, Ancylostoma duodenale, Ascaris lumbricoides, Anisakis simplex, Ditylenchus destructor, Dracunculus medinensis, Dictyocaulus viviparus, Enterobius vermicularis, Heterorhabditis bacteriophora, Haemonchus placei, Heligmosomoides polygyrus, Oesophagostomum dentatum, Pristionchus pacificus, Panagrellus redivivus, Steinernema carpocapsae, Steinernema feltiae, Steinernema monticolum, Syphacia muris, Steinernema scapterisci, Strongylus vulgaris, Toxocara canis, Teladorsagia circumcincta.*

```
                               1                                               50
      A. caninum NLP-97    (1) --------------------------------------------------
   A. ceylanicum NLP-97    (1) --------------------------------------------------
A. costaricensis NLP-97    (1) --------------------------------------------------
    A. duodenale NLP-97    (1) --------------------------------------------------
 A. lumbricoides NLP-97    (1) --------------------------------------------------
      A. simplex NLP-97    (1) --------------------------------------------------
         A. suum NLP-97    (1) --------------------------------------------------
       B. malayi NLP-97    (1) --------------------------------------------------
   B. xylophilus NLP-97    (1) --------------------------------------------------
   D. destructor NLP-97    (1) --------------------------------------------------
      D. immitis NLP-97    (1) --------------------------------------------------
   D. medinensis NLP-97    (1) --------------------------------------------------
    D. viviparus NLP-97    (1) --------------------------------------------------
 E. vermicularis NLP-97    (1) --------------------------------------------------
      G. pallida NLP-97    (1) --------------------------------------------------
H. bacteriophora NLP-97    (1) MLHASPSNSDTTRVDSPIVGYEASTRHMDDAFDHHIPIHAVGPSRPHIQP
    H. contortus NLP-97    (1) --------------------------------------------------
       H. placei NLP-97    (1) --------------------------------------------------
    H. polygyrus NLP-97    (1) --------------------------------------------------
   N. americanus NLP-97    (1) --------------------------------------------------
     O. dentatum NLP-97    (1) --------------------------------------------------
    P. pacificus NLP-97    (1) --------------------------------------------------
    P. redivivus NLP-97    (1) --------------------------------------------------
  S. carpocapsae NLP-97    (1) --------------------------------------------------
      S. feltiae NLP-97    (1) --------------------------------------------------
   S. monticolum NLP-97    (1) --------------------------------------------------
        S. muris NLP-97    (1) --------------------------------------------------
  S. scapterisci NLP-97    (1) -------------------MLKGYVCSKPVLSMRHLSDYVHVELSFSTFF
     S. vulgaris NLP-97    (1) --------------------------------------------------
        T. canis NLP-97    (1) --------------------------------------------------
 T. circumcincta NLP-97    (1) --------------------------------------------------
                               51                                             100
      A. caninum NLP-97    (1) --------------------------------------------------
   A. ceylanicum NLP-97    (1) --------------------------------------------------
A. costaricensis NLP-97    (1) --------------------------------------------------
    A. duodenale NLP-97    (1) --------------------------------------------------
 A. lumbricoides NLP-97    (1) --------------------------------------------------
      A. simplex NLP-97    (1) --------------------------------------------------
         A. suum NLP-97    (1) --------------------------------------------------
       B. malayi NLP-97    (1) --------------------------------------------------
   B. xylophilus NLP-97    (1) -----------------------------------------------MEH
   D. destructor NLP-97    (1) -------------------------------------------------M
      D. immitis NLP-97    (1) --------------------------------------------------
   D. medinensis NLP-97    (1) --------------------------------------------------
    D. viviparus NLP-97    (1) --------------------------------------------------
 E. vermicularis NLP-97    (1) --------------------------------------------------
      G. pallida NLP-97    (1) --------------------------------------------------
H. bacteriophora NLP-97   (51) SKNVRMTMISRDEESVLFAAMKFANTVRRRQYRGQCEDDLLIFLSTKDKV
    H. contortus NLP-97    (1) --------------------------------------------------
       H. placei NLP-97    (1) --------------------------------------------------
    H. polygyrus NLP-97    (1) --------------------------------------------------
   N. americanus NLP-97    (1) --------------------------------------------------
     O. dentatum NLP-97    (1) --------------------------------------------------
    P. pacificus NLP-97    (1) --------------------------------------------------
    P. redivivus NLP-97    (1) --------------------------------------------------
  S. carpocapsae NLP-97    (1) --------------------------------------------------
      S. feltiae NLP-97    (1) ----------------------MSAEHRANPPRLESRTDPRDSAAHSISL
   S. monticolum NLP-97    (1) --------------------------------------------------
        S. muris NLP-97    (1) --------------------------------------------------
  S. scapterisci NLP-97   (32) IFVSSTFGHKLHLMTNFLLFARSKLDIRDEEGPKEPEEATTFHTRFPRHS
     S. vulgaris NLP-97    (1) --------------------------------------------------
        T. canis NLP-97    (1) --------------------------------------------------
 T. circumcincta NLP-97    (1) --------------------------------------------------
                               101                                            150
      A. caninum NLP-97    (1) ---------------------MSPLSILCVLLLVYTISAEVVSEIRGTNE
   A. ceylanicum NLP-97    (1) --------------------------------------------------
A. costaricensis NLP-97    (1) -------------------------------MFHISYFYVLLLVISAAAE
    A. duodenale NLP-97    (1) -----------------------------------------------MEW
 A. lumbricoides NLP-97    (1) ------MLAKFIVVTVALLEFGYSGGNADLSADTVDVAKRSRNPYSWMNE
      A. simplex NLP-97    (1) ------MIQKLVVLFFTMIIVDFTQCSVDESVDESDVSKRSRNPYSWMNN
         A. suum NLP-97    (1) --------MCESGCSGKHHVKAYLARATPLVMFGVYVSKRSRNPYSWMNE
       B. malayi NLP-97    (1) -------------------------------MCHYHIISYHFIIISIIII
   B. xylophilus NLP-97    (4) VNHPRVERRGEARGWATPFGGNGGLFARSLIVNCYLDKGRQVGNYSTRDS
   D. destructor NLP-97    (2) GYSQSSIFRFSTVFVEMFFCIVSIGLLACGGINAAPGSGVREVDKTEVDT
      D. immitis NLP-97    (1) ----------------MLYLMNPNIGNVKMDRYRYSIPFYSIIIISTIII
   D. medinensis NLP-97    (1) --------MMGNILLLYFLLVTRMFYANAKSDVIGDDDKRSRNPYSWMNY
    D. viviparus NLP-97    (1) ------------------------------------MLLLVCIVSVTVNS
 E. vermicularis NLP-97    (1) -----------------------------------------------MEE
      G. pallida NLP-97    (1) ----MSSFTFTSSTTGFIFSALFVSASFAFPPFVGQKAEQDKRGFPVRNP
H. bacteriophora NLP-97  (101) VWTSVGKVTQRYLSTQMINSITISSETHFRNGEYTEGLQYMVDSYTSILN
    H. contortus NLP-97    (1) -----------------------------MSPLSLLCALLVVYTALADVN
       H. placei NLP-97    (1) -----------MLGRDCGCPNRVTAADNGIFDKDLLLTEKQANFLLNELG
    H. polygyrus NLP-97    (1) --------------------------------------------------
   N. americanus NLP-97    (1) ---------------------MSSLSFLFVLLLVYTISADAVNEIQGTNE
     O. dentatum NLP-97    (1) --------------------------------------------------
    P. pacificus NLP-97    (1) ---------------------------------------MARSLFSSLLL
    P. redivivus NLP-97    (1) -------------------MLSQPWILFIFASAIGGAAVIADVPVDETDK
  S. carpocapsae NLP-97    (1) -------MSRPLICCLLALGVAHVLAFEDSPP--SEMIKRPMNPYSWMNK
      S. feltiae NLP-97   (29) ARSIVAQMTRPLICCLLALGVA--LAYDDASSSVSEMAKRPQNPYSWMNK
   S. monticolum NLP-97    (1) -------MTRHLICCLLALSVAQVLSFDDSRSAGADLVKRPQNPYSWMNK
        S. muris NLP-97    (1) -----MVVTSAVLQTGVVIVNLFLIISALEVSGPKTDSKRARNPYSWLEE
  S. scapterisci NLP-97   (82) TTYHLAEMTRPLICCLLALGVANVLAFEDSPPSASEMVKRPLNPYSWMNK
     S. vulgaris NLP-97    (1) --------------------------------------------------
        T. canis NLP-97    (1) -----------------------------------------------MNE
 T. circumcincta NLP-97    (1) -----------------------------MSPLSLLCALLVVYTVLADVN
                               151                                            200
      A. caninum NLP-97   (30) DENSMDIT----------EEKRMHARNPYSWMK-VKRAR-NPYSWMLSEQ
   A. ceylanicum NLP-97    (1) -------------------MVTYEDSKLTAPGLEISSFT-YRAVMNRSRK
A. costaricensis NLP-97   (20) SQLQHPN----------EHVMNEDVAKILPFWTDVKRAR-NPYSWMAVEQ
    A. duodenale NLP-97    (4) KGVRDACQ----------ESVLVDESETSTKSLLLDAQR-ARQEGKKPIQ
 A. lumbricoides NLP-97   (45) EKRIDN-------PYAWTNVNEN--MLGLDLDELEKRAR-NPYSWMNYKE
      A. simplex NLP-97   (45) LKRTDA-------LYQWLPNSENSNFNEIEAETSDKRAIKNPYSWQNYKV
         A. suum NLP-97   (43) EKRMDN-------PYAWTNVNEN--MLGLDLDELEKRAR-NPYSWMNYKE
       B. malayi NLP-97   (20) PTIANPAIG---------NIYEKPS-KYFTYKNEPDNYGFMPEEYFNTAK
   B. xylophilus NLP-97   (54) VNGFGDMALLRPSIVVALTLVVVYGLDSFETTKRRLGRPMNPYSWQNGGY
   D. destructor NLP-97   (52) KKPESPDTQFLVQSGVEAAAFRKRAYELASMFDQYYGSLQNEQSPLTEEH
      D. immitis NLP-97   (35) ATTASPTN----------DIYDKSSNHFALYKNEPNNDDSMLDEYFVADK
   D. medinensis NLP-97   (43) ARVSRSP--AQWSEQADENSIFPWGREFMDAGRLEKKAR-NPYSWMNTMY
    D. viviparus NLP-97   (15) QLQQSDDE----------QMNEERLAKASPIWMNAKRAR-NPYSWMAAEE
 E. vermicularis NLP-97    (4) KRSGSR--VANKADMLAAPLPEEYENMFEESSEMQKRFR-NPYSWMNDLD
      G. pallida NLP-97   (47) YSWMAQLQ--------KRSVPSGMDNTSPSDGELPNLLLEEPSKRFPSPA
H. bacteriophora NLP-97  (151) GEHLEVIS----------NAVSKRARNPYSWME-TKRAR-NPYSWMTEQG
    H. contortus NLP-97   (22) DEAEQHDR----------DLLQLNDDGISS----VKRAR-NPYSWMVVDQ
       H. placei NLP-97   (40) KAGEGADV----------PPPGNGAKKFKSMSFLHCNCH-PASVNIVLET
    H. polygyrus NLP-97    (1) ------------------------------------MAR-KRGRHMPGQC
   N. americanus NLP-97   (30) DENAIDIN----------EEKRVHSRNPYSWMSQEKRAR-NPYSWMVSE-
     O. dentatum NLP-97    (1) -------------------------------MM-DKRAR-NPYSWMATEQ
    P. pacificus NLP-97   (12) LAILAS---------FIISILALDDIEE---EKRGRNP----YSWQNVDK
    P. redivivus NLP-97   (32) TEVSTPYQGPAWLKRIPDNLNAFAPDELPLYKRPVGHVSRNPYSWMATNT
  S. carpocapsae NLP-97   (42) YRVKKS---------HPVDMVNEADEGIM-EDEMEKRAR-NPYAWTETIK
      S. feltiae NLP-97   (77) YRVKKS---------HPDADDIILDGVGGDVEDVEKRAR-NPYAWTVTDE
   S. monticolum NLP-97   (44) YRVKKS---------HPLALDEDVEVLG--EDDIEKRAR-NPYAWTVTDE
        S. muris NLP-97   (46) KRGGFRSEVPNKFDLTAPALPEEYDTLLENTFEIPKRAR-NPYSWMNDLE
  S. scapterisci NLP-97  (132) YRVKKS---------HPVDMANEADEGIM-EEEMEKRAR-NPYAWTETMK
     S. vulgaris NLP-97    (1) -------------------------------MK-VKRTR-NPYSWMASEQ
        T. canis NLP-97    (4) DKRSG--------LYEWLQGNDN--LNDFDIDEMPKRAR-NPYSWMNYME
 T. circumcincta NLP-97   (22) DEVQQQDR----------ELLQLNDDGISSGSA-IKRAR-NPYSWMVTDQ
                               201                                            250
      A. caninum NLP-97   (68) G----------------------KRARNPYSWMVME-------------K
   A. ceylanicum NLP-97   (31) G----------------------DSSLASK---DEL-------------Q
A. costaricensis NLP-97   (59) G----------------------MGGSNPYSWMTQMSKTNFAQIFLYEFK
    A. duodenale NLP-97   (43) L----------------------DGYGEGENCCLDW-------------K
 A. lumbricoides NLP-97   (85) K----------------------RALRNPYSWMA----------------
      A. simplex NLP-97   (88) VNF------------------GFLAPREPYWWMG----------------
         A. suum NLP-97   (83) K----------------------RALRNPYSWMA----------------
       B. malayi NLP-97   (60) R----------------------SRNSYSWMSNSAI-------------K
   B. xylophilus NLP-97  (104) KIKRSLENELETETGALPEEKRAPMPRNPYSWLARS--------------
   D. destructor NLP-97  (102) KRA------------------GSVPSRNPYSWMAVE-------------E
      D. immitis NLP-97   (75) R----------------------FQN------DKGI-------------K
   D. medinensis NLP-97   (90) G------------------------ERDPYGWMAFG-------------G
    D. viviparus NLP-97   (54) V----------------------NSARNPYYWMGQT-------------K
 E. vermicularis NLP-97   (51) K-----------------------RSRNPYSWMN----------------
      G. pallida NLP-97   (89) ASFG------------------GRRPQNPYSWMSFG--------------
H. bacteriophora NLP-97  (189) -----------------------KRARNPYSWLAKE-------------K
    H. contortus NLP-97   (57) S----------------------KRSRNPYSWMVHS-------------K
       H. placei NLP-97   (79) S----------------------ENHGLVWAVSIET-------------Q
    H. polygyrus NLP-97   (14) A----------------------VGPEGGHRSEDES-------------R
   N. americanus NLP-97   (68) G----------------------KRSRNPYSWMAME-------------K
     O. dentatum NLP-97   (18) D----------------------KRARNPYSWMASE-------------K
    P. pacificus NLP-97   (46) R------------------------GRNPYSWMS-G-------------T
    P. redivivus NLP-97   (82) P---------------------SKRSRNPYAWLALA-------------E
  S. carpocapsae NLP-97   (81) ---------------------GKRGARNPYSWLN---------------V
      S. feltiae NLP-97  (117) RFM------------------GKRGARNPYSWMNNN-------------A
   S. monticolum NLP-97   (82) RFM------------------GKRGARNPYSWMN-D-------------A
        S. muris NLP-97   (95) K-----------------------RSRNPYSWMN----------------
  S. scapterisci NLP-97  (171) ---------------------GKRGARNPYSWLN---------------V
     S. vulgaris NLP-97   (18) Y----------------------KRARNPYSWMVSE-------------K
        T. canis NLP-97   (43) K----------------------RSPRNPYSWMA----------------
 T. circumcincta NLP-97   (60) G----------------------KRSRNPYSWMAHP-------------K
                               251                                            300
      A. caninum NLP-97   (83) -DKR---TLRFSDDVYRQPRNPYSWMYTMF--------------------
   A. ceylanicum NLP-97   (43) -DKR---ALRFSDDVYRQPRNPYSWMYTMF--------------------
A. costaricensis NLP-97   (87) LGGSAEKTARNIEEVWGESNAGVSAVQTWFRKFQSG--------------
    A. duodenale NLP-97   (58) -DKR---ALRFSDDVYRQPRNPYSWMYTMF--------------------
 A. lumbricoides NLP-97   (97) ------------FGEKRAPLNPYSWMYTLQ--------------------
      A. simplex NLP-97  (104) ------------YGEKRAPLNPYSWMYTLQ--------------------
         A. suum NLP-97   (95) ------------FGEKRAPLNPYSWMYTLQ--------------------
       B. malayi NLP-97   (75) SQPR---N-MLIVLEKPAPLNPYSWQYMMKK-------------------
   B. xylophilus NLP-97  (140) --------QKEGPYQQKRTRDAYFWNNQDFLLDSLFRSETKKSP------
   D. destructor NLP-97  (121) KRSRNPYSWLAKNDDEKRSRNPYSWLATNDGSIDRTKTWGDAPPPPKRLA
      D. immitis NLP-97   (84) LPP------VLIAFEKPVPMNPYSWQYMVKK-------------------
   D. medinensis NLP-97  (103) ---------------KRAPMNPYSWQYMLA--------------------
    D. viviparus NLP-97   (69) -GKR---LIRYADQIYSHPRNPYSWMYTMF--------------------
 E. vermicularis NLP-97   (62) ------------FGFKRAPMNPYSWQNA----------------------
      G. pallida NLP-97  (107) ------DADQSNMDWLFTRLRPFSELVSKRLNALG---------------
H. bacteriophora NLP-97  (203) GTKR---SPRFAEDVLRQPRNPYSWMYSIF--------------------
    H. contortus NLP-97   (72) -GKR---SLRYFDDAYRQPRNPYSWLYTMF--------------------
       H. placei NLP-97   (94) -GKR---SLRYFDDAYRQPRNPYSWLYTMF--------------------
    H. polygyrus NLP-97   (29) -EKR---AVRFADDIYRQPRNPYSWMYTMF--------------------
   N. americanus NLP-97   (83) -EKY---YLNYRLLSGQTCASFFR--------------------------
     O. dentatum NLP-97   (33) -TKR---SVRFSEEFR-QPRNPYSWMYTMF--------------------
    P. pacificus NLP-97   (58) KRAR---NPYSWMEFQKRARNPYSWME-----------------------
    P. redivivus NLP-97   (98) DSAS---KRAYYPTFDKKSRNPYSWMSDVK--------------------
  S. carpocapsae NLP-97   (95) KPKR---SPYAYFSEQRAPLNPYSWMSSMV--------------------
      S. feltiae NLP-97  (136) KPKR---G-YAFFAEQRMPLNPYSWMSSMV--------------------
   S. monticolum NLP-97  (100) KPKR---S-YAFFAEQRMPLNPYSWMSSMV--------------------
        S. muris NLP-97  (106) ------------FGSKRAPMNPYSWQYA----------------------
  S. scapterisci NLP-97  (185) KPKR---S-YSYFSEQRAPLNPYSWMSSMV--------------------
     S. vulgaris NLP-97   (33) -NKR---SLRFADDGFRQPRNPYSWMYTML--------------------
        T. canis NLP-97   (55) ------------YGEKRAPLNPYSWMYTLQ--------------------
 T. circumcincta NLP-97   (75) -GKR---SLRFNDDGFRQPRNPYSWLYTMF--------------------
                               301                                  340
      A. caninum NLP-97  (109) -----------DKRARNPYSWMNE----------------
   A. ceylanicum NLP-97   (69) -----------DKRARNPYSWMNE----------------
A. costaricensis NLP-97  (123) ------DFDLEDEEGRGCPNELNDDEFDCLQHVMFSKMPS
    A. duodenale NLP-97   (84) -----------DKRARNPYSWMNE----------------
 A. lumbricoides NLP-97  (115) -----------KRAPFGYFRNPYSWLNSPASW--------
      A. simplex NLP-97  (122) -----------KRSAFGRARNPYSWLNTSR----------
         A. suum NLP-97  (113) -----------KRAPFGYFRNPYSWLNSPASW--------
       B. malayi NLP-97  (102) ---------KNQIILRNPYSWMNNHNN-------------
   B. xylophilus NLP-97  (176) ------YVGFNGRRARNPYSWMNFV---------------
   D. destructor NLP-97  (171) RVGPMSWAEGLKRGAKNPYSWMNYA---------------
      D. immitis NLP-97  (109) ---------KSGKIARNPYSWMNNIY--------------
   D. medinensis NLP-97  (118) -----------KRALRNPYSWQMAD---------------
    D. viviparus NLP-97   (95) -----------DKRTNTPYSWMNE----------------
 E. vermicularis NLP-97   (78) ----------------------------------------
      G. pallida NLP-97  (136) -------MGMGFRRPKNPYLWAMNRQLLAE----------
H. bacteriophora NLP-97  (230) -----------EKRTRNPYSWMNE----------------
    H. contortus NLP-97   (98) -----------DKRARNPYSWMNE----------------
       H. placei NLP-97  (120) -----------DKRARNPYSWMNE----------------
    H. polygyrus NLP-97   (55) -----------EKRARNPYSWMNTDA--------------
   N. americanus NLP-97  (103) ----------------------------------------
     O. dentatum NLP-97   (58) -----------DKRARNPYSWMNYE---------------
    P. pacificus NLP-97   (82) ----------------------------------------
    P. redivivus NLP-97  (125) -------------RARNPYAWVNYV---------------
  S. carpocapsae NLP-97  (122) -----------KKSRNPS-GYFYVDERSPVRNPYSWLN--
      S. feltiae NLP-97  (162) -----------KKSRSPSPQYFFVDERSPIRNPYSWQN--
   S. monticolum NLP-97  (126) -----------KKSRTPS-QYFFVDERSPIRNPYSWQN--
        S. muris NLP-97  (122) ----------------------------------------
  S. scapterisci NLP-97  (211) -----------KKSRNPS-GYFYVDERSPVRNPYSWLN--
     S. vulgaris NLP-97   (59) -----------EKRTRNPYSWMNYE---------------
        T. canis NLP-97   (73) -----------KRVPVRYFRNPYSWQSSSASW--------
 T. circumcincta NLP-97  (101) -----------DKRARNPYSWMNE----------------
```

NLP-98 Pan-phylum

```
                                 1                                               50
      G. pallida NLP-98.1    (1) --------------------------------------------------
G. rostochiensis NLP-98.1    (1) ----------------------------------------MIDAASIPLQ
      G. pallida NLP-98.2    (1) ----------------------------------------MIVVALIPLQ
G. rostochiensis NLP-98.2    (1) MIVIALIPLQLMAIAVASALKPFFTLTTLCRLQLCKPSKAMIVIALIPLQ
                                 51                                             100
      G. pallida NLP-98.1    (1) --------------------------------------------------
G. rostochiensis NLP-98.1   (11) LMAIVGAAVLVIIAQCCGCGKK--NDKAEVPPTPAPAEQPDAASKKSQAP
      G. pallida NLP-98.2   (11) LMAIVVAAVLVIITQCCGGGKKNTEEPPQPPQASAVAEQPTTTAAGQSQN
G. rostochiensis NLP-98.2   (51) LMAIAVASALVIITQCCGGGKKNTEEPPPPPQMSAVAEQPTTTAAGQSQN
                                 101                                            150
      G. pallida NLP-98.1    (1) --------------------------------------------------
G. rostochiensis NLP-98.1   (59) TPTAVDQLG---TASKNGPPIATTTGYSTGG-------------------
      G. pallida NLP-98.2   (61) APTSVAEPQ-GATVSKNAPALGATTNYAAPGSTENIFSQSPSTSAAPGAA
G. rostochiensis NLP-98.2  (101) PPTSAVAEQPSPTASKNAPALGATTNYAAPGSTENIFSKMS---------
                                 151                                            200
      G. pallida NLP-98.1    (1) -----------DPGGVSKKSLAP---STSAADPGGVSKKSLAPSTSAADP
G. rostochiensis NLP-98.1   (87) G--------GADMNIFSKMSTAGEPKPPAADLGAVSKKSAIGPTAAAADL
      G. pallida NLP-98.2  (110) SKKSVAPTTSASPGAVSKKSVAP---TTSASPGAVSQKS--VAPTTSASP
G. rostochiensis NLP-98.2  (142) ---AQSPTTSAAPGAVSKKSVAP---TTSAAPGAVSKKS--VAPTTSAAP
                                 201                                          248
      G. pallida NLP-98.1   (37) GGVSKKSLAPSTSAGVTKAQNNQSIALGAPGARSTRI-----------
G. rostochiensis NLP-98.1  (129) GAVSKKSLAPTTSAADLGATVCNSIYIKSSGDESVWMKNLFV------
      G. pallida NLP-98.2  (155) GAVSKKSVAPTTSAS-PAAVSQKSAAPTTSAADPGGVSKK---SSAAP
G. rostochiensis NLP-98.2  (184) DAVSKKSVAPTTSAADPAGVSKKSAAPTTSAADPGGVSKKSAAGSAAP
```
