## Supplementary Table 2 for "*In silico* analyses of neuropeptide-like protein (NLP) profiles in parasitic nematodes"

**Table S2: *Caenorhabditis elegans* Neuropeptide-like protein (NLP) prepropeptide sequences used as BLAST query sequences in this study.**

| NLP | *C. elegans* prepropeptide search string |
| --- | --- |
| 1 | MKATFVLACLLVIAAVSHADLLPKRMDANAFRMSFGKRSVSNPAEAKRMDPNAFRMSFGKRSAEQNEQANKEDKATSDKLYDDTKFEEMKRMDANAFRMSFGKRSDAHQAADDQVEYVNDDFSLPEQKRMDANAFRMSFGKRVNLDPNSFRMSFGKRSTVGYNLDARNYFVGLGRR |
| 2 | MRATLVLFALLCAVYSEAVPLQVYRPDESSAVDVVVLENSPELYDSEDEDEWKQEEEFTEGAMGKRSIALGRSGFRPGKRSMDNFHTVDVSDLIMKRSMAMGRLGLRPGKRSMAYGRQGFRPGKRSMAYGRQGFRPGKRSMAYGRQGFRPGKRSNDMKEVFPQHVPEIYII |
| 3 | MSKIVACLVLLALSVMCVYSAPYEFRAKRAINPFLDSMGKRAVNPFLDSIGKRSFRPDMITEEKRYFDSLAGQSLGKRSNNRYEMLENYY |
| 4.1 | MLVVVLSGIIATCLAVKQTDFSVHIKGQLVCDDVPAANMPMDIFSEAQMFRYEDFMDRWTNEKGEFDFKAYVTTDAYMITPYVVIIHRCWESKEVPHRCDRKIVQPIPPLNVTVNSKTPDEKKVYNFGKVDLRFKNPAEVTVKCG |
| 4.2 | SLILFVILLVAFAAARPVSEEVDRVDYDPRTEAPRRLPADDDEVDGEDRVDYDPRTDAPIRVPVDPEAEGEDRV |
| 5 | MLMKIMVLMGIANIAASFVVSSVRFQSAPMRALIEINRELAKRSVSQLNQYAGFDTLGGMGLGKRSEPDQAGEKRALSTFDSLGGMGLGKRSSSSSRVFVYDKRALQHFSSLDTLGGMGFGRK |
| 6 | MLSFSRLAFVLLVSACVMAMAAPKQMVFGFGKRSMADISDMPEDVPMKRYKPRSFAMGFGKRAAMRSFNMGFGKRSAEPQIDYEDMSADLEEIPVFIQKRRLIMGLGRR |
| 7 | MYIKAALLIVVLFGVASQITSALYLKQADFDDPRMFTSSFGKRSAIESEPQAYPKSYRAIRIQRRSMDDLDDPRLMTMSFGKRMILPSLADLHRYTMYDKRGSDIDDPRYFLFSNRLTCRC |
| 8 | MSQKLLPISPLQLLFLQCLLIGFTAAYPYLIFPASPSSGDSRRLVKRAFDRFDNSGVFSFGAKRFDRYDDETAYGYGFDNHIFKRSADPYRFMSVPTKKAFDRMDNSDFFGAKRKRSFDRMGGTEFGLMKRSAPESREQLINNLAESIITLRRAREAESSPESQRTIITYDD |
| 9 | MDRFATRFIALLLVLLQIGSIFATPIAEAQGAPEDVDDRRELEKRGGARAFYGFYNAGNSKRDQAAALPYYLYEKRGGGRAFNHNANLFRFDKRGGGRAFAGSWSPYLERFYDYKRSSYPVYFSDNSYY |
| 10 | MWYIALLLAVIATSVTAQKADDEPIVFLVRVPIDEMDDDSSLLESYYHPRDILSKRAIPFNGGMYGKRSTMPFSGGMYGKRSGQIFAQRRAAIPFSGGMYGKRSLVPQSYSNNENQIKRGAMPFSGGMYGR |
| 11 | MMSTLALVSLAIFGIAVVCAAPKPATVPVANEEDYLAALYGFEAPGSQFKGAPLQSKRHISPSYDVEIDAGNMRNLLDIGKRSAPMASDYGNQFQMYNRLIDAGKKKRSPAISPAYQFENAFGLSEALERAGRR |
| 12 | MLRHHSCALLMLILVFVEVFATQSPTFDRQDRDYRPLQFGKRDGYRPLQFGKRDYRPLQFGKRSSGSSGPVVLEPIWEWQ |
| 13 | MQRSLQIFCIMSAIAMAYSQGSRDDNQSAKRNDFSRDIMSFGKRSGNTADLYDRRIMAFGKRQPSYDRDIMSFGKRSAPSDFSRDIMSFGKRSSSMYDRDIMSFGKRSPVDYDRPIMAFGKRAEDYERQIMAFGRRK |
| 14 | MLHLIVLLVALSSAVTAGRPRRALDGLDGSGFGFDKRALNSLDGAGFGFEKRALNSLDGQGFGFEKRALDGLDGAGFGFDKRALNSLDGAGFGFEKRALDGLDGSGFGFDKRALNSLDGAGFGFEKRALNSLDGAGFGFEKRALDGLDGAGFGFDKRALNSLDGAGFGFEKRALDGLDGAGFGFDKRALNSLDGNGFGFDKRTFKHSSNKLRSVFRNLKGFKQH |
| 15 | MPSSSSSSSFFAAVLLVIVMMSTVESAAVRLRPVGSLFFLNRPHEKRAFDSLAGSGFDNGFNKRAFDSLAGSGFGAFNKRAFDSLAGSGFGAFNKRAFDSLAGSGFSGFDKRAFDSLAGQGFTGFEKRAFDTVSTSGFDDFKL |
| 16 | MSFRFLILLLLASTLLVNSFPIGKDNESDESEVEVDTTTEAENIEVESQEASLDEQLNEVSEDPTREKRGLYSSERTEEEVEISHGMHHREKRHSEHLPHPDPPSHTAKRSTEHHRVKRSEGHPHEKKATHSPEGHIVAKDDHHGHEKRSSDSHHGHQKRSVDEHHGHQKKNAEDHHEHQKRSEHVEHQAEMHEHQKRSTQEVSGHPEHHLVKRSGGEGGHRHHRSTDQGLDEDEPEDEIQTDENDEVTEEPGSRKRRNTDTPMPSFPSDHDSEHSNSVAIRVKRVSRAGSSHKVRTLNKNRGNSKAGETTQNDSLTPNSGGVFNS |
| 17 | MFSKSILFCLLVLVFNVFGANFENDQDVMRPPFQALKRGSLSNMMRIGKRQMSRQQEYVQFPNEGVVPCESCNLGTLMRIGRR |
| 18 | MNANVYSIVYFLSFLVLCISAQLHADSGATEVDGIVDKRSPYRAFAFAKRSDEENLDFLEKRARYGFAKRSPYRTFAFAKRASPYGFAFAKRGQFSSFA |
| 19 | MLLRGVCLALLILVTIVQCQNDNDLKEEKRRIGLRLPNFLRFKDPDALMIHKRRIGLRLPNMLKFKDSSNMYHLEKRRMGMRLPNIIFLRNEKKNVLEY |
| 20 | MQVTLLALLLLIVPFFAFAASQYSDDDSELMSNNERFARDLELRKKFAFAFAKRSAGDADVVIEARSGPQAHEGAGMRFAFAKRRAPKEFARFARASFA |
| 21 | MRNSLFTTLFFGLAALVMVLNAQYTSELEEDEKRGGARAMLHKRGGARAFSADVGDDYKRGGARAFYDEKRGGARAFLTEMKRGGARVFQGFEDEKRGGARAFMMDKRGGGRAFGDMMKRGGARAFVENSKRDEDWVIRPFEDDRLEKRGGGRSFPVKPGRLDD |
| 22 | MRSIIVFIGLTIFALDILLVQTSALGLQGGIDVFRGLGVVDQVDFNQILHRANYLRNTREGRLRYWRLRTLPIMKKSIAIGRAGFRPGKRTTDELTGFPIGV |
| 23 | MAAHLVIWMALLGVSAHALELGFYSEHPTSKDDSYESQLLDSNDLDQDDAVDLATPMIVIPNDEDEIYADEEPEPLTMEKRLYISRQGFRPAKRSMAIGRAGMRPGKRAFAAGWNRGKRSMPFAESYYPLYRNDFSE |
| 35 | MPRVSSFIVFLTFMVALLAVSHAAVVSGYDNIYQVLAPRFRRARMMSFDAEAPQYLQHLLQNLKPRFRRSV |
| 36 | MSVDLKQQLELADYLGALAVWCIFFGVLFILSVIFNFVCIKKDDDVTALERWGYKKNIDMKLGPHRRSMVARQIPQTVVADH |
| 37 | MSSRISVSLLLLAVVATMFFTANVVDATPRSQGNMMRYGNSLPAYAPHVLYRFYNSRQFAPINKRNNAEVVNHILKNFGA  LDRLGDVGK |
| 38 | MQLIHFIVGLAMLISLSLAASDDRVLGWNKAHGLWGKRSVQEASQDKRTPQNWNKLNSLWGKRSASSFDDDYTTENGDDDVTMLYKRSNLSPRFLGRMTFARIPKISPAQWQRANGLWGR |
| 39 | MKLLILFSLFAIFFGVIALDSPIPEFYSTGRSTRVPSHRHRIPRLGKREVPNFQADNVPEAGGRVRRYVGPPLKLVILI |
| 40 | > isoform a  MKLVILLSFVATVAVFAAPSAPAGLEEKLRALQEQLYSLEKENGVDVKQKEQPAAADTFLGFVPQKRMVAWQPMKRSMINEDSRAPLLHAIEARLAEVLRAGERLGVNPEEVLADLRARNQFQ  >isoform b  MVAWQPMKRSMINEDSRAPLLHAIEARLAEVLRAGERLGVNPEEVLADLRARNQFQ |
| 41 | >isoform a  MLGLVCKILVVLCLSVLCLTVSAAPGLFELPSRSVRLIRSDPSAYDGYENSFYRGYGSDNQQFRFNSPQNW  >isoform b  MDMRTVSTGDMVPTISNSDSILLKIGNFSGGAKLQERILIIYKILQNERIICRRR |
| 42 | MRVQVVTLLAVLLAVLQFTSAAGNYYSGYPSDRTMKRSALLQPENNPEWNQLGWAWGKRSAGMEIPHRAARALHPVKKNPDWQDLGFAWGRK |
| 43 | MSLAQSTFYLLFVAFLAVVIAVTADKQQSAYDLETPISAYKRFYSWEDAKRAASSEEGMRNKRKQFYAWAGKRSSAPVHYFEDSIAQEAGPSMEKRKQFYAWAGK |
| 44 | MLLWIVATLLIFSLPVSTALDYNDFSLQRIARAPHPSSALLVPYPRVGKRSNILNNNSESQNSVQKRLYMARVGKRAFFYTPRIGK |
| 45 | MRAVTILSLLGAIIVVNASYRATESPEQWRQRFLKSQSEDPSWLRFPTPPSRLPPLYVETEKQIRDRALAYREKYEKEHPRPTVAADSDIDVEKRRNLLVGRYGFRIGKRFVGDQNDSPSSSAVQMLPVQVMYDDVLQ |
| 46 | MLSIRTFVLVLLVLVGLAAALPYFRLNYDYDPVDLDNDEKNAARQFLPYMKRNIAIGRGDGLRPGK |
| 47 | MQLYVVLCFLVLLGLSAGQMTFTDQWTKKRATLHKQLPVVTPEEPICPSDRVQAVFEQLDQLQKAQQRLTEYLASCAYPVEVPQKAEKM |
| 48 | MMSSRRIVYTLLLITIALCAYSTAYAPRNIRGVGDVPSMFFSPFRMMGKRNQFYGLFKKIRSNEVVDY |
| 49 | MMKWLLLAVFCIAAYAWADGTEDSNIDQLMSRVYRTVLLKSKRSPSMGLSLAEYMASPQGGDNFHFMPSGRK |
| 50 | MRFSVFVALFALLAVSYGFMYEMEDSYPVSGELPKRSLENFWRNIHLQMPSGRGGQKRTEGLSRASANAYYRLG |
| 51 | MRFLILALLVLFAITQAYPSSDYQPRYRKSQTQEANIQPFIRFRKSTQPQHNWMFRPDVAPYFE |
| 52 | MFRTLLICFFLMAAVSITSAYILDYDVPERQTRGDVKSVFFSPFRMVGKRSQFYGLYKQLGHSKRETSLE |
| 53 | MSSWLRTLFVFFIVLVVSTSALPSDYVRFLIQSARNHENSYYPQEEAGFMRVNRNQGAGSVSLDSLASLPMLRYG |
| 54 | MMNYSVVLLLFALCSIVAASPMWFDEEEPMNLRAFRVMPNQLDSIDASRRLLKRASMFNKRRGRELFGKRSLPIEYPDETVMYESRFRRRANELFG |
| 55 | MSCISMLFILLVACLLVSNAMYINPDYYYVEQLPTMKKSGQLRALAGSRNCFFSPVNCIITHDINSYRRLAKGSSYA |
| 56 | > isoform a  MPSPSSLLGSLLLVCAVLTITSRASSIMTDDVEPPQLLTRQLRSFPYSVSFYRMLGHDRQLRPYYGVNDEVAALIDSMNSDNVANEDVFPTRPRRSDGLRGSFYWARIRLPLQIL  > isoform b  MPSPSSLLGSLLLVCAVLTITSRASSIMTDDVEPPQLLTRQLRSFPYSVSFYRMLGHDRQLRPYYGVNDEVAALIDSMNSDNVANEDVFPTRPRRSDGLRGYACRFKFCRIYDA  > isoform c  MPSPSSLLGSLLLVCAVLTITSRASSIMTDDVEPPQLLTRQLRSFPYSVSFYRMLGHDRQLRPYYGVNDEVAALIDSMNSDNVANEDVFPTRPRRSDGLRGSFYWARFVVCQPVLVI |
| 57 | MRFILGLLIAIVAFVASSPIHGIWNNLPAPPQKRVYGFYNYLPKEEDDRDKRNTILLLTPNEDYVE |
| 58 | MISKCSVMGLLLLLFVHITTAQFETHFDDAQVYWPSQYKRSGPSSASEGEAYAFPGLRGLRGKRDPTYHKRVPMMSLKGLRGKRARIFDGQEEQ |
| 59 | MSALHIARLFLLIAIMSTVVSTYAVLPSGEILFRRPASLFDRAADYSDDVAKFDESRDKRELDGIDTQLSKRTNLKRLVILSARGFGKK |
| 60 | MSSSSSTFISIFIITMVLVAFCSANLATGRASLRPSAGKRAAVVGRLPAHYYQLASAVRSPKQYENEFTCNENTMSVINARYQKLQEELNGLVELMEACQTLQSTISL |
| 61 | MSHWIATLLAFILIANAFAALSTGADVGASDDVRSYVMPFYGMEKKWSRREPSIRFFKRNGGGQDLPPSRFLWDY |
| 62 | MVKFFVFLLFFAFLCSFTSAIPLRSLFLRSYDDINQESIARGYFAPQPVDDDNADNRPKRGIDLLKRRVEIIERNRCFFNPITCY |
| 63 | MLVSLLICLALVAVASAMPFYASSNDGDVVRQFPQQGVLENMVNEIMRVQSVEQMNNHHVRRQIFSRKFW |
| 64 | MMAQKTLIIAVMLVCSILQPMLALGSLTPSAAFRANMQQRERSPNTLFYMDGASKQYGDEIKDPIYKRFKPCYYSPIQCLIKRK |
| 65 | MRSWIPLLVLFAVLAVFAQAGKSSESDESRRRPSKSSESSDSDSKSSDSDSSSSEESSGDVPSEAPNTDSTPVEILAAAKPDSGILLGPGDNRVKRDGLPSFYDIRKKRGLPSAYDIRRK |
| 66 | MSSGKLFQFFIVFLATLLLADAIPMVSSRDEDDQIIQKRLSNDALIRLLMRNRGTQTQLGLKRGLVKKAEVERRSIDEDFSNCFLSPVQCMLPSSRK |
| 67 | MRVLTFLLVTLFALANVMQAQRYDRAIYEALLNDLEREFVERELAQHVLEKRELLRQDRQELDRVRRASEKKSYPRNCYFSPIQCLFTRN |
| 68 | MLLVLLFSLFSVGFGMRVMRRELLKDQPTVLLLGPLEVLTSSSSEDDTPDFPSLRDKRGVDPMSIPRLIKEPPLKKRSGDFKRGDVVYPSAAKQTVPLVRVREPPLKRGQMFLEELLQFNDDAHRQFMDPLRRIRYGPNRLIYTW |
| 69 | MHFFPILLLSILLILISTCSSTLVNNSPTAAFDTNAYSELNAKAKASMRRLADILDFEMYQRRLSAAPDNSYYYQISPPHHQRISSLPLTIFPLPDKRLHRIGGNIVMGK |
| 70 | MSVSRPLNLAILSILVGLLYLSACTCAPSVSAHHLGLRLKKWYEWNNDMEITKKWYDWQNVPHALQQKRQPFETSDYIIE |
| 71 | MKSSSVLSVALIVLVIVQLISASLASVPSSSAVSDGQIDFDALAAKIEMLRPNRYWKRAHNIDTRALNQFKNCYFSPIQCVLMERRRK |
| 72 | MLTRVPVLILAVIVMLALCQEPEKPEKRPALLSRYGRAVLPRYGKRSGNLMESSQNSLTEESSDVVCQLIDGKYICLPVDAVRFRPFFL |
| 73 | MSCSSSSMLFLVLIATTVLIAESRVFYNRFDGGLSSDRFMEQKRDGAEASYDYDANQVIRNTMKRNRQCLLNAGLSQGCDFSDLLHAQTQARKFMSFAGPGK |
| 74 | MNRFIISMIALLAVFCAVSTASPLLYRAPQYQMYDDVQFVKRSNAELINGLIGMDLGKLSAVGKRSNAELINGLLSMNLNKLSGAGRR |
| 75 | MGSSPILLVLAISIGLASACFLNSCPYRRYGRTIRCSSCGIENEGVCISEGRCCTNEECFMSTECSYSAVCPELFCKIGHHPGYCMKKGYCCTQGGCQTSAMC |
| 76 | MSRKFLLVFIAICVLSQNAMALRGALFRSGRSLIHRQSQNEEAVLNQPHLLPAEKRSPIVELYPVVDSGNVEPEAFPAYFRF |
| 77 | MCRLAVFALLAVAAVSVYGQPAGGQDVPPFLRNATPAQLQSFQQLIQANGHLTETALDGKVQAWVNQQGGKVAADWADFQKFIKGQQGQAEAAHQAAVSNFSPAAKKADADLTAISNDSSLSVQAKGQKIQAYLNSLPANVKAELEKAQGQ |
| 78 | MSRFLVIFAIFLAFSNICSAYPDYRLPERRIRGDVDSVFFSPFRIIGKRALLAGPHDYDLGDFISNPNV |
| 79 | MSTRWFVFVALMALVLLSAHTTEAMPRPGQMSLSNTRNCFFSPMGCVFVPKMSRIRKLSADFRNEIPPPDYI |
| 80 | MAVINRALLLLCILFALSEAYSRMELEDRMQMSRFQEPVKGAAGQMGGDPYIHYLSEYFGRPMKRHSAGSTYPESL |
| 81 | MLKSSIFSLLIVLLMVCSFGNVSANDFFLRSAKWVSKMNPSGGALVSGRGGFRPGFVSRDWRHAMAEPNFVKRSYNDY |
| 82 | MPSYHTVIIILLISIISTTSFILQDLPLERFERSGHGSDELLGGGSQFDRHIRSGLSYRPMPNLMARYRMMSG |
| 83 | MARFTPLLMILLALVPLYYSLDCRKFSFAPACRGIMLKRSGGHPMIAEQQPMIDNAKAREVQMMELLIRNIEDEALIANSDCVSMSWLRDRLVNAKNEMPQ |
| 84 | MFGHSTSFCLLALFVLSMTVNCKDPWYDWKVGKGLNGRSSFMGYQFDSPLERNAIKGYRYLSTRGRVTDCNSNCRYYYNANSGYCAPSRSNDCSTYCGCGFVCRCR |
