## Supplementary Table 5 for "*In silico* analyses of neuropeptide-like protein (NLP) profiles in parasitic nematodes"

**Table S5: Parasite *nlp-*gene family classifications based on sequence or functional similarity to known invertebrate neuropeptide families, or conservation of nematode peptide motif.** X denotes a variable amino acid**.** All peptides with a C-terminal glycine are predicted to be amidated. Note that for *nlp-36* and *nlp-77*, the portion of the prepropeptide that is mostly highly conserved in parasite species does not appear to represent the predicted peptide regions, but rather a predicted transmembrane region and Domain of unknown function (Prion-like-(Q/N-rich)-domain-bearing protein/DUF 148) respectively (see Figure S1). **^a^**Similar peptide conserved in atleast one nematode parasite species examined here.

| Family designation | *Nlp*-gene | *C. elegans* predicted peptides (Van Bael et al., 2018) | Similar peptide encoded in parasite sequelogs | Novel conserved *C. elegans* peptides predicted here^a^ | Characteristic peptide motif | Reference |
| --- | --- | --- | --- | --- | --- | --- |
| Buccalin/sulfakinin-like | *nlp-1* | **MDANAFRMSFG**  **MDPNAFRMSFG**  **VNLDPNSFRMSFG**  **STVGYNLDARNYFVGLG** | **Yes**  **Yes**  **Yes**  **Yes** |  | **M(S/G)FG** | **Nathoo et al. (2001)** |
|  | *nlp-7* | **LYLKQADFDDPRMFTSSFG**  **QADFDDPRMFTSSFG**  **SMDDLDDPRLMTMSFG**  **MILPSLADLHRYTMYD**  **GSDIDDPRFFSGAFG** | **Yes**  **Yes**  **Yes**  **No**  **Yes** |  | **FDDPR(M/F/L/Y)F(S/T)(S/A)FG** | **Nathoo et al. (2001)** |
|  | *nlp-13* | **NDFSRDIMSFG**  **SGNTADLYDRRIMAFG**  **QPSYDRDIMSFG**  **SAPSDFSRDIMSFG**  **SSSMYDRDIMSFG**  **SPVDYDRPIMAFG**  **AEDYERQIMAFG** | **Yes**  **Yes**  **Yes**  **Yes**  **Yes**  **Yes**  **Yes** |  | **M(H/S)FG** | **Nathoo et al. (2001)** |
| RP-amide | *nlp-2* | **SIALGRSGFRPG**  **SMAMGRLGLRPG**  **SMAYGRQGFRPG** | **Yes**  **No**  **Yes** |  | **SXA(L/Y)(G/S)RX(G/N)FRPG** | **Nathoo et al. (2001)** |
|  | *nlp-22* | **SIAIGRAGFRPG** | **No** |  |  |  |
|  | *nlp-23* | **LYISRQGFRPA**  **SMAIGRAGMRPG**  **AFAAGWNRG** | **No**  **No**  **No** |  |  |  |
|  | *nlp-46* | **NIAIGRGDGLRPG** | **Yes** |  | **NIAIGR(G/A)DG(F/L)RPG** | **McVeigh et al. (2008)** |
|  | *nlp-85* | **n/a** | **n/a** | **n/a** | **A(A/V/T)MISGRGFRPG** |  |
|  | *nlp-86* | **n/a** | **n/a** | **n/a** | **GRW(G/Q)LRPG** |  |
|  | *nlp-87* | **n/a** | **n/a** | **n/a** | **S(I/L)ALGR(F/L)(S/N)LRPG** |  |
| Orkokinin-like | *nlp-3* | **AINPFLDSMG**  **AVNPFLDSIG**  **YFDSLAGQSLG**  **SFRPDMITEE**  **SNNRYEMLENYY** | **Yes**  **Yes**  **Yes**  **No**  **No** |  | **RYFD(S/A)LAGQ(S/A)LG** | **Nathoo et al. (2001)** |
|  | *nlp-8* | **AFDRFDNSGVFSFGA**  **AFDRMDNSDFFGA**  **SFDRMGGTEFGLM**  **YPYLIFPASPSSGDSRRLV**  **PYLIFPASPSSGDSRRLV**  **PYLIFPASPSSGDS**  **YPYLIFPASPSSGDS**  **FDRYDDETAYGYGFDNHIF**  **SADPYRFMSVPT** | **Yes**  **Yes**  **Yes**  **No**  **No**  **No**  **No**  **No**  **No** | **SAPESREQLINNLAESIITL** | **AFDR** | **Nathoo et al. (2001)** |
|  | *nlp-14* | **ALDGLDGSGFGFD**  **ALNSLDGAGFGFE**  **ALDGLDGAGFGFD**  **ALNSLDGQGFGFE**  **ALNSLDGNGFGFD** | **Yes**  **Yes**  **Yes**  **Yes**  **Yes** |  | **AL(N/D)XL(E/D)G** | **Nathoo et al. (2001); Husson et al. (2005)** |
|  | *nlp-15* | **AFDSLAGSGFDNGFN**  **AFDSLAGSGFGAFN**  **AFDSLAGSGFSGFD**  **AFDSLAGQGFTGFE**  **AFDTVSTSGFDDFKL** | **Yes**  **Yes**  **Yes**  **Yes**  **No** |  | **(A/S/P)FDX(L/F)** | **Nathoo et al. (2001); Husson et al. (2005)** |
| Allatostatin-A-like | *nlp-5* | **SVSQLNQYAGFDTLGGMGLG**  **ALSTFDSLGGMGLG**  **ALQHFSSLDTLGGMGFG** | **Yes**  **Yes**  **No** |  | **D(P/T)L(G/A)G(M/I)G/A)LG** | **Husson et al. (2009)** |
| Allatostatin-A-like/buccalin-like | *nlp-6* | **MAAPKQMVFGFG**  **APKQMVFGFG**  **YKPRSFAMGFG**  **AAMRSFNMGFG**  **LIMGLG**  **SMADISDMPEDVPM**  **SAEPQIDVDSF** | **Yes**  **Yes**  **Yes**  **Yes**  **Yes**  **No**  **No** |  | **(M/L/I)G(F/L)G** | **Husson et al. (2009); Mirabeau and Joly (2013)** |
| SIFamide-like | *nlp-10* | **AIPFNGGMYG**  **STMPFSGGMYG**  **AAIPFSGGMYG**  **GAMPFSGGMYG**  **SLVPQSYSNNENQI** | **Yes**  **Yes**  **Yes**  **Yes**  **No** |  | **GG(M/L/V/I)YG** | **Elphick (2013)** |
| Sulfakinin/Cholecystokinin/gastrin-like | *nlp-12* | **DYRPLQFG**  **DGYRPLQFG**  **TQSPTFDRQD**  **SSGSSGPVVLEPIWEWQ** | **Yes**  **Yes**  **Yes**  **No** |  | **RPL(Q/M)FG** | **Janssen et al. (2008)** |
| Pigment dispersing factor | *nlp-37* | **NNAEVVNHILKNFGALDRLGDVG** | **Yes** |  | **NNAE(V/L)VNHILKN(F/L)X_2_(L/V)D(R/L)LG(E/D)VG** | **Janssen et al. (2009)** |
|  | *nlp-74* | **SNAELINGLIGMDLGKLSAVG**  **SNAELINGLLSMNLNKLSGAG**  **SPLLYRAPQMYDDVQFV**  **SPLLYRAPQYQMYDDVQFV** | **Yes**  **Yes**  **No**  **No** |  | **NAEL(I/V)NGL(I/L)GM** | **Janssen et al. (2009)** |
| Allatostatin-B-like, Mioinhibiting peptide like | *nlp-38* | **VLGWNKAHGLWG**  **TPQNWNKLNSLWG**  **SPAQWQRANGLWG**  **ASDDRVLGWNKAHGLWG**  **SNLSPRFLG**  **SNLSPRFLGRMTFARIPKISPAQWQRANGLWG** | **Yes**  **Yes**  **Yes**  **Yes**  **No**  **Yes** |  | **GLWG** | **Husson et al. (2005)** |
| Allatostatin-B-like, Mioinhibiting peptide like | *nlp-42* | **SALLQPENNPEWNQLGWAWG**  **NPDWQDLGFAWG**  **AGNYYSGYPSDRTM** | **Yes**  **Yes**  **No** |  | **LGWXWG** | **Husson et al. (2007)** |
| Leucokinin-like | *nlp-43* | **KQFYAWAG**  **DKQQSAYDLETPISAY**  **SSAPVHYFEDSIAQEAGPSME** | **Yes**  **No**  **No** |  | **FYAWAG** | **Husson et al. (2007)** |
| Cardioactive peptide/ neuromedin-U/  pyrokinin/periviscerokinin-like | *nlp-44* | **APHPSSALLVPYPRVG**  **LYMARVG**  **AFFYTPRIG** | **Yes**  **Yes**  **Yes** |  | **(A/P)RVG** | **Lindemans et al. (2009a)** |
| Adipokinetic hormone/Gonadotropin releasing hormone-like | *nlp-47* | **QMTFTDQWT** | **Yes** |  | **Q(M/L/F)TF(S/T)XW** | **Husson et al. (2009); (Lindemans et al., 2009b)** |
| Thyrotropin releasing hormone | *nlp-54* | **RGRELFG**  **GRELFG**  **RANELFG**  **ANELFG** | **Yes**  **Yes**  **Yes**  **Yes** |  | **ELFG** | **Van Sinay et al. (2017)** |
| Allatostatin-C-like | *nlp-55* | **MYINPDYYYVEQLPTM**  **SGQLRALAGSRNCFFSPVNCIITHDINSY**  **LAKGSSYA**  **GSSYA** | **No**  **Yes**  **No**  **No** |  | **RNC(F/Y)F(S/T)PX_2_C** |  |
|  | *nlp-62* | **CFFNPITCY**  **RVEIIERNRCFFNPITCY** | **Yes**  **Yes** |  | **RCFFNPI(T/S)C** | **Mirabeau and Joly (2013)** |
|  | *nlp-64* | **PCYYSPIQCLI**  **FKPCYYSPIQCLI** | **Yes**  **Yes** |  | **PC(F/Y/M)(Y/F)(S/N)PIQC(L/I)X** | **Mirabeau and Joly (2013)** |
|  | *nlp-66* | **SIDEDFSNCFLSPVQCMLPSS** | **Yes** | **LSNDALIRLLM** | **DXDF(S/Q)NCFLSPVQCMLP** | **Mirabeau and Joly (2013)** |
|  | *nlp-67* | **NCYFSPIQCLFT**  **SYPRNCYFSPIQCLFT** | **Yes**  **Yes** | **AIYEALLNDLEREFVERELAQHVLE** | **SYPR(N/I)CYFSPIQCLFTR** | **Mirabeau and Joly (2013)** |
|  | *nlp-71* | **NCYFSPIQCVLMERR**  **AHNIDTRALNQFKNCYFSPIQCVLMERR** | **Yes**  **Yes** |  | **FKNCYFSP(I/V)QCVL** | **Mirabeau and Joly (2013)** |
|  | *nlp-79* | **LSADFRNEIPPPDYI** | **No** | **NCFFSPMGCVFVP** | **RNCFFSP(M/I)(G/N)C(M/V)F** |  |
|  | *nlp-88* |  |  | **YDRNCFFSPVQCMLTYNDDS**  **LPLVVTRKSS** | **(Y/F)DRNCFFSPVQCML** |  |
|  | *nlp-89* |  |  | **NCFFSPVQCMFDTSSADALDSF** |  |  |
|  | *nlp-90* |  |  | **NCFFSPAQCILENFP** |  |  |
|  | *nlp-91* |  |  | **NCFFTPVQCMLPMTDSH** |  |  |
|  | *nlp-92* | **n/a** | **n/a** | **n/a** | **LR(N/K)CF(F/L)SPVQCLLPF** |  |
|  | *nlp-93* | **n/a** | **n/a** | **n/a** | **RNC(F/Y)(F/L)SP(V/I/M)QCX_2_P** |  |
|  | *nlp-94* | **n/a** | **n/a** | **n/a** | **F(R/K)RNCFFSPVQCMF** |  |
|  | *nlp-95* | **n/a** | **n/a** | **n/a** | **KNCF(F/L/M)S(P/L)(V/L)QCSF(Y/F)Y** |  |
|  | *nlp-96* | **n/a** | **n/a** | **n/a** | **SRX(P/L)(S/K)(L/F)DRNCFFSPVQCM(L/F)** |  |
| Tachykinin-like | *nlp-58* | **VPMMSLKGLRG**  **ARIFDGQEEQ** | **Yes**  **No** | **SGPSSASEGEAYAFPGLRGLRG** | **G(L/V/S)RG** | **Mirabeau and Joly (2013)** |
| Allatotropin-like | *nlp-59* | **TNLKRLVILSARGFG**  **LVILSARGFG** | **Yes**  **Yes** |  | **ARGFG** | **Koziol et al. (2016)** |
| Luqin-like | *nlp-72* | **PALLSRYG**  **AVLPRYG** | **Yes**  **Yes** |  | **A(V/A/L)L(S/P)RYG** | **Mirabeau and Joly (2013)** |
| Calcitonin-like | *nlp-73* | **FMSFAGPG**  **NRQCLLNAGLSQGCDFSDLLHAQTQARKFMSFAGPG** | **Yes**  **Yes** |  | **(N/S)RXC(L/I)(L/I)NAGLSQGCD(L/F)-**  **SDXLXA(K/Q)XQAXKFXSFAGPG** | **Mirabeau and Joly (2013)** |
| Oxytocin/vasopressin related (nematocin) | *nlp-75* | **CFLNSCPYRRYG** |  |  | **(C/F)F(L/I)N(S/Q)CPYR(R/H)(Y/I)G** | **Beets et al. (2012)** |
| L-11 peptide-like | *nlp-83* | **FSFAPACRGIML** | **Yes** |  | **(F/Y)(S/P)(F/Y)AP(A/P/K)CRGI(M/I/V)L** | **Yamanda et al. (2010)** |
|  | *nlp-4.1* | **None predicted to date** | **No** |  |  |  |
|  | *nlp-4.2* | **SLILFVILLVAFAAARPVSEEVDRV**  **RPVSEEVDRV**  **PVSEEVDRV**  **DYDPRTEAPRRLPADDDEVDGEDRV**  **LPADDDEVDGEDRV**  **DYDPRTDAPIRVPVDPEAEGEDRV** | **No**  **No**  **No**  **No**  **No**  **No** |  |  |  |
|  | *nlp-9* | **GGARAFYGFYNAGNS**  **GGGRAFNHNANLFRFD**  **GGGRAFAGSWSPYLE**  **GGGRAFAGSWSPYLERFYDY**  **TPIAEAQGAPEDVDDRRELE**  **TPIAEAQGAPEDVDD**  **DQAAALPYYLYE** | **Yes**  **Yes**  **Yes**  **Yes**  **No**  **No**  **Yes** |  | **GGARAF** | **Nathoo et al. (2001)** |
|  | *nlp-16* | **STEHHRV**  **SEGHPHE**  **ATHSPEGHIVAKDDHHGHE**  **SSDSHHGHQ**  **SVDEHHGHQ**  **NAEDHHEHQ**  **SEHVEHQAEMHEHQ**  **STQEVSGHPEHHLV**  **GLYSSERTEEEVEISHGMHHRE**  **HSEHLPHPDPPSHTA** | **No**  **No**  **No**  **No**  **No**  **No**  **No**  **No**  **No**  **No** |  |  |  |
|  | *nlp-21* | **GGARAMLH**  **GGARAFSADVGDDY**  **GGARAFYDE**  **GGARAFLTEM**  **GGARVFQGFEDE**  **GGARAFMMD**  **GGGRAFGDMM**  **GGARAFVENS**  **GGGRSFPVKPGRLDD**  **QYTSELEEDE** | **Yes**  **Yes**  **Yes**  **Yes**  **Yes**  **Yes**  **Yes**  **Yes**  **Yes**  **No** |  | **GGARAF** | **Nathoo et al. (2001)** |
|  | *nlp-11* | **HISPSYDVEIDAGNMRNLLDIG**  **SAPMASDYGNQFQMYNRLIDAG**  **SPAISPAYQFENAFGLSEALERAG** | **Yes**  **Yes**  **Yes** |  | **LXDXG** | **Nathoo et al. (2001)** |
|  | *nlp-17* | **GSLSNMMRIG**  **QQEYVQFPNEGVVPCESCNLGTLMRIG**  **QMSRQQEYVQFPNEGVVPCESCNLGTLMRIG**  **ANFENDQDVMRPPFQAL** | **Yes**  **Yes**  **Yes**  **No** |  | **MRXG** | **Nathoo et al. (2001)** |
|  | *nlp-18* | **SPYRAFAFA**  **ARYGFA**  **SPYRTFAFA**  **ASPYGFAFA**  **SDEENLDFLE**  **GQFSSFA** | **Yes**  **Yes**  **Yes**  **Yes**  **Yes**  **Yes** |  | **FAFA** | **Nathoo et al. (2001)** |
|  | *nlp-20* | **FAFAFA**  **SGPQAHEGAGMRFAFA**  **APKEFARFARASFA**  **ASQYSDDDSELMSNNERFA** | **Yes**  **Yes**  **Yes**  **Yes** |  | **FAFA** | **Nathoo et al. (2001)** |
|  | *nlp-19* | **RIGRLPNFL**  **IGLRLPNFL**  **IGLRLPNFLRF**  **IGLRLPNML**  **IGLRLPNMLKF**  **MGMRLPNIIFL**  **MGMRLPNIIFLRNE** | **Yes**  **Yes**  **Yes**  **Yes**  **Yes**  **Yes**  **Yes** |  | **(I/F)(G/V)(L/I/M)X(L/M/V)P(N/G)** |  |
|  | *nlp-35* | **AVVSGYDNIYQVLAPRF**  **ARMMSFDAEAPQYLQHLLQNLKPRF** | **Yes**  **Yes** |  | **LX(H/Q)L(L/M)Q(N/R)LKP(R/K)F** |  |
|  | *nlp-36* | **DDDVTALERWGY**  **NIDMKLGPH**  **SMVARQIPQTVVADH** | **Yes**  **Yes**  **No** |  | **D(D/E)XT** |  |
|  | *nlp-39* | **EVPNFQADNVPEAGGRV**  **EVPNFQADNVPEAGG**  **LDSPIPEFYSTGRSTRVPSHRHRIPRLG** | **No**  **No**  **No** |  |  |  |
|  | *nlp-40* | **APSAPAGLEEKL**  **MVAWQPM**  **VAWQPM**  **EQPAAADTFLGFVPQ**  **APSAPAGLEEKLR** | **No**  **Yes**  **Yes**  **No**  **No** | **ALQEQLYSLE**  **APLLHAIEA**  **LAEVLRAGERLGVNPEEVLADL**  **APLLHAIEARLAEVLRAGERLGVNPEEVLADL**  **APLLHAIEARLAEVLRAGERLGVNPEEVLADLRARNQFQ** | **AWQPM** |  |
|  | *nlp-41* | **APGLFELPSRSV**  **APGLFELPSRSVRLI** | **No**  **No** |  |  |  |
|  | *nlp-45* | **RNLLVGRYGFRIG** | **No** |  |  |  |
|  | *nlp-48* | **GVGDVPSMFFSPFRMMG** | **Yes** | **NQFYGLF** | **F(N/F)(S/N)PFX_3_G, NQFYGLF** |  |
|  | *nlp-49* | **WADGTEDSNIDQLMSRVYRTVLLKS**  **SPSMGLSLAEYMASPQGGDNFHFMPSG**  **DGTEDSNIDQLMSRVY** | **Yes**  **Yes**  **Yes** |  | **S(M/I)GLSLAEY(M/V)AX_6_FHF** | **Chew et al. (2018)** |
|  | *nlp-52* | **GDVKSVFFSPFRMVG**  **YILDYDVPERQT** | **Yes**  **No** | **SQFYGLY** | **F(N/F)(S/N)PFX_3_G, SQ(F/L)YG(Y/L)Y** |  |
|  | *nlp-50* | **SLENFWRNIHLQMPSG**  **TEGLSRASANAYYRLG**  **FMYEMEDSYPVSGELP** | **Yes**  **Yes**  **Yes** |  | **TE(G/S)LSRA(S/R)ANAYYRLG** |  |
|  | *nlp-51* | **SQTQEANIQPFIRF**  **YPSSDYQPRY** | **Yes**  **Yes** |  | **(I/V)QPFIRF** |  |
|  | *nlp-53* | **NQGAGSVSLDSLASLPMLRYG** | **No** |  |  |  |
|  | *nlp-56* | **SSIMTDDVEPPQLLTRQL** | **No** | **GYACRFKFCRIYDA** | **YXCRFKFCRI(F/Y)** |  |
|  | *nlp-57* | **SPIHGIWNNLPAPPQ**  **VYGFYNYLPKEEDDRD**  **NTILLLTPNEDYVE** | **No**  **Yes**  **No** |  | **YGXYX_2_(L/I)P** |  |
|  | *nlp-60* | **NLATGRASLRPSAG** | **Yes** |  | **NLX(7)RPS(T/A)G** |  |
|  | *nlp-61* | **WSRREPSI** | **Yes** | **WSRREPSIRFF** | **W(S/T)X_2_EPS(I/V)** |  |
|  | *nlp-63* | **QIFSRKFW** | **Yes** |  | **Q(I/M)FSRKFW** |  |
|  | *nlp-65* | **DGLPSFYDIR**  **DGLPSFYDI**  **GLPSAYDIR**  **GLPSAYDI** | **No**  **No**  **No**  **No** |  |  |  |
|  | *nlp-68* | **GVDPMSIPRLIKEPPL**  **GDVVYPSAAKQTVPLVRVREPPL** | **Yes**  **Yes** |  | **EPPL** |  |
|  | *nlp-69* | **LHRIGGNIVMG** | **Yes** |  | **RIGGXI(V/M)MG** |  |
|  | *nlp-70* | **WYEWNNDMEIT**  **WYDWQNVPHALQQ** | **Yes**  **Yes** |  | **WYXW** |  |
|  | *nlp-76* | **SPIVELYPVVDSGNVEPEAFPAYFRF**  **GALFRSG** | **No**  **Yes** |  | **GAL(F/Y)R(S/T)** |  |
|  | *nlp-77* | **QPAGGQDVPPFL** | **Arguably** |  | **PPFL** |  |
|  | *nlp-78* | **GDVDSVFFSPFRIIG**  **ALLAGPHDYDLGDFISNPNV** | **No**  **No** |  |  |  |
|  | *nlp-80* | **HSAGSTYPESL** | **No** |  |  |  |
|  | *nlp-81* | **NDFFLRSA** | **Yes** | **WVSKMNPSGGALVSGRGGFRPGFVS** | **LVSGRG(G/N)FRPGF** |  |
|  | *nlp-82* | **FILQDLPLE**  **FILQDLPLERFE** | **Yes**  **Yes** |  | **F(I/V)LQDLPLER(F/Y)ER** |  |
|  | *nlp-84*  */rgba-1* | **KVGKGLNG** |  |  |  |  |
|  | *nlp-97* | **n/a** | **n/a** | **n/a** | **NPYSW** |  |
|  | *nlp-98* | **n/a** | **n/a** | **n/a** | **S(L/V)AP(S/T)TS** |  |
