## Supplementary Table 6 for "*In silico* analyses of neuropeptide-like protein (NLP) profiles in parasitic nematodes"

**Table S5:** **The effects of *As*-NLP peptides on the ovijector of *A. suum****.* Contraction frequency and amplitude recorded two minutes prior to each timepoint, presented as percentage of time 0. Change in tension presented relative to time 0, where muscle relaxation caused an increase in circular muscle tension (+mg/mm; shortening of the tissue) whereas circular muscle contraction caused a decrease in tension (-mg/mm; lengthening of tissue). P.a. denotes post addition, overall indicates the significance of the Repeated Measures ANOVA test comparing pre- and post-addition, NS indicates results not statistically significant, P denotes P value (significant when P< 0.05), F denotes F ratio. ↑ indicates increase in contraction frequency/amplitude, ↓ indicates decrease in contraction frequency/amplitude. 🗸 indicates significant Dunnet’s post test, 🗴 indicates non-significant Dunnet’s post test.

| **Peptide tested**  [10µM] | **Time**  (min p.a.) | **Contraction frequency**  (% 0 min, 2 min  prior to min p.a.) | **Contraction amplitude**  (% 0 min, 2 min  prior to min p.a.) | **Change in tension**  (mg/mm) |
| --- | --- | --- | --- | --- |
| **NLP-12A1** (DLTPTRFDRQDRDYRPLQF-NH_2_)  n≥3 | N/A | No effect | No effect | No effect |
| **NLP-12B** (DGYRPLQF-NH_2_)  n≥3 | N/A | No effect | No effect | No effect |
| **NLP-12C** (SYRPLQF-NH_2_)  n≥3 | N/A | No effect | No effect | No effect |
| **NLP-12D** (AYDFSEYTPY)  n≥3 | N/A | No effect | No effect | No effect |
| **NLP-36A1** (DDRTVFEKTE)  n≥3 | N/A | No effect | No effect | No effect |
| **NLP-36A2** (DDRTVFE)  n≥3 | N/A | No effect | No effect | No effect |
| **NLP-36B1** (TERTHTEAAWGL)  n=6 | 50secs  2  5  10  **overall** | N/A  ↓ 89.34±13.27 🗴  ↑ 119.40±7.37 🗴  ↑ 108.20±8.12 🗴  **P=0.0268, F=4.065** | N/A  ↓ 82.21±8.05 🗸  ↓ 72.95±5.94 🗸  ↓ 77.28±6.97 🗸  **P=0.0049, F=6.514** | Lengthening -0.046±0.026 🗴  Shortening +0.002±0.024 🗴  Shortening +0.018±0.029 🗴  Shortening +0.029±0.024 🗴  **NS** |
| **NLP-36B2** (THTEAAWGL)  n=6 | 50secs  2  5  10  **overall** | N/A  ↑ 147.50±45.15 🗴  ↑ 105.20±19.85 🗴  ↑ 137.90±24.66 🗴  **NS** | N/A  ↓ 77.31±5.14 🗴  ↓ 83.59±12.10 🗴  ↓ 71.44±5.47 🗸  **P=0.0379, F=3.623** | Shortening +0.011±0.008 🗴  Shortening +0.017±0.015 🗴  Shortening +0.031±0.008 🗸  Shortening +0.019±0.012 🗴  **NS** |
| **NLP-36C1** (HPDAGFLLDSSENF)  n≥3 | N/A | No effect | No effect | No effect |
| **NLP-36C2** (HPDAGFLLDSSENFRVIGFI)  n≥3 | N/A | No effect | No effect | No effect |
| **NLP-47** (PQITFTDQWT)  n=6 | 50secs  2  2.5  5  10  **overall** | N/A  ↑ 114.70±6.73 🗴  N/A  ↑ 131.40±11.58 🗸  ↑ 132.10±10.78 🗸  **P=0.0219, F=4.328** | N/A  ↑ 102.20±7.71 🗴  N/A  ↓ 94.73±8.59 🗴  ↓ 86.03±11.12 🗴  **NS** | Lengthening -0.043±0.034 🗴  Lengthening -0.032±0.023 🗴  Lengthening -0.065±0.051 🗴  Lengthening -0.010±0.019 🗴  Shortening +0.006±0.020 🗴  **NS** |
| **NLP-49** (ASCYSVSTFWLSHIFIAASMGLSLAEYMASPQGQDNFHFIPS-NH_2_)  n=6 | 2  5  10  **overall** | ↑ 122.60±29.90 🗴  ↑ 105.30±49.97 🗴  ↓ 93.02±36.82 🗴  **NS** | ↓ 79.30±7.09 🗴  ↓ 71.78±13.33 🗸  ↓ 72.81±14.98 🗸  **NS** | Shortening +0.067±0.015 🗸  Shortening +0.068±0.016 🗸  Shortening +0.068±0.013 🗸  **P=0.0008, F=7.308** |
| **PDF-1A** (SNAELINGLIGMDLNKLSAI-NH_2_)  n=6 | 2  5  10  **overall** | ↑ 110.50±6.74 🗴  ↑ 123.70±15.62 🗴  ↑ 136.80±16.30 🗸  **NS** | ↓ 88.52±15.94 🗴  ↓ 93.64±21.53 🗴  ↓ 85.72±21.16 🗴  **NS** | Shortening +0.025±0.019 🗴  Shortening +0.027±0.014 🗴  Shortening +0.031±0.012 🗴  **NS** |
| **PDF-1B** (SNAELINGLLGMNLNRLSSA-NH_2_)  n=6 | 2  5  10  **overall** | ↑ 158.50±36.26 🗴  ↑ 148.20±42.84 🗴  ↑ 149.00±48.25 🗴  **NS** | ↓ 60.61±7.33 🗸  ↓ 75.48±12.82 🗴  ↓ 80.97±18.36 🗴  **P=0.0320, F=3.836** | Shortening +0.005±0.005 🗴  Lengthening -0.019±0.025 🗴  Lengthening -0.018±0.026 🗴  **NS** |
| **PDF-2C** (NNAEVVNHILKNFGALDRLGDV-NH_2_)  n≥3 | N/A | No effect | No effect | No effect |
